# Persistent spectral theory-guided protein engineering

**DOI:** 10.1101/2022.12.18.520933

**Authors:** Yuchi Qiu, Guo-Wei Wei

## Abstract

While protein engineering, which iteratively optimizes protein fitness by screening the gigantic mutational space, is constrained by experimental capacity, various machine learning models have substantially expedited protein engineering. Three-dimensional protein structures promise further advantages, but their intricate geometric complexity hinders their applications in deep mutational screening. Persistent homology, an established algebraic topology tool for protein structural complexity reduction, fails to capture the homotopic shape evolution during the filtration of a given data. This work introduces a **T**opology-**o**ffered **p**rotein **Fit**ness (TopFit) framework to complement protein sequence and structure embeddings. Equipped with an ensemble regression strategy, TopFit integrates the persistent spectral theory, a new topological Laplacian, and two auxiliary sequence embeddings to capture mutation-induced topological invariant, shape evolution, and sequence disparity in the protein fitness landscape. The performance of TopFit is assessed by 34 benchmark datasets with 128,634 variants, involving a vast variety of protein structure acquisition modalities and training set size variations.

## 1 Introduction

Protein engineering aims to design or discover proteins with desirable functions, such as improving the phenotype of living organisms, enhancing enzyme catalysis, and boosting antibody efficacy [1]. Traditional protein engineering approaches, including directed evolution [2] and rational design [3], resort to experimentally screening an astronomically large mutational space, which is both expensive and time-consuming. As a result, only a small fraction of the mutational space can be explored even with high-throughput screenings. Recently, data-driven machine learning models have been developed to reduce the cost and expedite the process of protein engineering by establishing the protein-to-fitness map (i.e., fitness landscape) from sparsely sampled experimental data [4, 5].

The most successful machine learning models for the protein fitness landscape are based on protein sequences using self-supervised models. There are two major types of self-supervised deep protein language models, i.e., local evolutionary ones and global ones. The former infers mutational effects by utilizing homologs from the multiple sequence alignment (MSA) [6] to model the mutational distribution of a target protein, such as Potts models [7], variational autoencoders (VAEs) [8, 9], and MSA Transformer [10]. Global evolutionary models learn from large sequence databases such as UniProt [11] to infer natural selection paths. Natural language processing (NLP) models have been adopted for extracting protein sequence information. For examples, tasks assessing protein embeddings (TAPE) [12] constructed three models, including Transformer, dilated residual network (ResNet), and long short-term memory (LSTM) architectures. Bepler [13] and UniRep [14] are two LSTM-based models. Recently, large-scale protein Transformer models trained on hundreds of millions of sequences exhibit advanced performance in modeling protein properties, such as evolutionary scale modeling (ESM) [15] and ProtTrans [16] models. In addition, the hybrid fine-tune approaches combining both local and evolutionary data can further improve the models. For examples, eUniRep is a fine-tuned LSTM-based UniRep model by learning from local MSAs [17]. The ESM model can be fine-tuned using either training data from downstream tasks [15] or local MSAs [18]. The Transformer-based Tranception score mutations via not only a global auto-regressive inference, but also a local retrieval inference utilizing MSAs [19]. One usage of evolutionary models is to generate the latent space representation for building downstream fitness regression model [4]. The other is to generate a probability density to evaluate the likelihood that a given mutation occurs [20]. Such a probability likelihood, referred to as evolutionary score or zero-shot prediction, predicts the relative fitness of protein variants of interest in an unsupervised approach without requirement of supervised label acquisition. The recent integration of local and global evolutionary models [21] and the combination between evolutionary score and sequence embedding [20] discovered the complementary roles of various sequence modeling approaches in building accurate fitness predictors.

Alternatively, three-dimensional (3D) structure or geometry-based models have also played an important role in predicting protein fitness landscape [22]. They offer somewhat more comprehensive and explicit descriptions of the biophysical properties of a protein and its fitness. With in-depth physical chemical knowledge, biophysical models such as FoldX [23] and Rosetta [24] are widely used to analyze protein fitness from its structure. However, the intricate structural complexity of biomolecules hinders the application of the first principle, such as quantum mechanics, to excessively deep mutational screening and thus, the most successful methods rely on their effective reduction of biomolecular structural complexity.

Topology concerns the invariant properties of a geometric object under continuous deformations (e.g., filtration). Topological data analysis (TDA) offers an effective abstraction of geometric and structural complexity [25, 26]. Persistent homology, a main workhorse of TDA, was integrated with machine learning models to reduce the structural complexity in drug designs [27], and protein-protein interactions [28]. Filtration of a given point cloud may induce both homotopic shape evolution and changes in topological invariants, which allow multiscale analysis from TDA. However, persistent homology only captures changes of topological invariants and is insensitive to the homotopic shape evolution. The persistent spectral theory (PST), a multiscale topological Laplacian, was designed to overcome this drawback [29]. PST fully recovers the topological invariants in its harmonic spectra and captures the homotopic shape evolution of data during the multiscale analysis [29, 30]. Its advantages for drug design have been confirmed [31].

In this work, we introduce a PST-based topological machine learning model, called **T**opology- **o**ffered **p**rotein **Fit**ness (TopFit), to navigate the protein fitness landscape. We demonstrate that the machine learning models with the PST embedding show substantial improvement over those with sequence-based approaches. On the other hand, the PST-based models require high-quality structures, whereas sequence-based models can deal with more general cases without resorting to 3D structures. These two approaches naturally complement each other in machine learning-assisted protein engineering. To further extend the potential of the topology-based methods, we combine 18 regression models into an ensemble regressor to ensure robust and accurate predictions over a variety of protein engineering scenarios, ranging from small training sets, imperfect structures, and varying experimental modalities, noisy data, etc. TopFit integrates complementary structure-sequence embeddings and ensemble regressor to achieve better performance over existing methods on 34 deep mutational scanning (DMS) datasets.

## 2 Results

### 2.1 Overview of TopFit

TopFit infers fitness landscape, a function that maps proteins to their fitness, from a small number of labeled data. Our framework consists of two embedding modules, i.e., topological structure-based and residue sequence-based embeddings, and an ensemble regressor weighting many supervised learning models (Figure 1).

**Figure 1:**
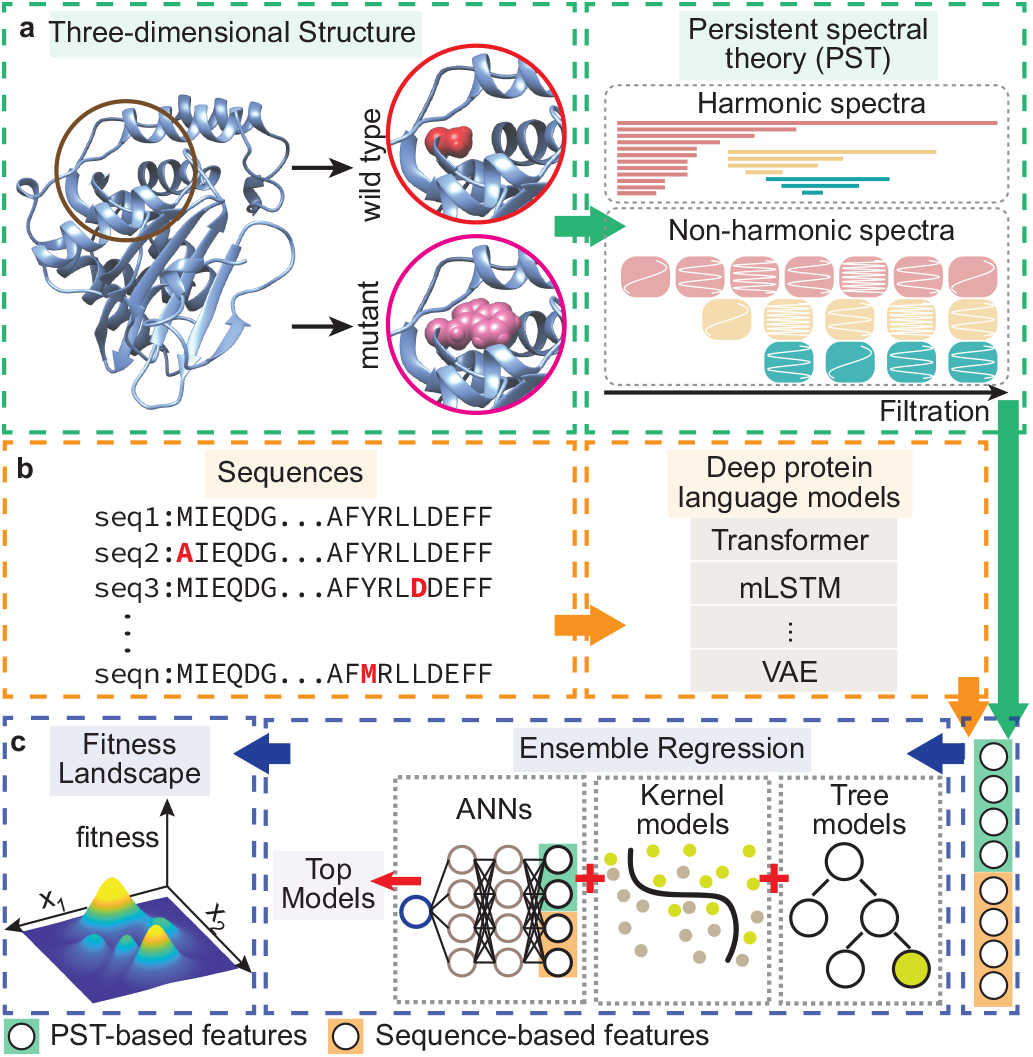
Conceptual diagram of the TopFit method. **(a).** Topological structure-based module generates PST embedding. A wild-type three-dimensional structure is obtained from the PDB or AlphaFold database. Both wild-type and mutant structures are optimized for local mutational-site structure extraction. PST embedding is generated from the spectra of persistent Laplacians where the harmonic and non-harmonic spectra delineate the mutational atlas. **(b).** Sequence-based embedding and evolutionary scores are generated by various deep protein language models. **(c).** Ensemble regression recruits multiple regressors in predicting fitness. Features from two featurization modules are concatenated and fed into the downstream ensemble regression. The ensemble regression ranks and selects top performing models from 18 regressors in three major classes: artificial neural networks (ANNs), kernel models, and tree models. Hyperparameters of each regressor are optimized. The ensemble regression averages the predictions from top regressors to navigate the fitness landscape.

In topological structure-based embedding, PST [29] is employed to generate protein structurebased embedding. A structure optimization protocol generates and optimizes both wildtype and mutational structures (Methods). PST assesses the mutation-induced local structure changes near the mutational sites (Figure 1**a**). Mathematically, PST constructs a family of topological Laplacians via filtration, which delineates interatomic interacting patterns at various characteristic scales. The topological persistence [30] and homotopic shape evolution of the local structure during filtration can be assessed from harmonic and non-harmonic spectra of the Laplacians (i.e., zero- and non-zero-eigenvalues), respectively (Figure 1**a**). The site- and element-specific strategies are further introduced in assisting PST to capture critical relevant biological information, such as hydrogen bonds, electrostatics, hydrophobicity, etc. (Methods). PST embeddings are generated on each pair of atoms for both local wild-type and mutant structures, and they are concatenated for downstream supervised models (Methods).

In sequence-based embedding, both protein sequences and evolutionary scores can be utilized (Figure 1**b**). We tested 9 sequence embedding methods and 5 evolutionary scores. Among them, the top-performing sequence embedding and evolutionary score are adopted in TopFit.

Features from two embedding modules are concatenated and fed into the downstream ensemble regression to predict the protein fitness landscape. In particular, we combined PST embedding, one sequence-based embedding, and one evolutionary score to build TopFit. In protein engineering, the number of experimental labeled data is typically small (i.e., 10^1^ ~ 10^2^) and accumulated at each round of screening. We utilize an ensemble regression model to provide a unified model with high accuracy and strong generalization for various sizes of training data. From a pool of multiple regressors, hyperparameters for individual regressors were first optimized by Bayesian optimization, and predictions from top *N* models were averaged (i.e., consensus) for fitness predictions [32]. Specifically, we employed 18 regressors of three major types: artificial neural networks (ANNs), kernel models and tree models (Figure 1**c**, Methods, and Supplementary Table 1). Since the evolutionary scores contain relatively higher-level features than embeddings do, we treated them differently in ANNs and ridge regression to enhance their weights over other features (Methods). When multiple structures (e.g., nuclear magnetic resonance (NMR) structure) are available for the same target protein, the ensemble regression can further average the predictions from individual structures to improve accuracy and robustness.

### 2.2 PST for delineating protein mutational atlas

Persistent homology [25, 26] has been widely applied to analyze the topology of point cloud data. However, it is not sensitive to homotopic shape evolution of data given by filtration. Recently, PST was proposed to reveal the homotopic shape evolution of data (Figure 2) [29]. Here we briefly introduce and compare these two topological methods.

**Figure 2:**
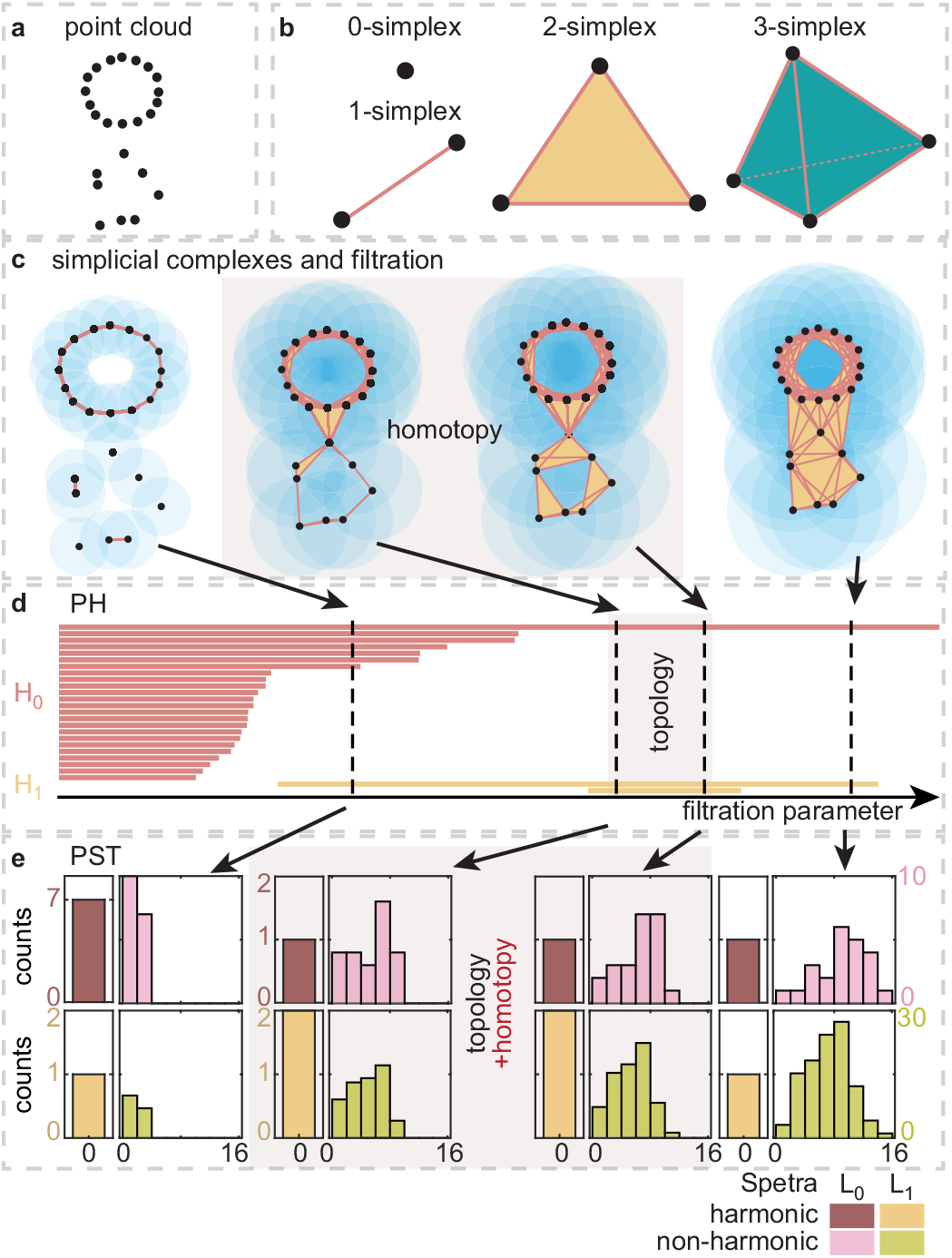
Persistent spectral theory for topological persistence and homotopic shape evolution. **(a).** Three dimensional protein structure data is given as a point cloud. **(b).** Simplex provides the basic building block of the simplicial complex. **(c).** Balls with a fixed radius are placed at each point to generate a covering of the point cloud. The simplicial complex is constructed from the covering patterns of the point cloud. Filtration generates a family of simplicial complexes with a series of increasing radii. **(d).** Persistent homology (PH) provides a topological representation from persistent barcodes at homology group *H_i_* at *i*-th dimension. **(e).** Persistent spectral theory (PST) analyzes the spectra of persistent Laplacians. The harmonic spectra reveal topological persistence as PH does, and the non-harmonic spectra capture the homotopic shape evolution of data. Abscissa shows the range of spectra, and ordinate shows the number of spectra in the range.

Atoms in a protein structure are described as a set of point cloud (Figure 2**a**). Simplicial complex glues a set of interacting nodes, called simplexes (Figure 2**b**), to construct a shape realization of the point cloud. A simplicial complex can be built from the intersection pattern defined by fixed-radius balls centered at the sample points. *Č*ech complex, Rips complex, and alpha complex are commonly used to construct simplicial complexes [25]. The filtration process creates a family of simplicial complexes by continuously growing radii of balls to establish a multiscale analysis (Figure 2**c**).

Persistent homology is a popular approach for analyzing the family of simplicial complexes obtained from filtration. For an individual simplicial complex, topological invariants, i.e., Betti numbers, are calculated as the rank of the homology group. In particular, Betti-0, Betti-1, and Betti-2 describe numbers of independent components, rings, and cavities, respectively. Persistent homology analyzes the family of simplicial complexes and tracks the persistence of topological invariants, visualized by persistent barcodes [25]. The number of *k*-th Betti bars at a fixed filtration parameter is the Betti-*k* for the corresponding simplicial complex (Figure 2**d**). Persistent homology only captures topological changes while failing to detect homotopic geometric changes. For example, two middle patterns in Figure 2**c** are topologically equivalent (i.e., no change in Figure 2**d**) but have different geometric shapes.

Alternatively, the topology of point cloud data can be described by combinatorial Laplacians [33], one kind of topological Laplacians. Loosely speaking, the 0-combinatorial Laplacian, *L*_0_, is a graph Laplacian that describes the pairwise interactions (i.e., edges, 1-simplexes) among vertices (0-simplexes) [34, 35]. The high-dimensional *k*-combinatorial Laplacian, *L_k_*, describes interactions among high-dimensional components (i.e., *k*-simplexes) upon their adjacency associated with lower- and higher-dimensional simplexes (Methods). According to the combinatorial Hodge theorem [36], the topological connectivity (or invariant) described by the combinatorial Laplacian is revealed by its kernel space or the harmonic spectra (i.e., zero eigenvalues), whereas the non-harmonic spectra, such as the Fiedler value (i.e., smallest non-zero eigenvalue), describes the so-called algebraic connectivity and homotopic shape information. Nonetheless, like homology, combinatorial Laplacian has limited descriptive power for complex data. We introduced PST to construct multiscale topological Laplacians [29]. Specifically, we used filtration to create a family of simplicial complexes at various scales and study their topological and geometric properties from a series of spectra of combinatorial Laplacians. PST not only recovers the full topological persistence of the persistent homology [30] but also deciphers the homotopic shape evolution of data. As shown in Figure 2**e**, the number of harmonic zero eigenvalues are always the same as the number of persistent barcodes in Figure 2**d**. And the non-harmonic spectra reveal both topological changes and the homotopic shape evolution when there were no topological changes (Figure 2**e**). PST comprehensively characterizes the geometry of an object from a family of frequencies manifesting evolving shapes induced by the filtration of data, which provides another illustration of the famous question: “Can one hear the shape of a drum?”, raised by Mark Kac [37].

### 2.3 Protein fitness benchmark overview

We compared the performance of machine learning models built on various featurization strategies. We benchmarked 2 topological structure-based embeddings, 9 sequence-based embeddings, and 5 evolutionary density scores on 34 benchmark deep DMS datasets across 27 proteins. In addition, TopFit was benchmarked which integrates one PST embedding, one sequence-based embedding, and one evolutionary score in its concatenated feature library. Three-dimensional structures of the target protein at the domain of interest were obtained from PDB [22], or AlphaFold [38]. X-ray structures are selected with the highest priority. Otherwise, NMR, cryo-EM (EM) or AlphaFold (AF) structures are employed depending on their availability and quality (Methods). For each dataset, 20% of the available labeled data are randomly selected as the testing set in each run. The training set is randomly sampled within the remaining 80% data. We are particularly interested in small training data to mimic the circumstances in protein engineering. The fixed number of training data is picked as the multiples of 24 ranging from 24 to 240 to mimic the smallest 24-well experimental assay. Low-N is referred to extremely small 24 training data. In addition, we also tested 80/20 train/test split and five-fold cross-validation to have direct comparisons with other methods. To ensure reproducibility, 20 independent repeats are performed for the fixed size of training data, and 10 independent repeats are performed for five-fold cross-validation or 80/20 train/test split.

The computational predictions of mutational effects are typically employed to rank and prioritize candidate proteins for experimental screening in the next round. The primary goal is to assess the ranking quality of protein fitness. The Spearman rank correlation (*ρ*) [7, 8, 39, 21, 20, 17, 18] and the normalized discounted cumulative gain (NDCG) [32, 40, 20] are two commonly used evaluating metrics in protein engineering (Methods). The Spearman correlation evaluates the overall ranking on all data where the high- and low-fitness proteins contribute equally. In contrast, NDCG quantifies the ranking quality with a higher weight on the high-fitness proteins. This may be more useful for the greedy search in selecting top samples. Since the specific fitness is not available for the unsupervised evolutionary models, a high-quality evolutionary scores can either positively or negatively rank the fitness. Indeed, absolute values of these quantities are used to evaluate evolutionary scores.

### 2.4 Comparing PST embedding with sequence-based embedding

Here we compared various embeddings in building up machine learning models over 34 datasets. Results evaluated by Spearman correlation were mainly reported in this section (Figure 3, Extended Data Figures 1, 2, 3). Similar results evaluated by NDCG were also provided (Extend Data Figures 1, 4, 5).

**Figure 3:**
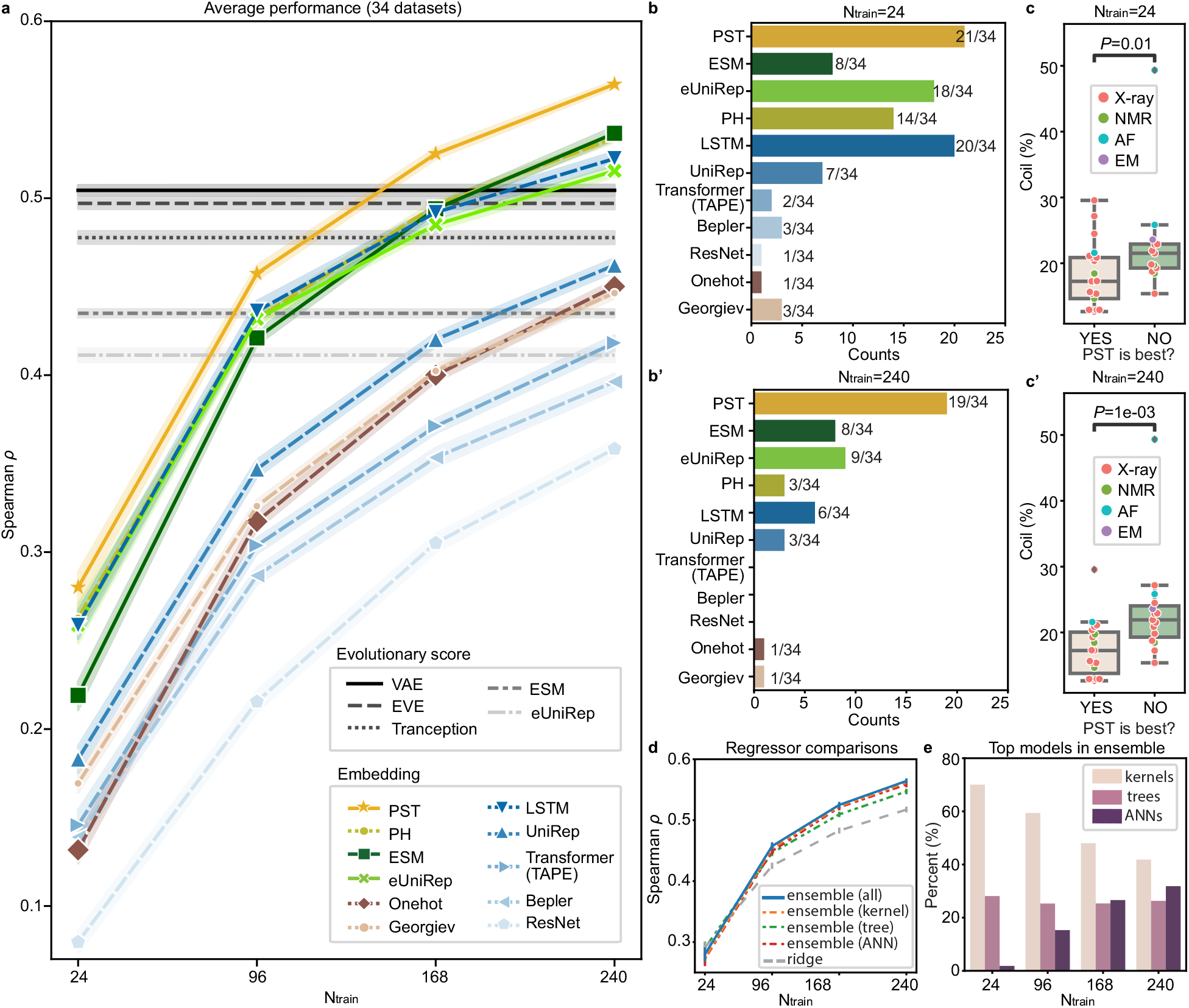
Prediction from single embedding on fitness landscape measured by Spearman correlations (*ρ*). **(a).** Line plots show average *ρ* over 34 datasets. Ensemble regression is used, except ridge regression for Georgiev and one-hot embeddings. Absolute values of *ρ* were shown for evolutionary scores. The width of shade shows 95% confidence interval from *n* = 20 independent repeats. **(b-b’).**Histograms show the frequency that an embedding is ranked as the best across 34 datasets for training data size **(b)** 24 and **(b’)** 240. For each dataset, the best embedding has average *ρ* over *n* = 20 independent repeats within the 95% confidence interval of the embedding with the highest average *ρ*. In **(a)-(b)**, Structure-based embeddings include persistent homology (PH) and PST. Sequence-based embeddings include ESM [15], eUniRep [17], Georgiev [49], UniRep [14], Bepler [13], and three TAPE models (LSTM, Transformer, and ResNet) [12]. Evolutionary scores include DeepSequence VAE [8], EVE [9], Tranception [19], ESM [15], and eUniRep [17]. **(c-c’).** Boxplots show distribution of percentages of coils for protein structure for each dataset, where boxplots were presented on 34 datasets classified into two classes depending on whether PST embedding is the best embedding. Scatter plots show same data with boxplots but for individual datasets. Onesided Mann-Whitney U test examines the statistical significance that two classes have different percentages of coils. Boxplots display five-number summary where center line shows median, upper and lower limits of the box show upper and lower quartiles, and upper and lower whiskers show the maximum and the minimum by excluding “outliers” outside the interquartile range. Training data size is **(c)** 24 and **(c’)** 240. The classes that PST is ranked as the best or not the best have sample sizes **(c)** n = 21 versus n = 13, and **(c’)** *n* = 19 versus n = 15. P-values are **(c)** *P* = 0.01 and **(c’)** *P* = 1 × 10^−3^. **(d).** Line plots show comparisons between different regression for average *ρ* over 34 datasets using PST embedding. Error bars show the 95% confidence interval from *n* = 20 independent repeats. **(e).** Histograms show model occurrence in ensemble regression over 34 datasets using the PST embedding.

For topological structure-based embeddings, the PST embedding outperforms persistent homology embedding on nearly all datasets for any size of training data (Extended Data Figure 2). PST achieves a higher average Spearman correlation than persistent homology does (Figure 3**a**). This indicates the inclusion of non-harmonic spectra from PST provides critical homotopic shape information in learning fitness landscape.

Comparisons of sequence-based embeddings were discussed in Supplementary Note 1. Overall, ESM achieves the best average performance for large training data with over 168 data, and eUniRep and TAPE LSTM achieve the best average performance for small training data. Comparisons of evolutionary scores were discussed in Supplementary Note 2. Overall, DeepSequence VAE [8] achieves the best average performance on 34 datasets (Figure 3**a**) and it ranks as the best model on a majority of datasets (21/34).

Comparing all embedding methods, the topological structure-based PST embedding achieves the best average Spearman correlation over 34 datasets for any size of training data. It outperforms the best sequence-based for training data size *N*_train_ = 24, 96, 168, and 240 with average differences Δ*ρ* = 0.02, 0.02, 0.03, and 0.03, and p-values from one-sided Mann-Whitney U test 5 × 10^−3^, 2 × 10^−5^, 3 × 10^−8^, and 4 × 10^−8^, respectively. PST always ranks most frequently as the best model over 34 datasets (Figure 3**b**, Extended Data Figure 3**b**). Categorized datasets by their taxonomy, PST shows improvement over sequence-based embeddings on eukaryote and prokaryote datasets, but no clear improvement is observed on human datasets (Supplementary Figure 1).

### 2.5 Impact of structural data quality on PST-based model

PST extracts explicit biochemical information from three-dimensional structures that is correlated to protein fitness. While sequence-based embeddings may provide less precise information by encoding them indirectly by learning from a large sequence database. It is important to understand the potential limitations of PST embedding. It is intuitive that the performance of the PST-based model may depend on the quality of structural data. Percentage of random coils in a target protein is a quantity for structural data quality, where the appearance of random coils, an unstable form of secondary structures, reduces the overall quality of the structure data. Whether PST embedding achieves the most accurate performance is correlated with the percentages of random coils (Figure 3**c**). Another quantity for structure data quality is B-factor which quantifies the atomic displacement in X-ray structures, and it similarly affects PST performance (Extended Data Figure 6).

### 2.6 Impact of ensemble regression on model generalization

In protein engineering, labeled data are iteratively collected from experiments, resulting in tens or hundreds of training data. The ensemble regression by weighting top-performing models has the adaptivity to various sizes of training data in a single model. We discuss the advantages of ensemble strategy to enhance the model accuracy, robustness, adaptivity, and scalability.

The ensemble strategy improves model accuracy (Figure 3**d**). It outperforms ridge regression on 33/34 datasets with *N*_train_ = 240 (Supplementary Figure 4), while the only underperforming Calmodulin-1 dataset exhibits poor performance *ρ* < 0.15 for both methods (Extended Data Figure 2). The ensemble of single class of models show slightly lower Spearman correlation than the full ensemble model, except the low-*N* case (Figure 3**d** and Supplementary Figure 3).

We further performed analysis on the frequency of top-performing model occurrence in the ensemble to demonstrate the adaptivity and scalability of the ensemble regression. Kernel models with a relatively weak fitting ability may be more suitable for small training data to prevent overfitting. ANN models with powerful universal approximation ability may be more accurate with sufficient training data. The tree-based models avoid overfitting by selecting useful features from the tree hierarchy independent of training data size. As results, as the training data increases, kernel models have initial large and decreasing occurrence, ANN models have opposite observations to kernel models, and the occurrence of tree-based models remains stationary (Figure 3**e** and Extended Data Figure 7).

The ensemble strategy can also be used by averaging the predictions from multiple structures to enhance the robustness against the uncertainty from structure data. The ensemble treatment on multiple NMR models improves the predictions from single NMR model (Supplementary Figure 5). But the ensemble requires much higher computational costs, which are proportional to the number of structures.

### 2.7 TopFit for navigating fitness landscape

Among PST embedding, sequence-based embedding and evolutionary scores, we showed at least one strategy can provide accurate predictions regardless of the quality of structural data and the number of training data. TopFit combines these three complementary strategies in navigating the fitness landscape.

We first built TopFit using the PST embedding, aided with DeepSequence VAE evolutionary score and ESM embedding. With 240 training data, TopFit outperforms DeepSequence VAE score, ESM embedding, and PST embedding on the majority of datasets with median differences in Spearman correlation of Δ*ρ* = 0.087, 0.083, and 0.062, respectively (Figure 4**a-b**). Similar observations were found for TopFit using PST embedding, aided with VAE score and eUniRep embedding (Extended Data Figure 8). The concatenated featurization endows TopFit advantages over other embedding strategies on most datasets regardless of training data sizes (Extended Data Figures 9–10). An exception is that VAE is more accurate for the low-N case. Similarly, the concatenated features augmented with other evolutionary scores also achieve substantial improvement (Supplementary Figure 6 and Supplementary Figure 7).

**Figure 4:**
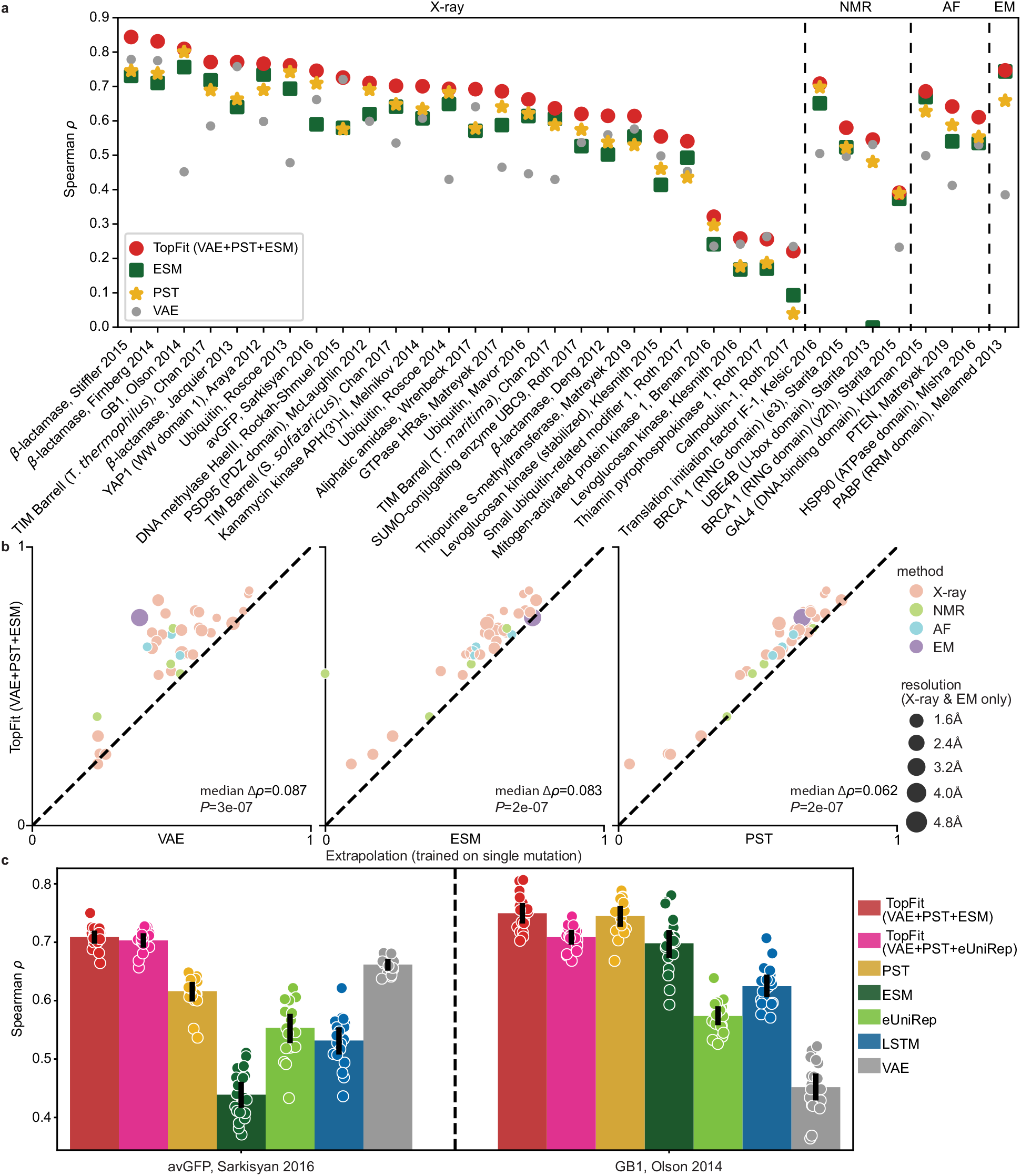
Comparisons between TopFit embeddings and other methods for fitness prediction. All supervised models use 240 labeled training data. Results are evaluated by Spearman correlation *ρ*. DeepSequence VAE takes the absolute value of *ρ*. In **(a-b)**, TopFit combines PST embedding, ESM embedding, and DeepSequence VAE score. The average *ρ* from 20 independent repeats is shown. 34 datasets are categorized by their structure modality used: X-ray, nuclear magnetic resonance (NMR), AlphaFold (AF), and cryogenic electron microscopy (EM). **(a)**. Dot plots show results across 34 datasets **(b).** Dot plots show pairwise comparison between TopFit with one method at each plot. Medians of difference for average Spearman correlation Δρ across all datasets are shown. One-sided rank-sum test determines the statistical significance that TopFit has better performance than VAE score, ESM embedding, and PST embedding for *n* = 34 datasets with p-values *P* = 0.087, 0.083, and 2 × 10^−7^, respectively. **(c).** Comparisons for extrapolation that predicts datasets with multiple mutations using the single-mutation data. Bars show average values and errorbars show 95% confidence interval from *n* = 20 independent repeats. Scatter plots show *ρ* from each repeat.

Furthermore, we carried out an extrapolation task where the model trained on single-mutation variants is used to navigate datasets including multiple-mutational variants for avGFP [41] and GB1 [42] datasets. Interestingly, we employed ridge regression due to its better performance than ensemble regression for extrapolation (Supplementary Figure 8). PST embedding achieves the best performance over other sequence-based embeddings in both datasets. TopFit further improves the performance robustness (Figure 4**c**). An exception was found on the GB1 where inclusion of poorly performing eUniRep embedding and VAE score underperforms PST embedding alone (Figure 4**c**).

Next, we compared TopFit with two existing regression models for fitness predictions (Figure 5). One is the augmented VAE model which is the best reported model combining sequence embedding and evolutionary score [20]. The other is the ECNet which combines both local and global evolutionary embeddings [21]. TopFit for comparisons uses PST embedding aided with VAE score and ESM embedding. Due to the high computational costs for the large training data size (i.e., 80/20 train/test split and five-fold cross validation), we only carry out TopFit on 27 single-mutation datasets using X-ray or AlphaFold structures. For low-N case, the augmented VAE model outperforms TopFit on 33/34 datasets by largely retaining accuracy from VAE. When more training data is available, TopFit becomes more accurate, where it outperforms the augmented VAE model in at least 19/34 datasets. For the 80/20 train/test split, TopFit completely outperforms the augmented VAE model on 25/27 datasets, and the average difference on Spearman for the rest of the two underperforming sets is relatively low (Supplementary Table 4 and Supplementary Table 5). Compared to ECNet, TopFit again shows the better performance on 25/27 datasets. Underperforming TopFit on Thiamin pyrophosphokinase dataset shows a relatively low average margin with ECNet Δ*ρ* = –0.012. The other underperforming dataset is Levoglucosan kinase with a relatively large average margin Δ*ρ* = –0.048 (Supplementary Table 4 and Supplementary Table 5). Interestingly, TopFit outperforms ECNet on a dataset generated from identical experiments using a similar but stabilized protein (i.e., Levoglucosan kinase (stabilized)). A trade-off between thermal stability and catalytic efficiency was discovered [43], and the unstablized dataset is more difficult to be predicted for all methods (Extended Data Figure 2). The quality of structure data is more sensitive to thermal stability than sequence data which leading to the outperformance of TopFit over ECNet on the stablized set and underperformance on the unstablized set.

**Figure 5:**
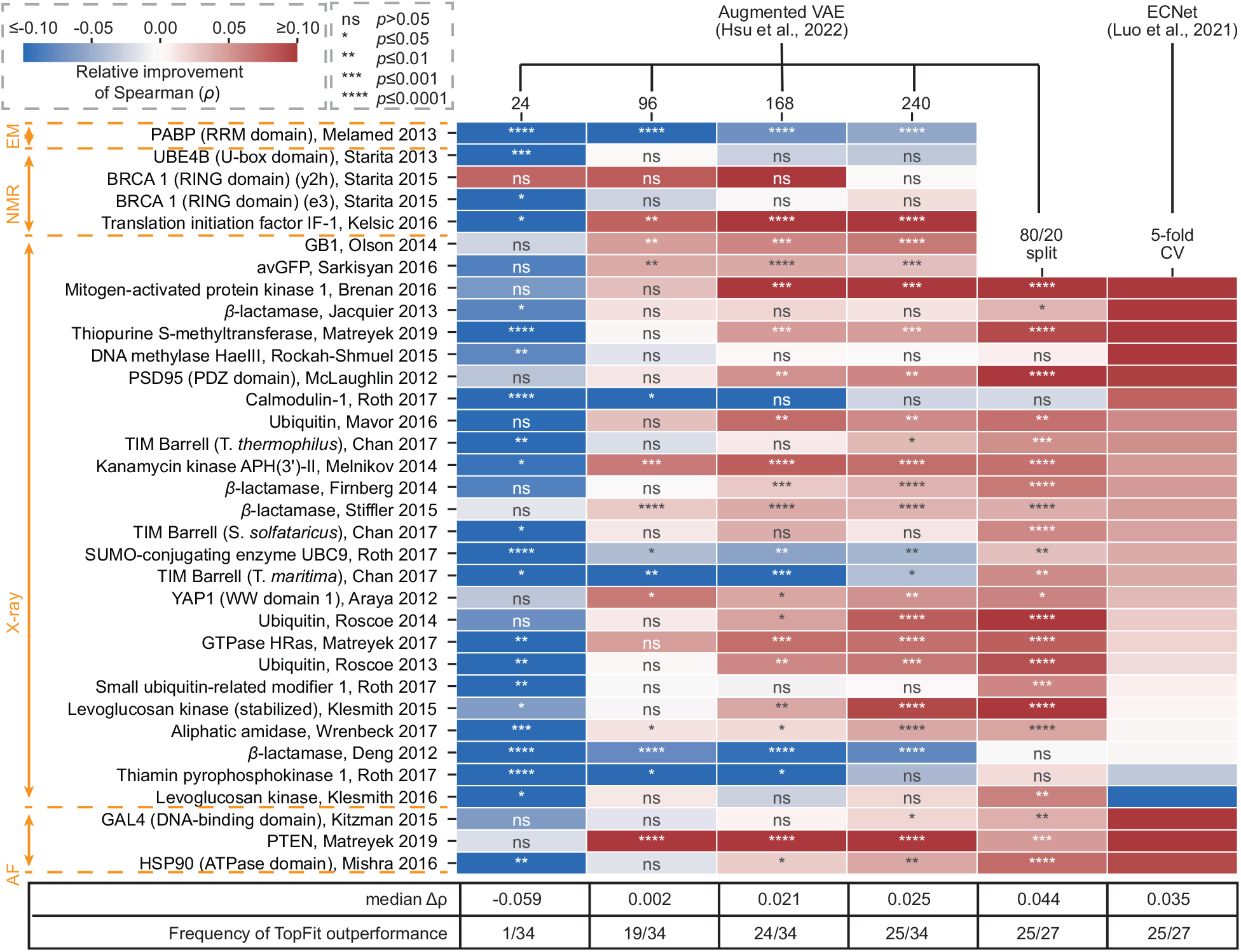
Comparisons between TopFit and other regression models for fitness predictions using Spearman correlation (*ρ*). Heatmap shows the relative improvement in *ρ* achieved by TopFit. TopFit compares with the augmented VAE model [20] for training data sizes 24, 96, 168, 240, and 80/20 train/test split. TopFit compares with ECNet [21] using five-fold cross-validation (5-fold CV). One-side Mann-Whitney U-test examines the statistical significance that two comparing methods have different *ρ*. The p-values are annotated in the heatmap using star form: ‘ns’ for not significant with *p* > 0.05, ‘*’ for *p* ≤ 0.05, ‘**’ for *p* ≤0.01, ‘***’ for *p* ≤ 0.001, ‘****’ for *p* ≤0.0001. TopFit has *n* = 20 independent repeats for 24, 96, 168, 240 training data, and *n* = 10 for 80/20 train/test split and 5-fold CV. Augmented VAE was reproduced by us with *n* = 20 independent repeats. Results for ECNet were obtained from their work with average *ρ*, and indeed p-values were not reported.

## 3 Discussions

Protein structures are intricate, diverse, variable, and adaptive to the interactive environment and thus are elusive to an appropriate theoretical description in deep mutational screening. PST improves persistent homology in offering ultimate abstraction of geometric and structural complexity of proteins. Our PST embedding for dimension 0 have relatively high dimension to provide the most critical information with basic connectivity between vertices. For dimensions 1 and 2, our PST embedding are relatively crude to retain essential information and keep the feature dimension low under the element- and site-specific strategies. The low-dimensional features can better accommodate with machine learning models in avoiding overfitting issues for small training data size, as well as reducing computational costs. Although the stable representations such as persistent landscape [44] and the persistent image [45] for persistent homology, and a potential informative representation of non-harmonic persistent spectra may provide more enrich information, they typically generate a large number of features (Supplementary Note 6). How to handle potential overfitting from these representations in biomolecular systems remains an interesting issue. The extraordinary performance of PST in predicting protein fitness opens a door for its further applications in a vast variety of interactive biomolecular systems, such as drug discovery, antibody design, molecular recognition, etc.

The quality of protein structures largely affects the performance of the PST-based model. Recently, AlphaFold structures have become a viable alternative to experimental structural data. The AlphaFold structure achieves similar accuracy with single NMR structure, while the ensemble techniques using multiple NMR structures improved the performance (Supplementary Figure 5). The combination of structure- and sequenced-based embeddings gives rise to robust predictions even for low-quality structures. In addition, structure and sequence data contain complementary information. PST delineates the specific geometry and topology of a mutation, while sequence-based featurization captures the rules of evolution from a huge library of sequences. This complementarity is valuable and generalizable to a wide variety of other biomolecular problems.

The ensemble of regressors has better generalizability by inheriting advantages from various classes of models. With larger training data, the deeper and wider neural networks or more sophisticated deep learning architectures such as attention layers [21] and convolution layers [32] may further improve the performance. The random split of training and testing sets mainly discussed in this work may be more important for late stage of protein engineering with sufficient training data. However, the initial stage may require the model generalization for extrapolations. One extrapolation task is to predict multiple-mutations from single-mutation data. A more effective supervised model is under discussion (Supplementary Figure 8). Another extrapolation task is to predict mutations at unseen sites. TopFit promises limited improvement over evolutionary scores (Supplementary Figure 9-Supplementary Figure 13; Supplementary Note 5). In general, predicting out-of-distribution data may violate the nature of supervised models, and how to build a more accurate supervised model for this task is interesting. Nonetheless, the unsupervised evolutionary scores may be more effective for extrapolation at the early protein engineering stage with insufficient training data (i.e. low-N case). In practice, the calibration of the balance between exploitation and exploration is essential for protein engineering. The commonly used supervised models including TopFit are only designed for exploitation. Models such as Gaussian process [46] and hierarchical subspace searching strategy [40, 47] may couple with TopFit mixture features to balance this trade-off.

TopFit provides a general framework to build supervised protein fitness models by combining PST structural features with sequence-based features. Arbitrary sequence-based embedding and evolutionary scores can be used. TopFit can be continuously improved by the quickly evolving state-of-art sequence-based models. In this work, we mainly tested TopFit with single type of score, especially, DeepSequence VAE. Tranception score may be more powerful than VAE for datasets with low MSA depths or viral protein datasets [19]. The equipment of any evolutionary score can largely enhance model generalization and accuracy (Supplementary Figure 6). Interestingly, inclusion of multiple scores can promise further improvement (Supplementary Note 3 and Supplementary Figure 7). Even more, one may combine multiple structure data for ensemble predictions for improvement (Supplementary Figure 5). However, all these approaches grant additional computational costs. The combined features tested in this work provide the minimal models that combine models built on distinct data resources such as local homologous sequences, large-scale sequence data, and threedimensional structure data.

## 4 Methods

### Datasets

For all DMS datasets, we followed the convention in EVmutation [7] and DeepSequence [8] to exclude data with mutational positions not covered by MSA.

The complete GB1 dataset includes almost all single (1045 mutants) and all double mutations (535917 mutants) at 55 positions [42]. In this work, we used all single mutations and double mutations with high epistasis selected in a previous work (its Supplementary Table 2) [48]. The MSA of GB1 covers the last 54 residues, and mutations that include the positions outside MSA are excluded. As a result, the GB1 dataset used in this work includes 2457 mutations.

The avGFP dataset [41] includes 54025 mutations. We followed previous works by picking up mutations at positions that have more than 30% gaps in MSA to focus on regions with sufficient evolutionary data [20, 17]. Positions 15-150 were selected out of 237 positions, resulting in 7775 mutants with numbers of mutations between 1 and 9.

Other 32 datasets were selected from DeepSequence [8]. We searched available three-dimensional structural data from UniProt [11] and PDB database [22] for each dataset. If a PDB entry largely covers the mutational domain, the PDB structure and the corresponding dataset would be selected. The full list of datasets can be found in Supplementary Data 1.

### Multiple sequence alignments

Multiple sequence alignments search the homologous sequences of the wild-type sequence in each DMS dataset. For the 32 DeepSequence datasets, we adopted the MSAs used in the original work [8]. MSAs of avGFP were obtained from [20]. The MSAs for GB1 were generated by EvCoupling server [6] by searching against the UniRep100 database with bitscore 0.4, resulting in 387 sequences that cover 54/55 positions. We also explored different bitscores for GB1 in the MSA search, but larger bitscore leads to fewer sequences and smaller bitscore fail to cover 70% positions of the wild-type sequence.

### Sequence-based models

Multiple sequence-based evolutionary models were used in this work to generate sequence embeddings or evolutionary scores.

#### Constant embedding

Two constant embeddings, one-hot and Georgiev, were used in this work. The one-hot embedding is an uninformative categorical strategy without biochemical information. We consider 20 canonical amino acids and a sequence of length *L* is encoded as a 20*L* vector. Georgiev embedding [49] provides a 19-dimensional representation for over 500 physicochemical quantities of amino acid in the AAIndex database [50]. A sequence is encoded as a 19*L* vector.

#### ESM Transformer

The ESM-1b Transformer [15] is a transformer model [51, 52] that learns 250 million sequences using a masked filling procedure. Its architecture contains 34 layers with 650 million parameters.

The ESM Transformer was used to generate sequence embedding. At each layer, the ESM model encodes a sequence with length *L* into a matrix with dimensions 1280 × *L* by excluding the start and terminal tokens. In this work, we took the sequence representation from the final (34-th) layer and average over the axis for sequence length, resulting in a 1280-component vector.

The ESM transformer was also used to generate evolutionary score to predict fitness. Specifically, the difference in conditional log-likelihoods between the mutated amino acids and the wildtype amino acids is taken as the evolutionary score. Given a sequence with length *L*, *s* = *s*_1_*s*_2_ ··· *s_L_*, the masked filling model generates probability distributions for amino acids at masked positions. For a masked *i*-th position residue *s_i_*, the distribution is given by a conditional probability *P*(*s_i_*|*s*^(−*i*)^), where *s*^(−*i*)^ is the remaining sequence excluding the masked *i*-th position. To reduce the computational cost, the pseudo-log-likelihoods (PLLs) estimate the exact log-likelihood of a given sequence:

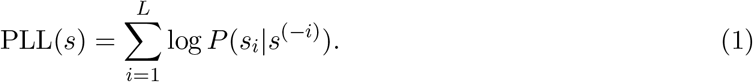

The evolutionary score is approximated by the difference of PLLs between wild-type (*w*) and mutant (*m*) sequences:

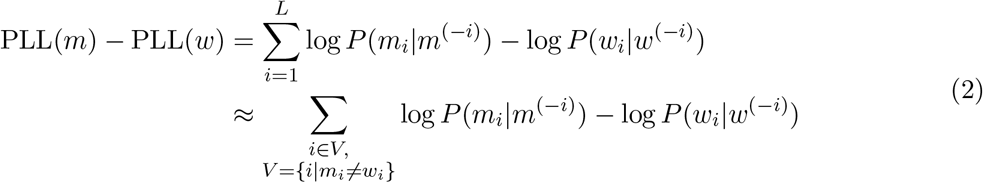

The evolutionary score can be approximated by the second line in the above equation by summing up the difference between mutant and wild-type pseudo-log-likelihoods at mutated positions since the PLLs at non-mutated positions have similar values. We conducted this approximation followed the previous work [20, 32] to save computational cost.

#### UniRep and eUniRep

UniRep [14] is the mLSTM model [53] learns from 27 million sequences from UniRef50 data. To adapt the UniRep to specific protein families, the “evo-tuning” procedure is introduced to fine-tune the models using evolutionary sequences (i.e. homologous sequences from MSA). The resulting eUniRep [17] model includes more specific information for the target sequence.

These mLSTM models can generate a sequence embedding by encoding a sequence into a 1900 × *L* matrix using the final hidden layer. The sequence embedding with 1900 dimension is obtained by average over the sequence length axis.

In addition, the evolutionary score can be generated. The mLSTM models can also be viewed as sequence distribution. The evolutionary score is also given by the log-likelihood. Given the first *i* amino acids of a sequence, the conditional probabilities of the next amino acid *P*(*s*_*i*+1_|*s*_1_ ··· *s_i_*) is parametrized by the mLSTM model. The sequence probability is given by

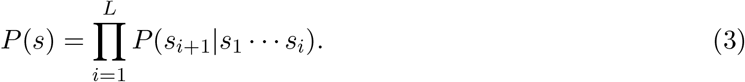

The log likelihood is then taken as the evolutionary score.

In the fine-tune procedure, we followed the protocols and hyperparameters used in the previous work [17, 20]. The evolutionary sequences from MSA were split into 80% training set and 20% validation set during the fine-tuning. The learning rate is taken as 1 × 10^−5^ and the number of epochs is 10,000. We performed the fine-tuned procedure if the fine-tuned model is not performed by previous work [20].

#### TAPE embedding

The TAPE models contain multiple pretrained protein language models to generate sequence embeddings [12]. Three models were created by TAPE, including ResNet [54], Transformer [51, 52], and LSTM [53] models. Two existing LSTM-based models, UniRep [14] and Bepler [13], were also implemented. The sequence embeddings generated from the latent space representation have dimensions 256, 512, 2048, 100, and 1900 for ResNet, Transformer, LSTM, Bepler, and UniRep, respectively. Both TAPE and the original UniRep implemented the UniRep model. In this work, we only used the one from the original implementation.

#### DeepSequence VAE evolutionary score

DeepSequence VAE [8] is a variational autoencoder model that learns sequence distribution from MSA. Since the log-likelihood is intractable, the evidence lower bound (ELBO) is used to estimate the sequence log-likelihood to predict the sequence mutational effect. Followed by the original DeepSequence, we trained five VAE models with different random seeds and generated 400 ELBO samples for each model, and the average of all 2000 ELBO samples is used. In this work, we only performed DeepSequence on avGFP, GB1, and two BRCA1 datasets. Other VAE scores were obtained from data provided in DeepSequence [8].

#### Tranception evolutionary score

Tranception [19] is an advanced self-supervised transformer-based protein language model with 700 millions parameters that promotes specialization across attention heads for enhanced protein modeling. The pre-trained model rank mutations using PLLs followed by the protocol in ESM score (Eq. (5)). Tranception employs a special bi-directional scoring to have improvement over the unidirectional scoring used in traditional Transformer-based model. It uses training data augmented by all sequences and their reverse. The evolutionary score from autoregressive inference mode, log *P_A_*(*s*), is obtained by averaging PLL scores for sequences in both forward and reverse order. In addition to the global autoregressive score, Tranception further proposed a local evolutionary score, retrieval inference, that is based on the empirical distribution of amino acids in MSAs. It calculates a local log likelihood, log *P_R_*(*s*), from retrieval inference for sequence *s*. Tranception score includes both global and local scores via a weighted geometric average in probability space:

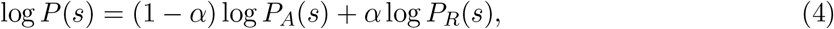

where *α* is the retrieval inference weight, and we used *α* = 0.6 which was reported as the optimal value for the validation data. The Tranception work has calculated scores for 32 datasets used in our work. We calculated the scores for the two BRCA1 datasets for RING domain by using the MSAs obtained from the description above.

#### EVE evolutionary score

Evolutionary model of variant effect (EVE) is a Bayesian variational autoencoder model that is self-trained on MSAs to learn the sequence distribution. The ELBO is used to estimate sequence log-likelihood to predict sequence mutational effect. We adopted the ensemble EVE scores from Tranception work [19] for 32 datasets. The scores for the remaining two BRCA1 datasets for RING domain were adopted from EVE work [9].

### Structural data preparation

The raw wild-type structure data is obtained from the PDB database [22] or AlphaFold [38]. VMD removes the water and selects the target chain in the corresponding DMS dataset [55]. Jackal package [56], a protein structure modeling tool, is used to optimize wild-type structures and generate mutant structures. The force parameter CHARMM22 is used in this work. Missing hydrogen atoms of the raw structure data are added using **profix** function in Jackal [56]:

~~~
./profix -fix 0 $pdbfile
~~~

where $pdbfile is the file name of the structure data. The raw structure data may have a few mutations away from the wild-type sequence. The wild-type structure is generated from the raw structure via mutations using **scap** with fixed backbone:

~~~
./scap -ini 20 -min 4 $mutant_list $pdbfile
~~~

where $mutant_list is the document for mutation information. Then the structure is optimized with 20 rounds of side-chain conformations minimization using **scap** in Jackal [56]:

~~~
./scap -ini 20 -min 4 $pdbfile
~~~

From the wild-type structure, each mutant structure in the dataset is obtained from using **scap** with fixed backbone, and further optimized by conformation minimization.

For both wild-type and mutant structures, the local structure are extracted by VMD [55] for the preparation of site-specific analysis. Specifically, atoms belong to mutational sites and atoms near the mutational sites within distance of 13 Å are selected.

### Simplicial complex and chain complex

Graph is a representation for a point cloud consisting of vertices and edges for modeling pairwise interactions, such as atoms and bonds in molecules. Simplicial complex, the generalization of graph, constructs more enriched shapes to include high dimensional objects. A simplicial complex is composed of simplexes up to certain dimensions. A *k*-simplex, *σ^k^*, is a convex hull of *k* + 1 affinely independent points *v*_0_, *v*_1_, *v*_2_, ···, *v_k_*:

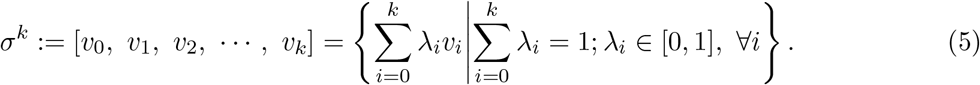

In Euclidean space, 0-simplex is a point, 1-simplex is an edge, 2-simplex is a triangle, and 3-simplex is a tetrahedron. The *k*-simplex can describe abstract simplex for *k* > 3.

A subset of the *k* + 1 vertices of a *k*-simplex, *σ^k^*, with *m* + 1 vertices forming a convex hull in a lower dimension and is called an *m*-face of the *k*-simplex *σ^m^*, denoted as *σ^m^* ⊂ *σ^k^*. A simplicial complex *K* is a finite collection of simplexes satisfying two conditions:

1. Any face of a simplex in *K* is also in *K*.
2. The intersection of any two simplexes in *K* is either empty or a shared face.

The interactions between two simplexes can be described by adjacency. For example, in graph theory, two vertices (0-simplexes) are adjacent if they share a common edge (1-simplex). Adjacency for *k*-simplexes with *k* > 0 includes both upper and lower adjacency. Two distinct *k*-simplexes, *σ*_1_ and *σ*_2_, in *K* are upper adjacent, denoted *σ*_1_ ~_*U*_ *σ*_2_, if both are faces of a (*k* + 1)-simplex in *K*, called a common upper simplex. Two distinct *k*-simplexes, *σ*_1_ and *σ*_2_, in *K* are lower adjacent, denoted *σ*_1_ ~_*L*_ *σ*_2_, if they share a common (*k* – 1)-simplex as their face, called a common lower simplex. Either common upper simplex or common lower simplex is unique for two upper or lower adjacent simplexes. The upper degree of a *k*-simplex, deg_*U*_(*σ^k^*), is the number of (*k* + 1)-simplexes in *K* of which *σ^k^* is a face; the lower degree of a *k*-simplex, deg_*L*_(*σ^k^*), is the number of nonempty (*k* – 1)-simplexes in *K* that are faces of *σ^k^*, which is always *k* + 1. The degree of *k*-simplex (*k* > 0) is defined as the sum of its upper and lower degree

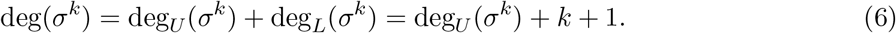

For *k* = 0, the degree of a vertex is:

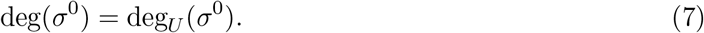

A simplex has orientation determined by the ordering of its vertices, except 0-simplex. For example, clockwise and anticlockwise orderings of three vertices determine the two orientation of a triangle. Two simplexes, *σ*_1_ and *σ*_2_, defined on the same vertices are similarly oriented if their orderings of vertices differ from an even number of permutations, otherwise, they are dissimilarly oriented.

Algebraic topology provides a tool to calculate simplicial complex. A *k*-chain is a formal sum of oriented *k*-simplexes in *K* with coefficients on 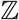. The set of all *k*-chains of simplicial complex *K* together with the addition operation on 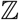 constructs a free Abelian group *C_k_*(*K*), called chain group. To link chain groups from different dimensions, the *k*-boundary operator, *∂_k_*: *C_k_*(*K*) → *C*_*k*–1_(*K*), maps a *k*-chain in the form of a linear combination of *k*-simplexes to the same linear combination of the boundaries of the *k*-simplexes. For a simple example where the *k*-chain has one oriented *k*-simplex spanned by *k* + 1 vertices as defined in Eq. (5), its boundary operator is defined as the formal sum of its all (*k* – 1)-faces:

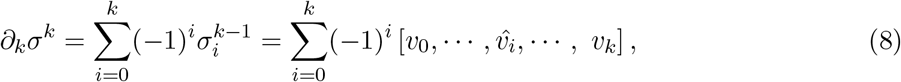

where 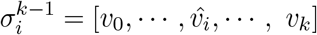 is the (*k* – 1)-simplex with its vertex *v_i_* being removed. The most important topological property is that a boundary has no boundary: 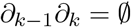.

A sequence of chain groups connected by boundary operators defines the chain complex:

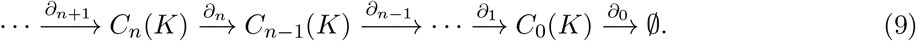

When *n* exceeds the dimension of *K, C_n_*(*K*) is an empty vector space and the corresponding boundary operator is a zero map.

### Combinatorial Laplacian

For *k*-boundary operator *•_k_*: *C_k_* → *C*_*k*–1_ in *K*, let 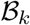 be the matrix representation of this operator relative to the standard bases for *C_k_* and *C*_*k*–1_ in *K*. 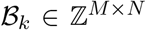 is the matrix representation of boundary operator under the standard bases 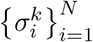 and 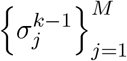 of *C_k_* and *C*_*k*–1_. Associated with the boundary operator *∂_k_*, the adjoint boundary operator is 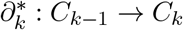, where its matrix representation is the transpose of the matrix, 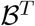, with respect to the same ordered bases to the boundary operator.

The *k*-combinatorial Laplacian, a topological Laplacian, is a linear operator Δ_*k*_: *C_k_*(*K*) → *C_k_*(*K*)

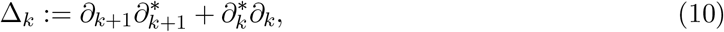

and its matrix representation, *L_k_*, is given by

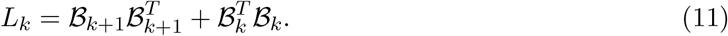

In particular, the 0-combinatorial Laplacian (i.e. graph Laplacian) is given as follows since *∂*_0_ is an zero map:

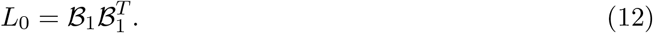

The elements of *k*-combinatorial Laplaicn matrices are

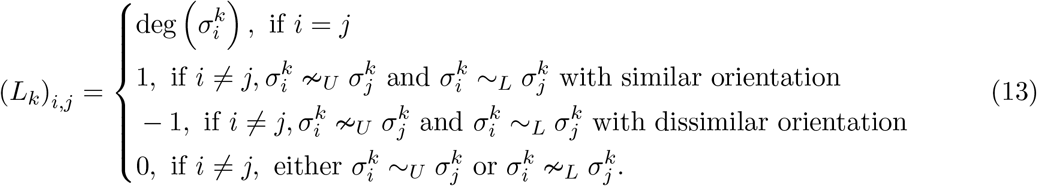

For *k* = 0, the graph Laplacian matrix *L*_0_ is

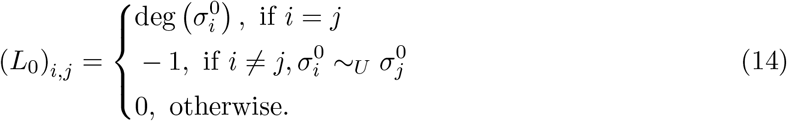

The multiplicity of zero spectra of *L_k_* gives the Betti-*k* number, according to combinatorial Hodge theorem [36]:

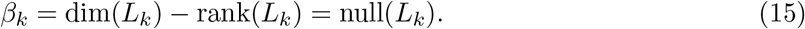

The Betti numbers describe topological invariants. Specifically, *β*_0_, *β*_1_, and *β*_2_ may be regarded as the numbers of independent components, rings, and cavities, respectively.

### Persistent spectral theory (PST)

The PST is facilitated by filtration, which introduces a multiscale analysis of the point cloud. The filtration is a family of simplicial complexes, 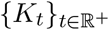, parameterized by a single parameter *t* (i.e., the radius of ball in Figure 2**c**) and ordered by inclusion. There are several properties of the family of simplicial complexes:

1. For two values of parameters *t′* < *t″*, we have *K_t′_* ⊆ *K_t″_*.
2. There are only finite number of shape changes in the filtration. And we can find at most *n* filtration parameters such that

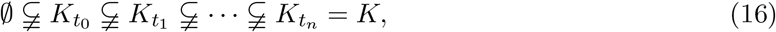

where *K* is the largest simplicial complex can be obtained from the filtration. Suppose *t_i_* is the smallest filtration parameter where we observe the *i*-th shape changes. Then for any filtration parameter *t*, the simplicial complex is corresponding to one simplicial complex in Eq. (16):

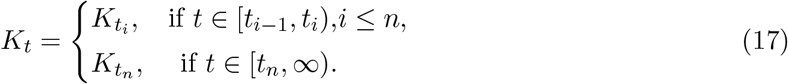

There are various simplicial complex that can be used to construct the filtration, such as Rips complex, *Č*ech complex, and Alpha complex. For example, the Rips complex of *K* with radius *t* consists of all simplexes with diameter at most 2*t*:

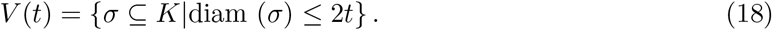

In the filtration, a family of chain complexes can be constructed:

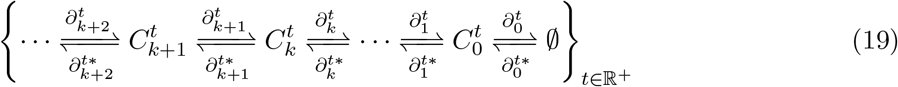

where 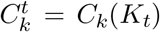 is the chain group for subcomplex *K_t_*, and its *k*-boundary operator is 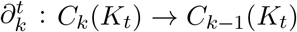. The family of any *k*-dimensional chain complexes 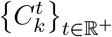 has similar two properties for the simplicial complexes 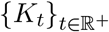 in filtration.

The homotopic shape changes with a small increment of filtration parameter may be subject to noise from the data. The persistence may be considered to enhance the robustness when calculating the Laplacian. First, we define the *p*-persistent chain group 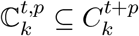 whose boundary is in 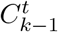:

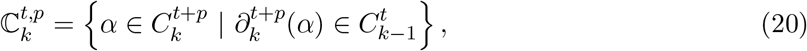

where 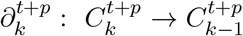 is the *k*-boundary operator for chain group 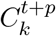. Then we can define a *p*-persistent boundary operator, 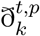, as the restriction of 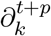 on the *p*-persistent chain group 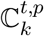:

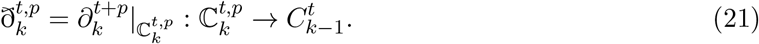

Then PST defines a family of *p*-persistent *k*-combinatorial Laplacian operators 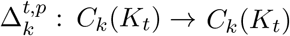 [29] which is defined as

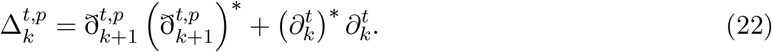

We denote 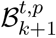 and 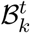 as the matrix representations for boundary operators 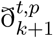 and 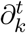, respectively. Then the Laplacian matrix for 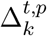 is

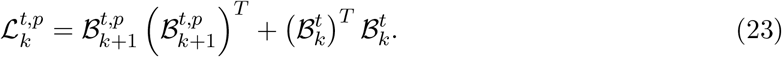

Since the Laplacian matrix, 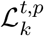, is positive-semidefinite, its spectra are all real and non-negative

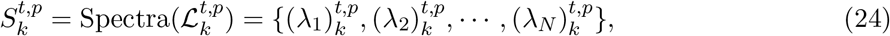

where *N* is the dimension of a standard basis for 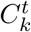, and 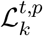 has dimension *N* × *N*. The *k*-persistent Betti number 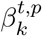 can be obtained from the multiplicity of harmonic spectra of 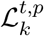:

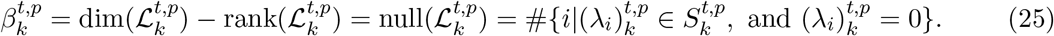

In addition, the rest of the spectra, i.e., the non-harmonic part, capture additional geometric information. The family of spectra of the persistent Laplacians reveals the homotopic shape evolution (Figure 2**e**).

### Site- and element-specific PST for protein mutation embedding

The site- and element-specific strategies are used to generate PST embedding to study mutation-induced structural changes. The site-specific strategy focuses on mutational sites to reduce computational complexity. The element-specific strategy considers the pairwise interactions between two atomic groups to encode appropriate physical and chemical interactions in embedding. For examples, hydrogen bonds can be inferred from the spectra of oxygen-nitrogen persistent Laplacians whereas hydrophobicity can be extracted from carbon-carbon pairs.

Specifically, the atoms in a protein are classified into various subsets:

1. 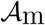: atoms in the mutation site.
2. 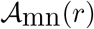: atoms in the neighborhood of the mutation site within a cut-off distance, *r*.
3. 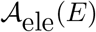: atoms in the system with element type, *E*.

An atomic group combines the atoms within 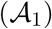 and near the mutational sites 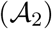:

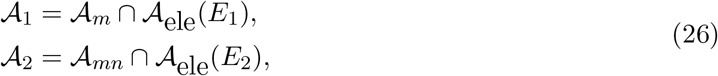

where element types *E*_1_ and *E*_2_ can be *C, N, O*, and all heavy atoms. Nine single atomic pairs are constructed where 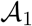 and 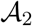 pick one of heavy elements in {*C, N, O*} each time. And a heavy atomic pair is constructed where both 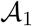 and 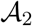 represent all heavy elements {*C, N, O*}.

PST analyzes the union of atoms 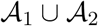 to embed critical biophysical atomic interactions. The Euclidean distance *D_e_* or a bipartite distance *D*_mod_ is used. The bipartite distance excludes interactions between the atoms from the same set:

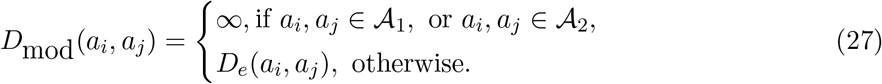

For 0-dimension, filtration using Rips complex with *D*_mod_ distance is used. The 0-dimensional PST features are generated at fixed scales. Specifically, 0-persistent Laplacian features are generated at 10 fixed filtration parameters: 2Å, 3Å,· · ·, 11A. The short scales below 2Å are excluded as bond-breaking events are irrelevant to the protein fitness of our interest. For each Laplacian, we count the number of harmonic spectra and calculate 5 statistical values of non-harmonic spectra: minimum, maximum, mean, standard deviation, and sum. A 60-dimensional vector is generated. This type of features is calculated on 9 atomic pairs. For multiple mutational sites, the features are summed up over mutational sites. Finally, the dimension of 0-dimensional PST features for a protein is 540.

For 1- or 2-dimension, filtration using alpha complex with *D_e_* distance is used. The local protein structure has a small number of atoms that can generate a limited number of high-dimensional simplexes, leading to negligible shape changes. As a result, we only extracted features from harmonic spectra of persistent Laplacians coding topological invariants for the high-dimensional interactions. The persistence of the harmonic spectra is represented by the persistent barcode (Figure 2) and implemented by GUDHI [57]. The topological feature vectors are generated from the statistics of bar lengths, births, and deaths. First, bars with lengths lower than 0.1^Å^ are excluded since they do not have a clear physical interpretation. The remaining bars are used to construct the featurization vector with: 1) sum, maximum, and mean for lengths of bars; 2) minimum and maximum for the birth values of bars; and 3) minimum and maximum for the death values of bars. Each set of point clouds leads to a 7-dimensional vector. These features are calculated on 9 single atomic pairs and 1 heavy atomic pair. The dimension of 1- and 2-dimensional PST feature vectors for a protein is 140. Overall, the dimension of PST embedding is 2040 by concatenating features at different dimensions for wild-type, mutant, and their difference.

In additional to the featurization described above, we also tested different representations of PST and persistent homology features. The persistent landscape is a popular vector representation for persistent homology at dimension 1 and 2. It is implemented by GUDHI in this work [57]. We use 3 landscapes and resolution 10 for each point cloud, and it results in a 30-dimensional vector. Using the same strategy describe for dimensions 1 and 2, we generated features at 10 point clouds using the element- and site-specific strategies. The persistent landscape features are calculated for wildtype, mutant, and their difference for both dimensions 1 and 2, which results in a 1800-dimensional vector for each mutational entry. We also consider the non-harmonic spectra for *p*-persistent Laplacians at dimensions 1 and 2. They are calculated at 10 fixed filtration parameters: 2Å, 3A,···, 11Åusing HERMES package [58]. Five statistical values of non-harmonic spectra: minimum, maximum, mean, standard deviation, and sum, are used for vectorization. It results in a 50-dimensional vector for each point cloud. With the same strategy using in persistent landscape, this approach results in a 3000-dimensional vector. These two types of featurizations are not used in the main results. They are discussed in Supplementary Note 6.

### Persistent homology (PH)

Similar to PST, persistent homology presents a multiscale analysis via filtration. Persistent homology uses homology groups to describe the persistence of topological invariants. As a result, persistent homology only provides the harmonic spectral information of PST.

The site- and element-specific persistent homology features are generated similar to those of PST. They use identical filtration construction. For the 0-dimension, the filtration parameter is discretized into bins with length 1Å in the range of [2,11], namely, [2, 3), [3, 4), ···, [10,11)^Å^. For each bin, we count the numbers of persistent bars, resulting in a 9-dimensional vector for each point cloud. This type of features is calculated on 9 single atomic pairs. Indeed, the dimension of 0-dimensional persistent homology features for a protein is 81. For 1- or 2-dimension, the identical featurization from the statistics of persistent bars in PST is used. Persistent homology embedding combines features at different dimensions as described above and concatenated for wildtype, mutant, and their difference, resulting in a 663-dimensional vector.

### Ensemble regression

Following by the previous work in machine learning-assisted protein engineering [32], we employed multiple regressors to build the ensemble regression. Features are first normalized by *StandardScaler*() in scikit-learn packages [59]. Each regressor is evaluated on training data by five-fold cross-validation using the root mean square error (RMSE). The Bayesian optimization is performed to find the best-performing hyperparameters for each regressor [60]. Then, top *N* models are picked up based on the RMSE metric. The ensemble regression averages the predictions from top *N* regressors to predict the fitness. There are 18 regressors including 3 artificial neural networks (ANNs), 10 kernel models, and 5 tree-based models (Figure 1**c**).

These 18 models were all used in ensemble regression for the small fixed-size training data (i.e., from 24 to 240), and top *N* = 3 models were selected and averaged. For the ensemble of the single type of regressors, kernel, tree, or ANN, *N* = 3 is also used. For 80/20 train/test data split or five-fold cross-validation, we only ensemble 3 tree-based models and 2 ANN models for adaption to large training data, and all *N* = 5 models are averaged to enhance the generalization of the model. In addition, the ANN architectures were made wider and deeper to fit with large training data. Lists of models, default and optimization ranges of hyperparameters can be found in Supplementary Table 1-Supplementary Table 3.

#### Kernel-based models

We employed multiple kernel-based models. These types of models have great generalization ability to the small size of training data, especially for the linear models. Models implemented by scikit-learn packages [59], including ridge regression, kernel regression, Bayesian ridge regression, linear support vector regression, Bayesian ARD regression, cross-validated Lasso, SGDRegressor, *k*-nearest neighbors regressor, and ElasticNet. In addition, we also included a boosting linear model in XGBoost [61].

Particularly, for ridge regression and kernel regression, we followed the strategy used in augmented models [20] to make the evolutionary score practically unregularized (using regularization coefficient 10^−8^). Otherwise, features are concatenated and treated equally in the model.

#### Tree-based models

The tree models can rank and keep important features. Their great fitting and generalization ability lead to robust and accurate performance for any size of training data. In utilizing the tree-based models, we treat all features equally. The important features such as evolutionary scores would be recognized by the tree models and contribute more to predictions. In this work, we employed four scikit-learn models [59]: decision tree, gradient boosting tree regressor, random forest regressor, bagging regressor, and one XGBoost tree model [61].

#### ANN models

The ANN models have universal approximation ability to learn data. Particularly, with large size of training data, ANN models show higher accuracy and generalizability than other machine learning models. We employed deep and wide neural network architectures to build ANN models [62]. The structure- and sequence-based embeddings are concatenated and fed into a series of hidden fully connected layers. Since embeddings provide relatively lower-level features than evolutionary scores, embedding features are fed-forward by several hidden layers, and evolutionary score concatenates with the last hidden layers, and the entire last layer is fed-forward to the output layer.

Each hidden layer except for the final layer sequentially combines a fully connected layer, “ReLu” activation function, a batch normalization layer, and a dropout layer. At the final hidden layer, the hidden units are concatenated with the evolutionary score, and a batch normalization layer and a fully connected layer are sequentially used to obtain the output. We took the Adam optimizer with maximal epochs *num-epochs*. An early stopping criterion is used where the training will be stopped if no decrease of *tol* is observed for training errors over *patience* epochs.

For fixed and small sizes of training data 24, 96, 168, and 240, three ANN models have one, two, and three hidden layers. For 80/20 split or five-fold cross-validation, the ANN models include more neurons at each layer, and the three ANN models have one, three, and five hidden layers. Specific hyperparameters can be found in Supplementary Table 3.

### Secondary structure predictions

Secondary structures were predicted by the dictionary of protein secondary structures (DSSP) [63]. Secondary structures are assigned based on hydrogen bonding patterns. We performed DSSP on wild-type protein for each dataset. DSSP classifies amino acids in a protein into 8 categories: 3-turn helix (G), 4-turn helix (H), 5-turn helix (I), extended strand in parallel and/or anti-parallel *β*-sheet conformation (E), residue in isolated *β*-bridge (B), bend (S), and coil (C). The experiments usually have low accuracy in measuring the structure of coils and the errors will propagate to the entire structure. We then classify these 8 categories into two types: coil and non-coil. And these two types of secondary structures can largely indicate the potential accuracy of the structure data.

### B-factors

The B-factor provides a direct measurement of the structural flexibility experiment. The B-factor, sometimes called the Debye–Waller factor, temperature factor, or atomic displacement parameter, is used in protein crystallography to describe the attenuation of X-ray or neutron scattering caused by thermal motion. In each X-ray structure, the B-factor is provided for each atom. In this work, we extracted B-factor at alpha carbons to represent the quality of each amino acid. Then we have a vector of B-factor with the same length with number of amino acids in a protein. Since high B-factor indicates low experimental accuracy, the third quartile (*Q*_3_), presented in this work, indicates the accuracy of those residues with relatively high B-factor, consequently, reflects the quality of less accurate domain in a X-ray structure.

### Statistical tests

We used two one-sided statistical tests to compare the performance of different methods. For two sets of samples *X* and *Y*, we make the one-side hypothesis based on their average values. For example, if *X* has larger average than *Y*, we make the hypothesis that *X* is greater than *Y*. The p-values are calculated to evaluate the statistical significance of the null hypothesis.

#### Mann-Whitney U-test

Mann-Whitney U-test is a nonparametric test of the null hypothesis that, for randomly selected values of *X* and *Y* from two populations, the probability of *X* being greater than *Y* is equal to the probability of *Y* being greater than *X*. The alternative hypothesis is that one population is stochastically greater than the other.

#### Wilcoxon signed-rank test

Wilcoxon signed-rank test is also a nonparametric test. It is particularly designed for two pairs of dependent samples. For example, we used it to compare the performance of two methods over all datasets which results in paired samples (e.g., Figure 4**b**). Otherwise, the Mann-Whitney U-test is used for unpaired samples.

### Evaluating metrics

#### Spearman correlation (ρ)

Spearman correlation assesses monotonic relationships between two variables. The Spearman correlation between two variables will be high when observations have a similar rank between the two variables and low when observations have a dissimilar rank between the two variables. It is defined as the Pearson correlation coefficient between the rank variables:

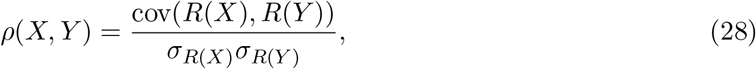

where cov is the covariance between two variables, *σ* is the standard deviation, and *R*(*X*) is the rank variable of *X*. The Spearman correlation is used to evaluate the rank correlation between ground truth and predicted fitness.

#### Normalized discounted cumulative gain (NDCG)

NDCG is another measure of ranking quality [64]. The highly ranked samples have a higher contribution to NDCG. It evaluates the ranking quality by focusing on the top samples. Especially, it is useful for greedy search in protein engineering which requires selecting top predicted samples. The predicted values are first sorted into a descending order 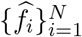 and the corresponding ground truth values are 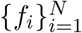. The discounted cumulative gain (DCG) is defined as

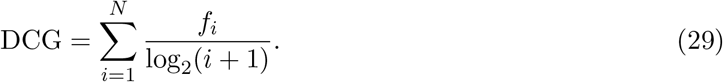

The ground truth fitness contributes more to the sum if its prediction has a higher rank. The DCG is then normalized into a range between 0 and 1, where 1 indicates perfect ranking quality. The normalized DCG (NDCG) is defined as

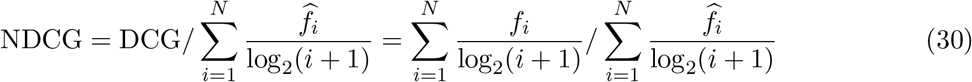

#### Root mean square error (RMSE)

RMSE is used to measure the accuracy of the regressor. Particularly, we used it to rank models in hyperparameter optimization and model selection in the ensemble regression. Since our tasks focus on the ranking quality of the data, RMSE is not used to evaluated fitness predictions in this work.

## Data Availability

There are 34 DMS datasets with experimentally measured fitness used in this work including: 32 DeepSequence datasets [8], avGFP dataset [41], and GB1 dataset [42]. The original data sources of the 32 DeepSequence datasets are provided in Supplementary Data 1 and Supplementary Note 7.

Structure data were obtained from PDB database [22] and AlphaFold [38], and the specific entry ID was provided in Supplementary Data 1.

The data analyzed and generated in this work, including fitness data, processed structure data, multiple sequence alignments, fine-tune parameters for eUniRep models, and fitness predictions from TopFit are available at https://github.com/WeilabMSU/TopFit.

Source data for Figures 3, 4, 5, and Extended Data Figures 1, 3, 5, 6, 7, 8, 9, 10 is available with this manuscript.

## Code Availability

All source codes and models are publicly available at https://github.com/WeilabMSU/TopFit [65].

## Acknowledgments

This work was supported in part by NIH grants R01GM126189 and R01AI164266, NSF grants DMS-2052983, DMS-1761320, and IIS-1900473, NASA grant 80NSSC21M0023, Michigan Economic Development Corporation, MSU Foundation, Bristol-Myers Squibb 65109, and Pfizer. We thank Chloe Hsu and Jennifer Listgarten for helpful discussions.

## Author Contributions Statement

All authors conceived this work, and contributed to the original draft, review and editing. Y.Q. performed experiments and analyzed data. G.W.W. provided supervision and resources and acquired funding.

## Competing interests Statement

The authors declare no competing interests.

## Figure Legends/Captions

**Extended Data Figure 1.**
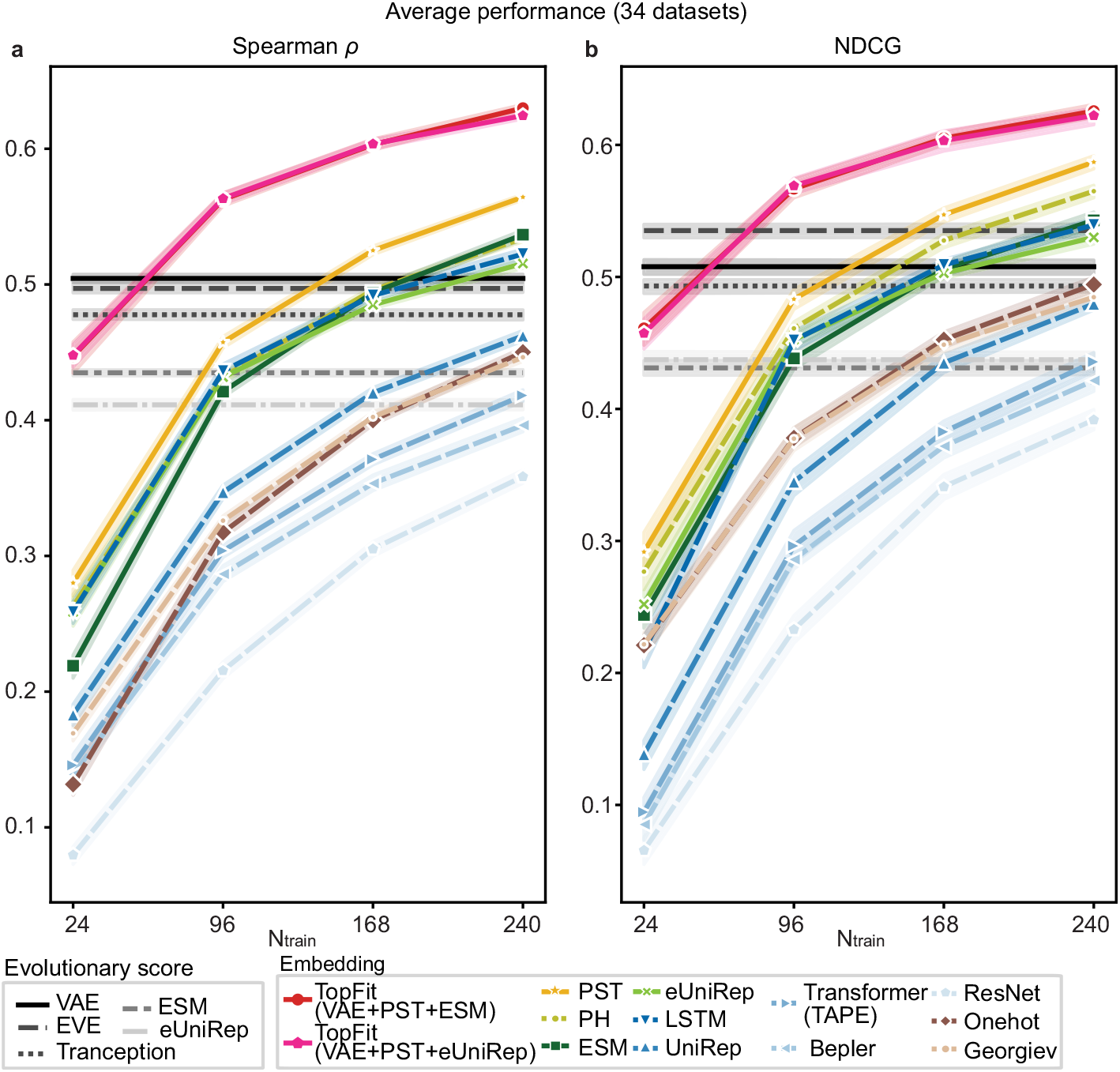
The average performance of various models over 34 datasets. It is the supplement of Figure 3**a**. **(a).** Line plots show identical data with Figure 3**a** with additional data for two TopFit strategies. **(b).** Results are evaluated by NDCG. In (**a**-**b**), ensemble regression is used, except ridge regression for Georgiev and one-hot embeddings. Absolute values of ρ were shown for evolutionary scores. The width of shade shows 95% confidence interval from n = 20 independent repeats. Evolutionary scores use absolute values for corresponding quantities.

**Extended Data Figure 2.**
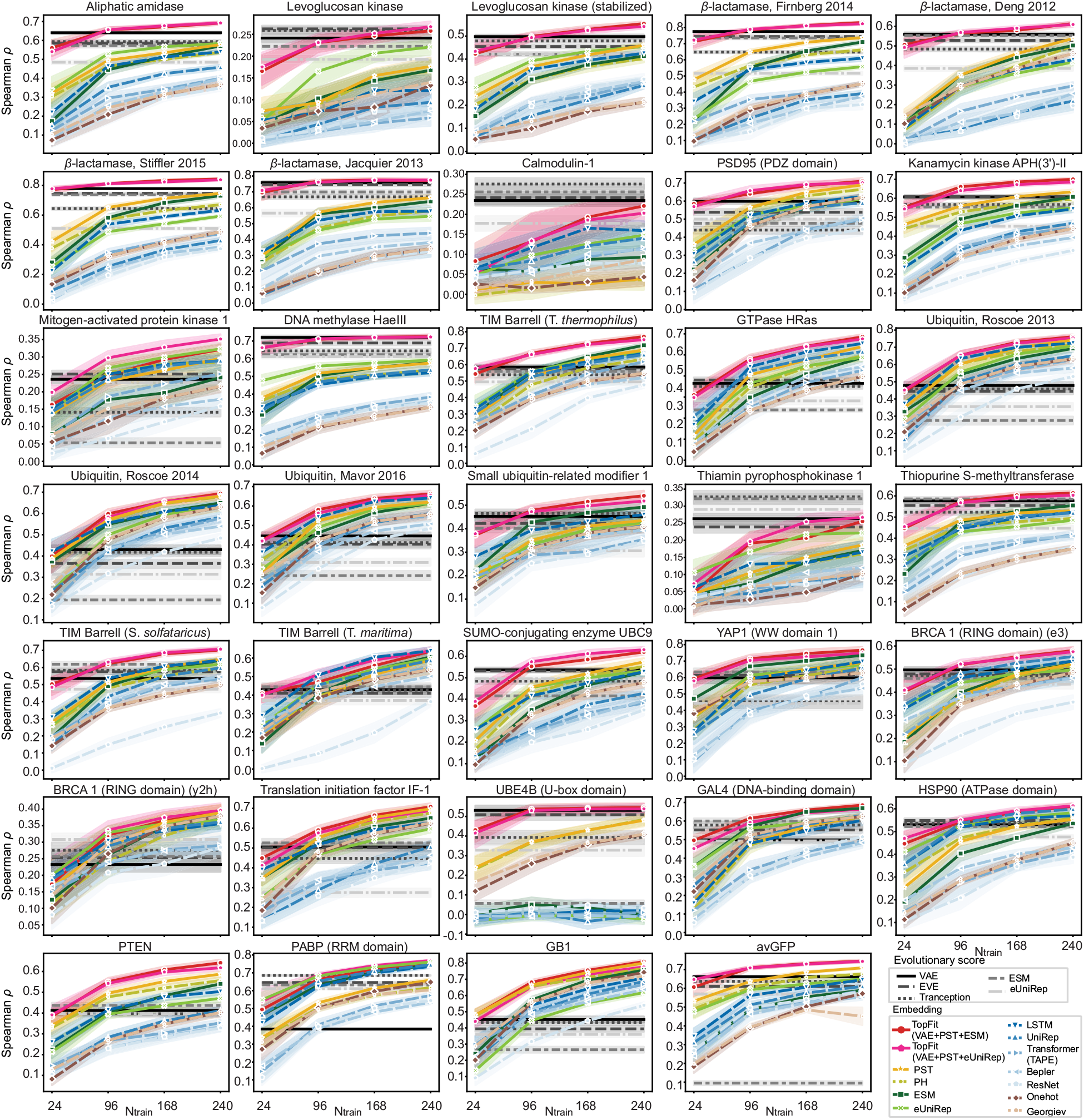
Spearman correlation for various models on individual datasets. It is the supplement of Figure 3**a**. Each line plots show average ρ for each dataset over n = 20 independent repeats. Training data sizes are 24, 96, 168, and 240. The width of the shade shows 95% confidence interval.

**Extended Data Figure 3.**
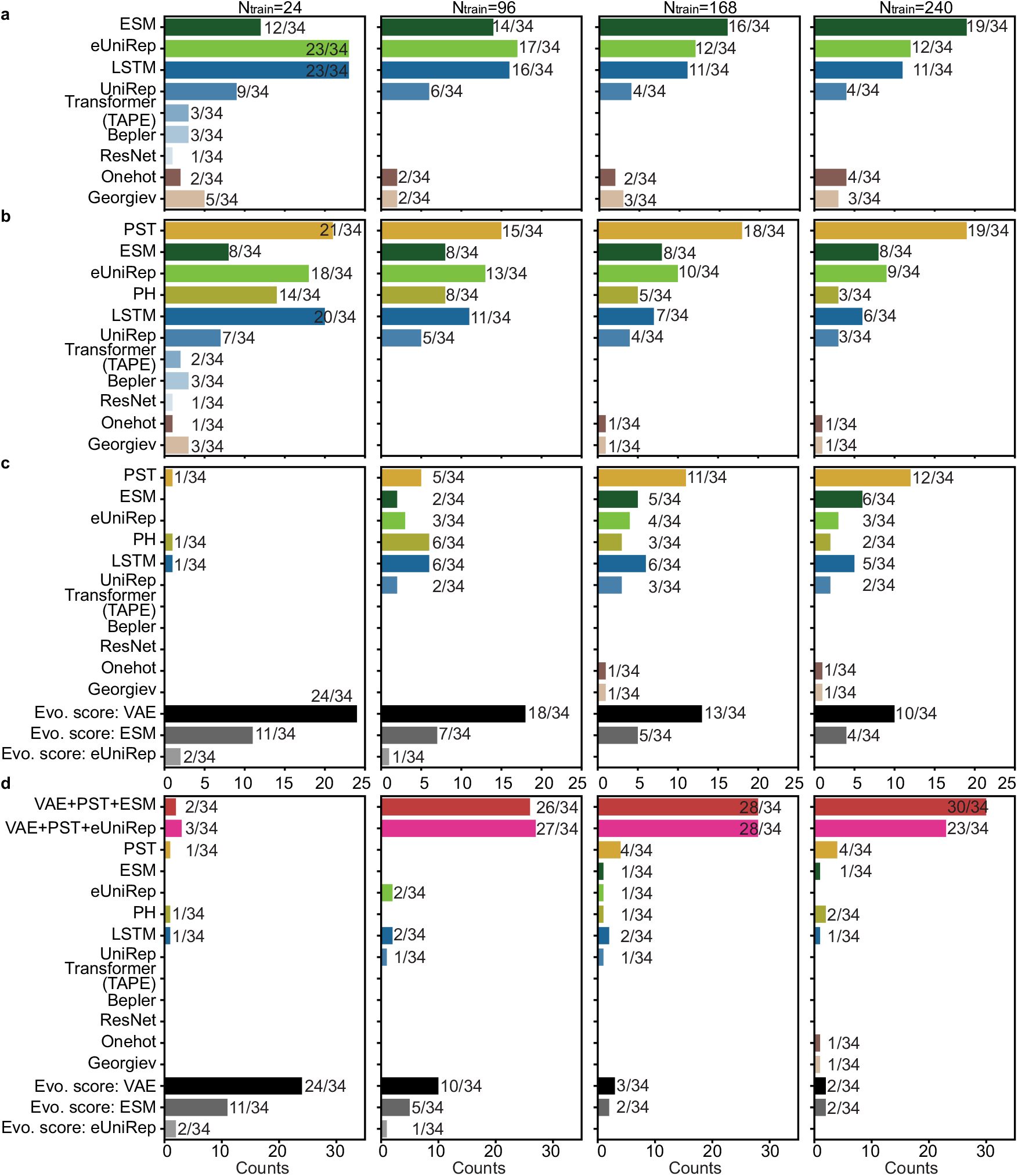
The frequency that an embedding is ranked as the best across 34 datasets using Spearman correlation. It is the supplement of Figure 3**b** including comparisons over different strategies. Histograms show the frequency that an embedding is ranked as the best across 34 datasets with 24, 96, 168, and 240 training data, respectively. For each dataset, the best embedding has average ρ over 20 independent repeats within the 95% confidence interval of the embedding with the highest average ρ. Comparisons were performed for **(a).** sequencebased embeddings; **(b).** structure- and sequence-based embeddings; **(c).** structure-based embeddings, sequence-based embeddings, and evolutionary scores; and **(d).** structure-based embeddings, sequence-based embeddings, evolutionary scores, and two sets of TopFit (VAE+PST+ESM and VAE+PST+eUniRep). We showed and used absolute values Spearman correlation for evolutionary scores.

**Extended Data Figure 4.**
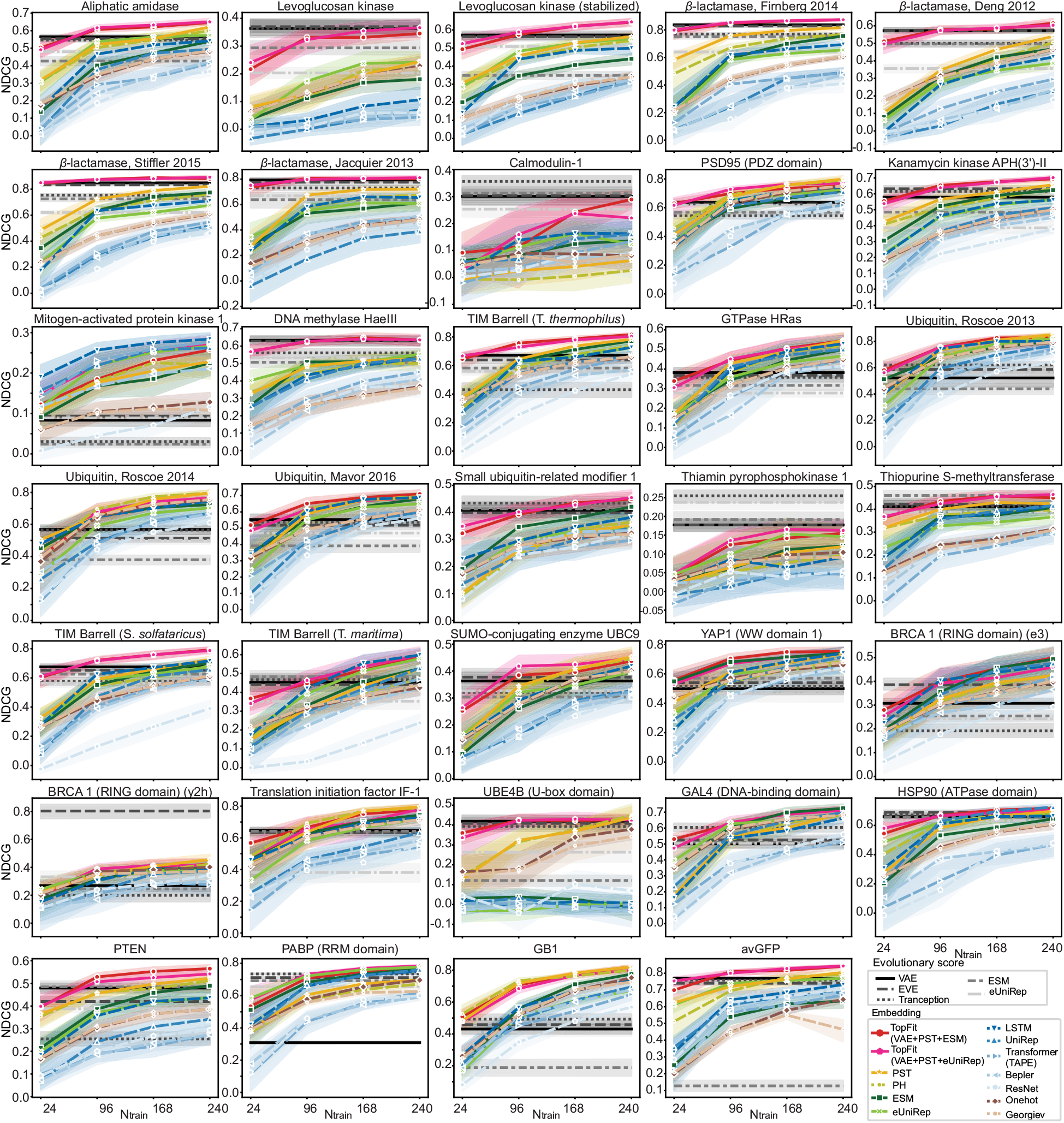
NDCG for various models on individual datasets. Analog of Extended Data Figure 2 but using NDCG. Each line plots show average NDCG for each dataset over n = 20 independent repeats. Training data sizes are 24, 96, 168, and 240. The width of the shade shows 95% confidence interval.

**Extended Data Figure 5.**
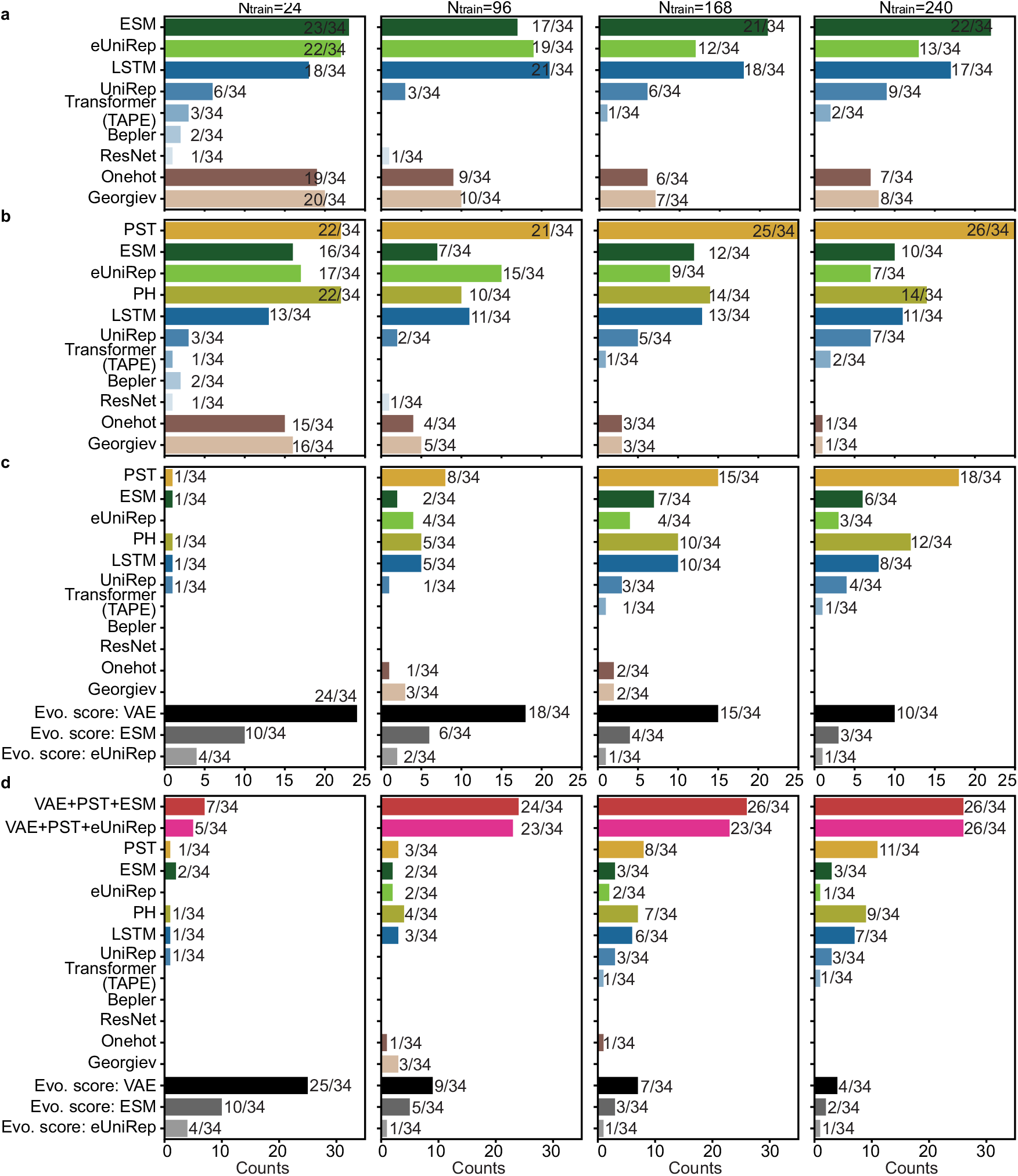
The frequency that an embedding is ranked as the best across 34 datasets using NDCG. Analog of Extended Data Figure 3 but measured by NDCG. Histograms show the frequency that an embedding is ranked as the best across 34 datasets with 24, 96, 168, and 240 training data, respectively. For each dataset, the best embedding has average NDCG over 20 independent repeats within the 95% confidence interval of the embedding with the highest average NDCG. Comparisons were performed for **(a).** sequence-based embeddings; **(b).** structure- and sequence-based embeddings; **(c).** structure-based embeddings, sequence-based embeddings, and evolutionary scores; and **(d).** structure-based embeddings, sequence-based embeddings, evolutionary scores, and two sets of TopFit (VAE+PST+ESM and VAE+PST+eUniRep). We showed and used absolute values NDCG for evolutionary scores.

**Extended Data Figure 6.**
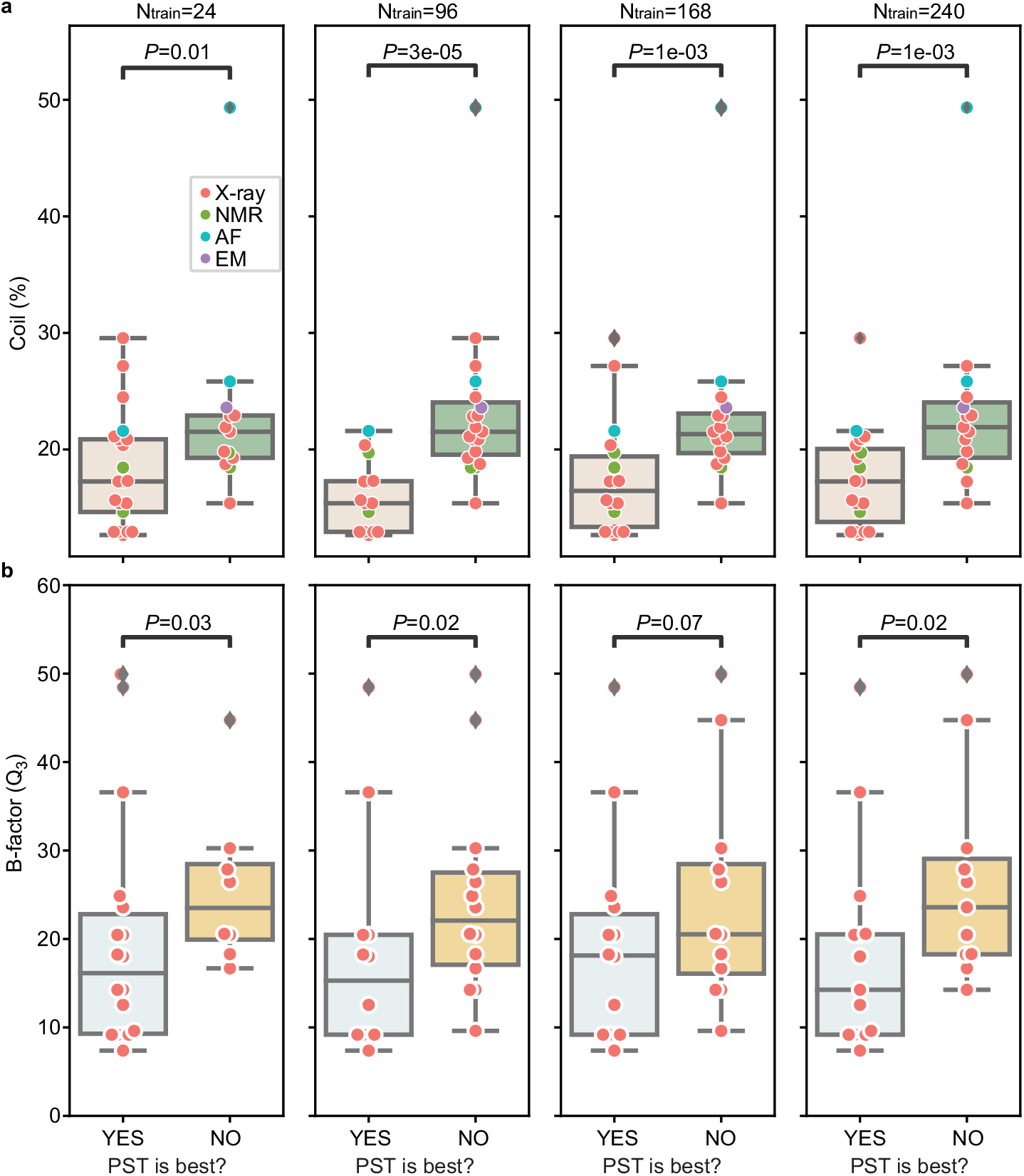
Relationships between quality of wild-type protein structure and PST performance. It is the supplement of Figure 3**c**. Boxplots show distribution of **(a)**. percentages of coils for protein structure over 34 datasets and **(b)**. third quartile (Q_3_) of B-factors at alpha carbons over 26 X-ray datasets. Datasets were classified into two classes depending on whether PST embedding is the best embedding. Scatter plots show same data with boxplots but for individual datasets. One-sided Mann-Whitney U test examines the statistical significance that two classes have different values. Boxplots display five-number summary where center line shows median, upper and lower limits of the box show upper and lower quartiles, and upper and lower whiskers show the maximum and the minimum by excluding “outliers” outside the interquartile range. In **(a)**, sample sizes for PST ranked as the best model are n = 21, n = 15, n = 18, and n = 19 for training data size 24, 96, 168, and 240, respectively. Sample sizes for PST not ranked as the best model are n = 13, n = 19, n = 16, and n = 15 for training data size 24, 96, 168, and 240, respectively. The p-values are 0.01, 3 ×10^−5^, 1 ×10^−3^, and 1 ×10^−3^ for training data size 24, 96, 168, and 240, respectively. In **(b)**, sample sizes for PST ranked as the best model are n = 8, n = 12, n = 14, and n = 15 for training data size 24, 96, 168, and 240, respectively. Sample sizes for PST not ranked as the best model are n = 18, n = 14, n = 12, and n = 11 for training data size 24, 96, 168, and 240, respectively. The p-values are 0.03, 0.02, 0.07, and 0.02 for training data size 24, 96, 168, and 240, respectively.

**Extended Data Figure 7.**
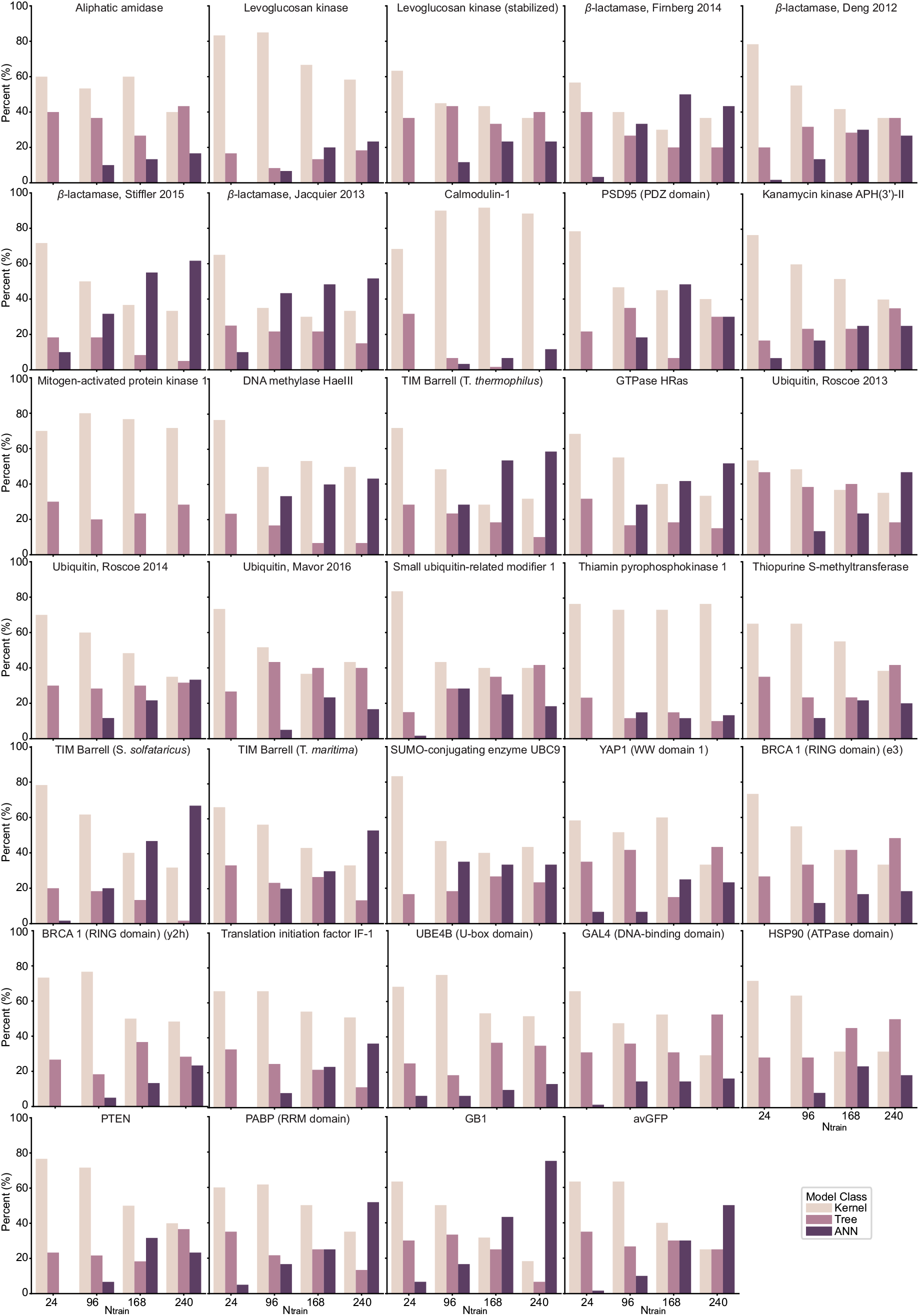
Model occurrence in ensemble regression. It is the supplement to Figure 3**e** to show model occurrence on individual datasets. For each repeat, the top N = 3 *regressors were picked and counted. Histograms count the model occurrence over* 20 *independent repeats*.

**Extended Data Figure 8.**
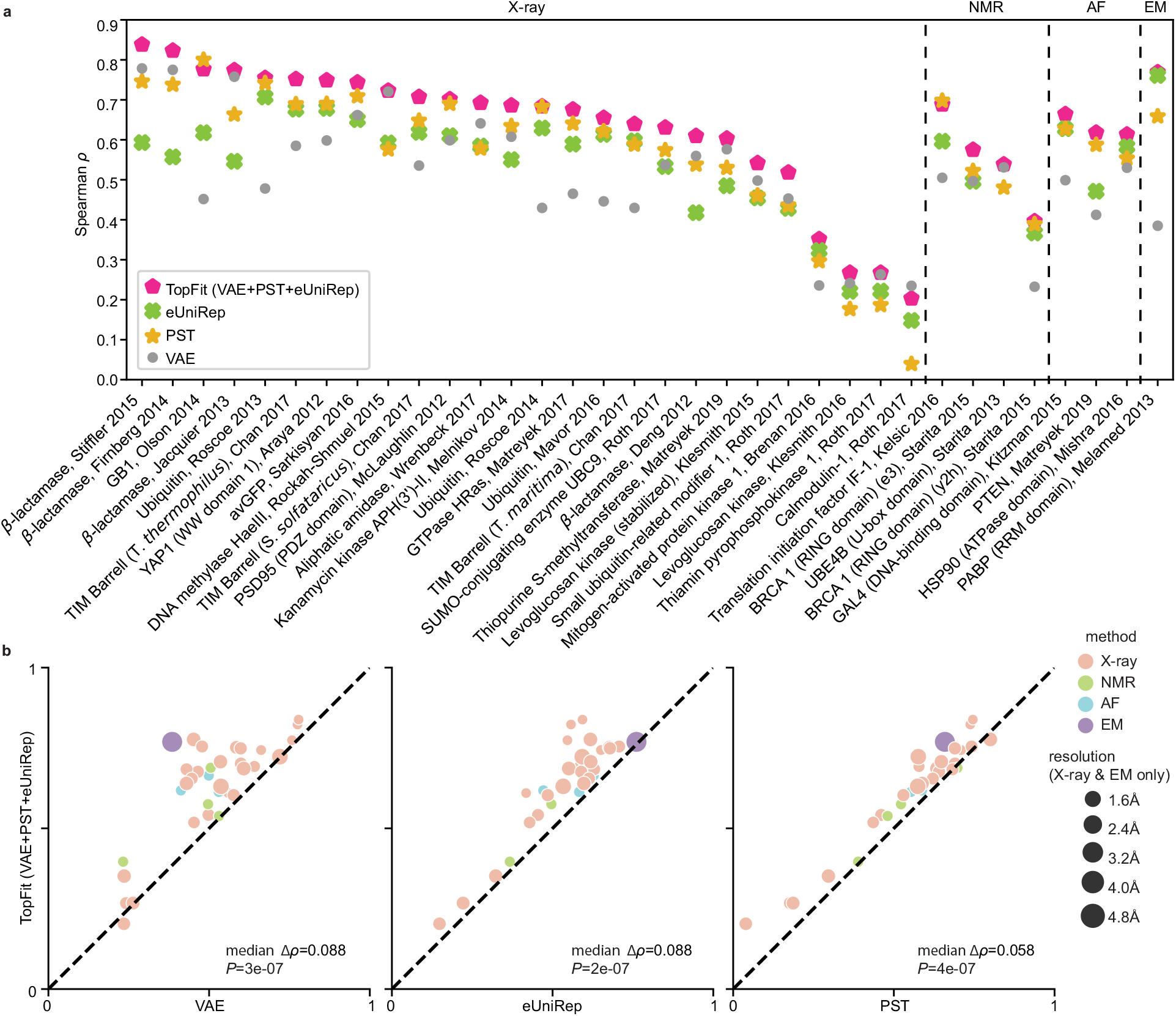
Comparisons between TopFit and other methods for mutation effects prediction using Spearman correlation. Analog to Figure 4**a-b**, but TopFit combines VAE score, eUniRep embedding, and PST embedding. All supervised models use 240 labeled training data. Results are evaluated by Spearman correlation ρ. DeepSequence VAE takes the absolute value of ρ. The average ρ from 20 independent repeats is shown. All 34 datasets are categorized by their structure modality used: X-ray, nuclear magnetic resonance (NMR), AlphaFold (AF), and cryogenic electron microscopy (EM). **(a)**. Dot plots show results across 34 datasets **(b).** Dot plots show pairwise comparison between TopFit with one method at each plot. Medians of difference for average Spearman correlation Δ ρ across all datasets are shown. One-sided rank-sum test determines the statistical significance that TopFit has better performance than VAE score, eUniRep embedding and PST embedding with p-values P = 3 ×10^−7^, 2 × 10^−7^, and 4 × 10^−7^, respectively.

**Extended Data Figure 9.**
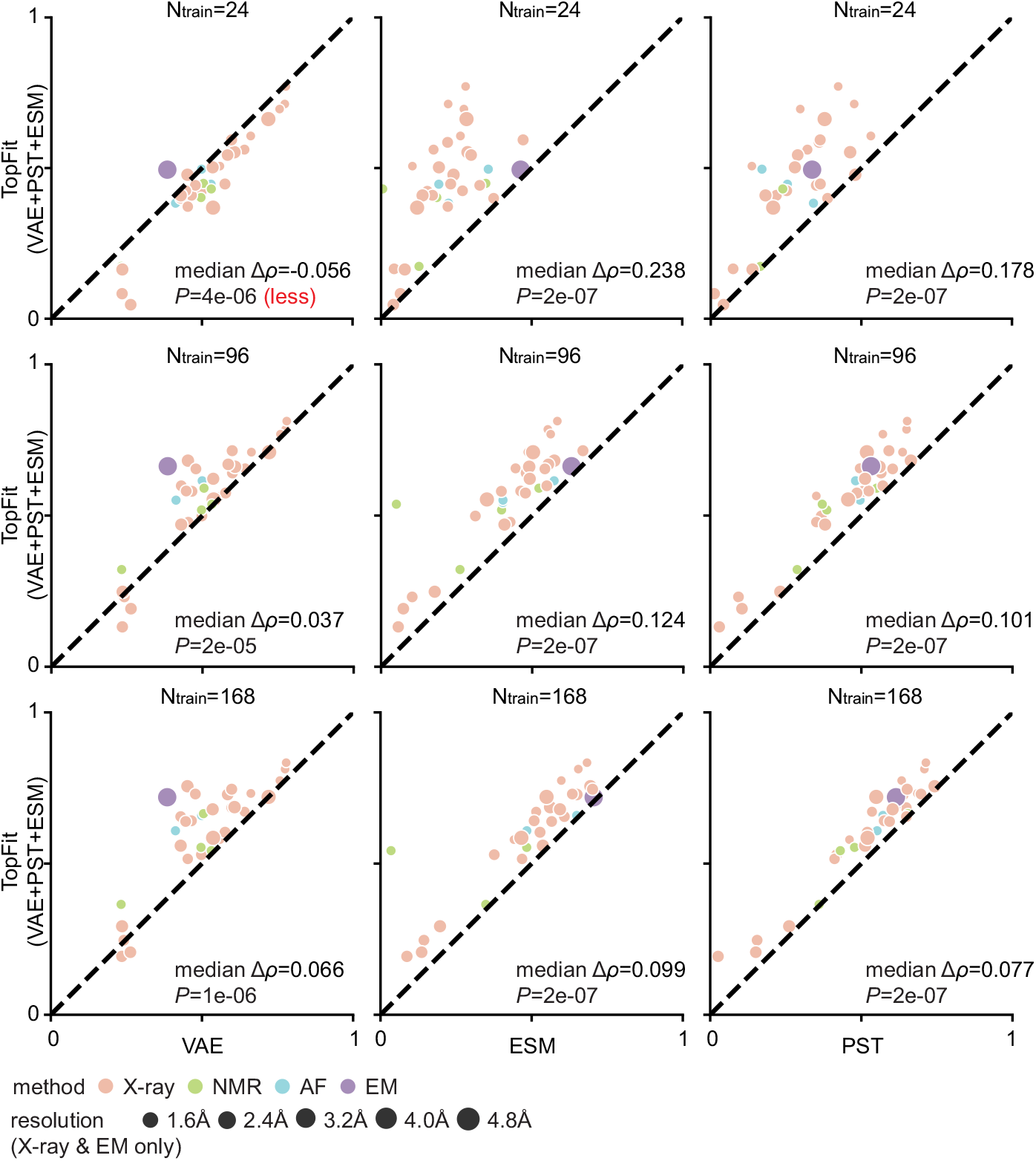
Comparisons between TopFit with other methods. TopFit consists of VAE score, ESM embedding, and PST embedding. It is supplement of Figure 4**b** to include results with various numbers of training data. Average Spearman correlation from 20 independent repeats are shown, and all datasets are categorized by their structure modality used: X-ray, nuclear magnetic resonance (NMR), AlphaFold (AF), and cryogenic electron microscopy (EM). One-sided rank-sum test determines the statistical significance that TopFit has better performance than other strategies, except we use null hypothesis that TopFit has worse performance than VAE with 24 training data. The p-values are shown in the corresponding subfigures. They are 1. TopFit versus VAE: P = 4 × 10^−6^, 2 ×10^−5^, and 1 × 10^−6^; 2. TopFit versus ESM: P = 2 × 10^−7^, 2 ×10^−7^, and 2 × 10^−7^; and 3. TopFit versus PST: P = 2 × 10^−7^, 2 × 10^−7^, and 2 × 10^−7^for training data size 24, 96, and 168, respectively.

**Extended Data Figure 10.**
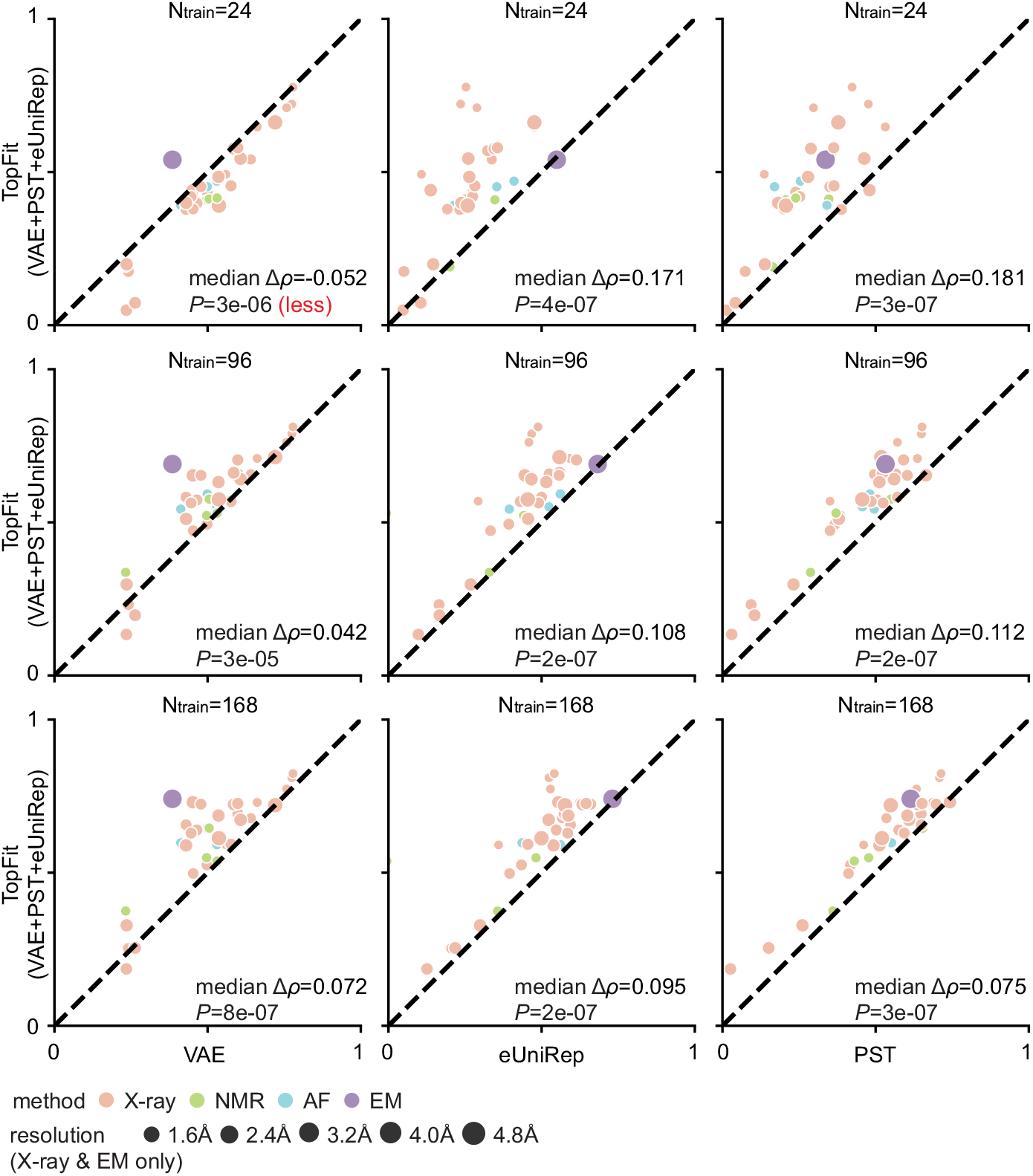
Comparisons between TopFit with other methods. TopFit consists of VAE score, eUniRep embedding, and PST embedding. It is the supplement of Extended Data Figure 8**b** to include results with various numbers of training data. Average Spearman correlation from 20 independent repeats are shown, and all datasets are categorized by their structure modality used: X-ray, nuclear magnetic resonance (NMR), AlphaFold (AF), and cryogenic electron microscopy (EM). One-sided rank-sum test determines the statistical significance that TopFit has better performance than other strategies, except we use null hypothesis that TopFit has worse performance than VAE with 24 training data. The p-values are shown in the corresponding subfigures. They are 1. TopFit versus VAE: P = 3 × 10^−6^, 3 × 10^−5^, and 8 × 10^−7^; 2. TopFit versus eUniRep: P = 4 × 10^−7^, 2 × 10^−7^, and 2 × 10^−7^; and 3. TopFit versus PST: P = 3 × 10^−7^, 2 × 10^−7^, and 3 × 10^−7^for training data size 24, 96, and 168, respectively.

## Notes

### Competing Interest Statement

The authors have declared no competing interest.

