## Supplementary Information for "Persistent spectral theory-guided protein engineering"

**This file includes the following subsections:**

1. **Supplementary Figures:**


- Supplementary Figure 1: The average performance of various embeddings for datasets categorized by their taxonomy.
- Supplementary Figure 2: The average performance of various embeddings for datasets categorized by the fitness types.
- Supplementary Figure 3: The average performance of various embeddings using ensemble models from different categories.
- Supplementary Figure 4: Comparisons of regressors using Spearman correlation.
- Supplementary Figure 5: Comparisons for performance utilizing different types of structure modality evaluated by Spearman correlation.
- Supplementary Figure 6: The average performance of embeddings augmented with different evolutionary scores.
- Supplementary Figure 7: The average performance of TopFit augmented with single or multiple evolutionary scores.
- Supplementary Figure 8: Extrapolation task predicting multiple-mutation datasets from single-mutation data.
- Supplementary Figure 9: The average performance of various models over 27 single-mutation datasets for predicting unseen mutational sites in training sets.
- Supplementary Figure 10: Spearman correlation for various models on individual datasets for predicting unseen mutational sites in training sets.
- Supplementary Figure 11: The frequency that an embedding is ranked as the best across 27 single-mutation datasets for predicting unseen mutational sites in training sets, where performance is evaluated by Spearman correlation.
- Supplementary Figure 12: NDCG for various models on individual datasets for predicting unseen mutational sites in training sets.
- Supplementary Figure 13: The frequency that an embedding is ranked as the best across 27 single-mutation datasets for predicting unseen mutational sites in training sets, where performance is evaluated by NDCG.
- Supplementary Figure 14: Structure-based embeddings using persistent landscape representations.
- Supplementary Figure 15: PST embedding with additional non-harmonic persistent spectral features from $L_1$ and $L_2$.
- Supplementary Figure 16: TopFit embedding with additional non-harmonic persistent spectral features from $L_1$ and $L_2$.


2. **Supplementary Tables:**


- Supplementary Table 1: List of regressors used in ensemble regression.

- Supplementary Table 2: List of hyperparameters and their optimization ranges for kernel and tree models.
- Supplementary Table 3: List of hyperparameters and their optimization ranges for ANN models.
- Supplementary Table 4: Spearman correlation obtained from TopFit by combining PST, ESM embedding and VAE score.
- Supplementary Table 5: Spearman correlation from Augmented VAE model and ECNet model.

3. **Supplementary Notes:**

- Supplementary Note 1: Embedding comparisons
- Supplementary Note 2: Evolutionary scores for unsupervised fitness predictions
- Supplementary Note 3: TopFit augmented with different evolutionary scores
- Supplementary Note 4: Improvements of TopFit
- Supplementary Note 5: Predictions of unseen mutational sites
- Supplementary Note 6: Representations of persistent features
- Supplementary Note 7: Sources of datasets

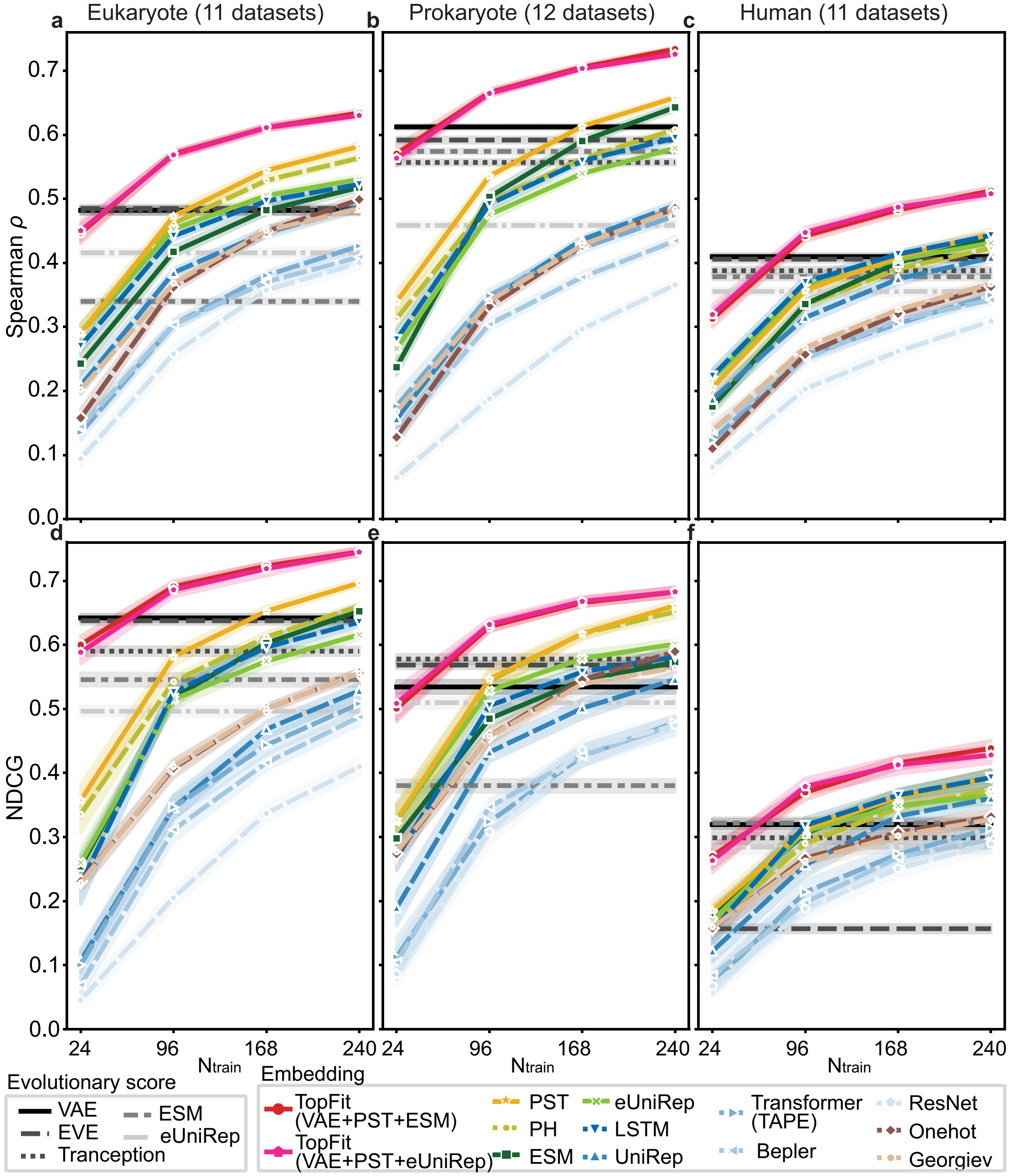


**Supplementary Figure 1: The average performance of various embeddings for datasets categorized by their taxonomy.** All 34 datasets are categorized by their taxonomy with 11 Eukaryote, 12 Prokaryote, and 11 Human datasets. The average Spearman correlations are shown for **(a).** Eukaryote, **(b).** Prokaryote, and **(c).** Human datasets. The average NDCGs are shown for **(d).** Eukaryote, **(e).** Prokaryote, and **(f).** Human datasets. The ridge regression is used for Georgiev and one-hot embedding. Otherwise, ensemble regression is used. The width of shade shows 95% confidence interval from $n = 20$ independent repeats. We showed and used absolute values of Spearman correlation and NDCG for evolutionary scores.

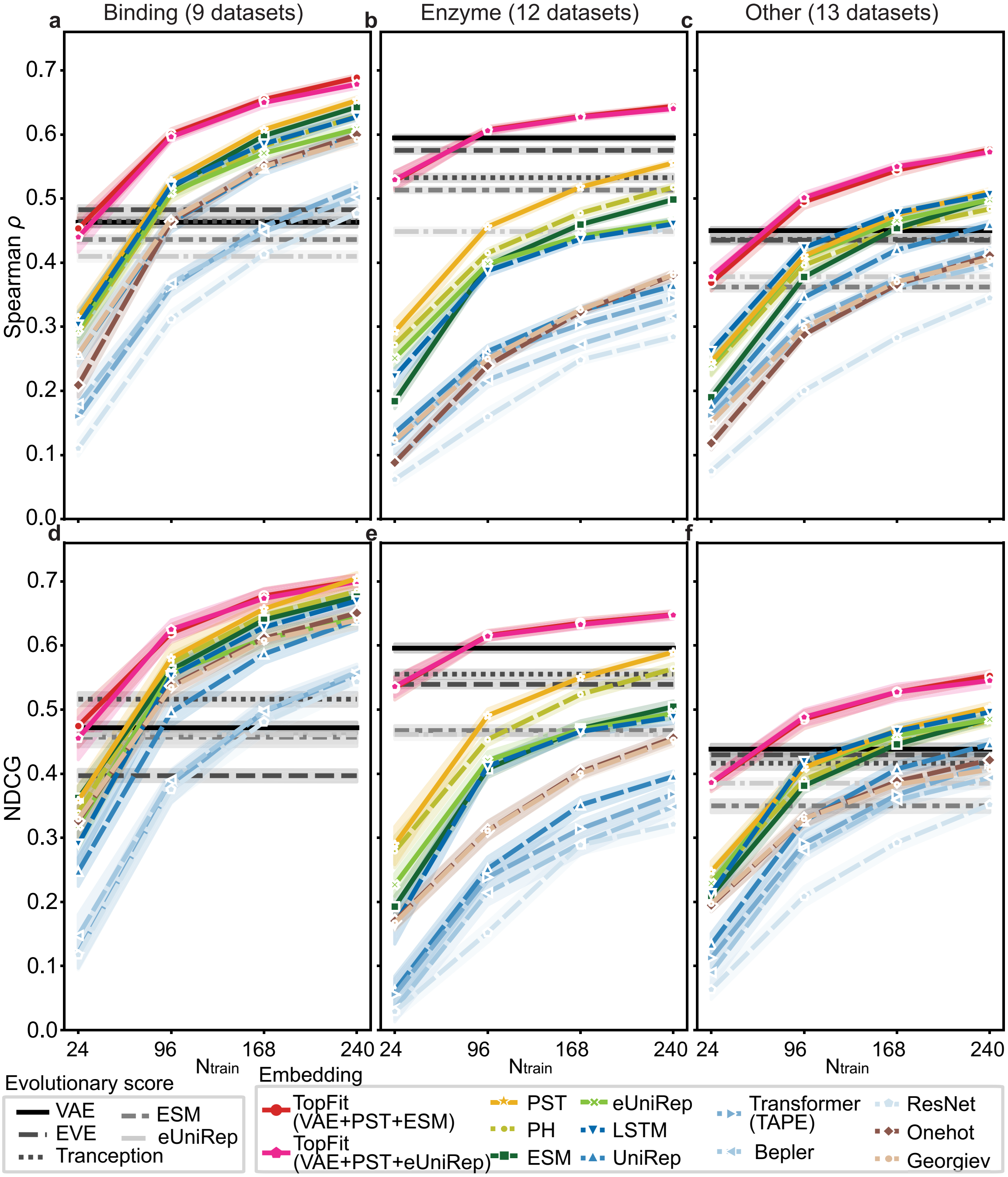


**Supplementary Figure 2: The average performance of various embeddings for datasets categorized by the fitness types.** All 34 datasets are categorized to three types by their types of fitness: 9 datasets for binding, 12 datasets for enzyme activity, and 13 datasets for other fitness. Others include various types of fitness such as protein stability, growth experiments, fluorescence, and drug and antibiotic resistance. The average Spearman correlations are shown for **(a).** binding, **(b).** enzyme activity, and **(c).** others. The average NDCGs are shown for **(d).** binding, **(e).** enzyme activity, and **(f).** other. The ridge regression is used for Georgiev and one-hot embedding. Otherwise, ensemble regression is used. The width of shade shows 95% confidence interval from $n = 20$ independent repeats. We showed and used absolute values of Spearman correlation and NDCG for evolutionary scores.

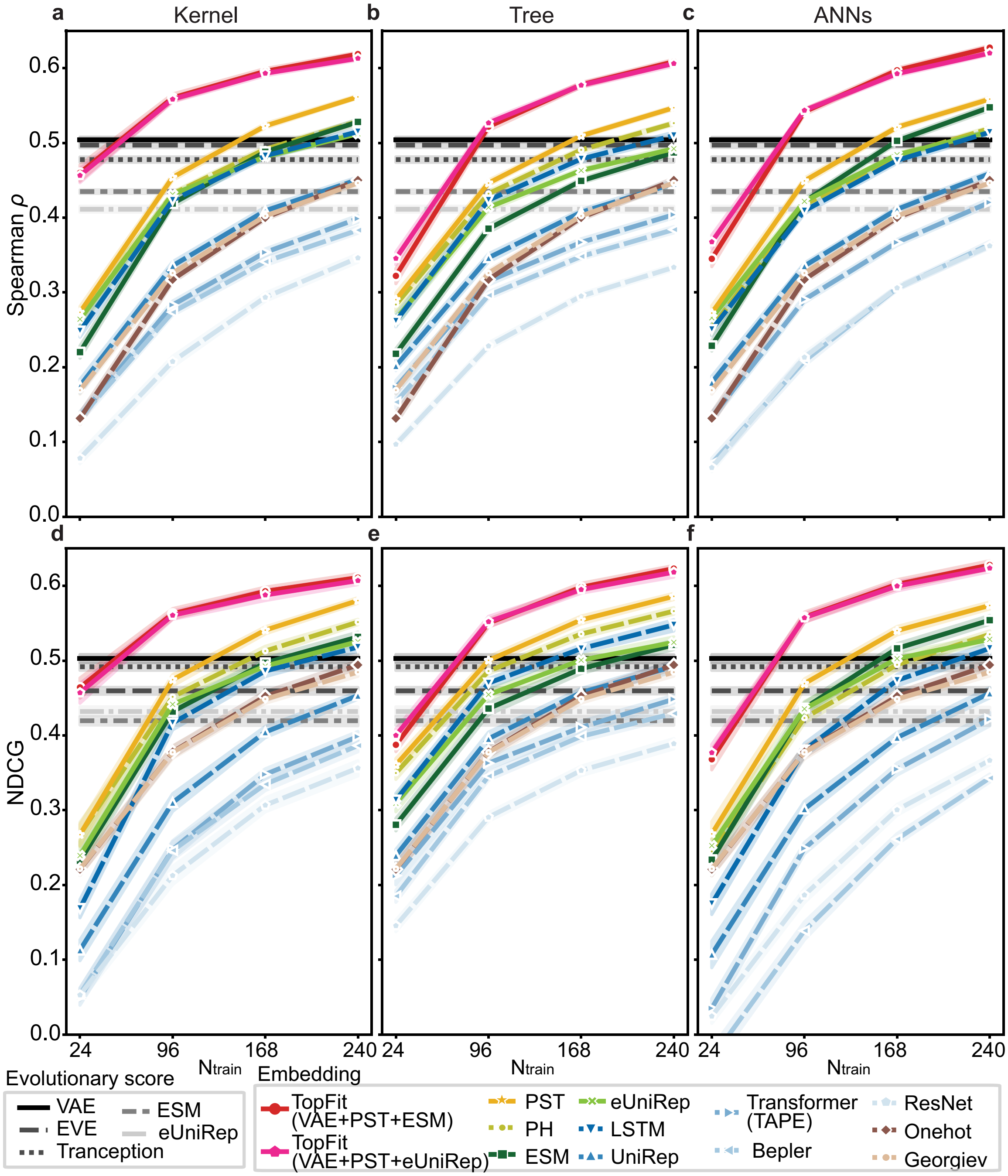


**Supplementary Figure 3: The average performance of various embeddings using ensemble models from different categories.** Different embeddings are tested on three categories of ensemble regression. The ensembles for kernel, tree, and ANN models contain 10, 5, and 3 regressors, respectively. The average performance over 34 datasets is shown. The average Spearman correlations are shown for ensemble of **(a).** kernel, **(b).** tree, and **(c).** ANN models. The average NDCGs are shown for **(d).** kernel, **(e).** tree, and **(f).** ANN models. The ridge regression is used for Georgiev and one-hot embedding for baseline comparisons. The width of shade shows 95% confidence interval from $n = 20$ independent repeats. We showed and used absolute values of Spearman correlation and NDCG for evolutionary scores.

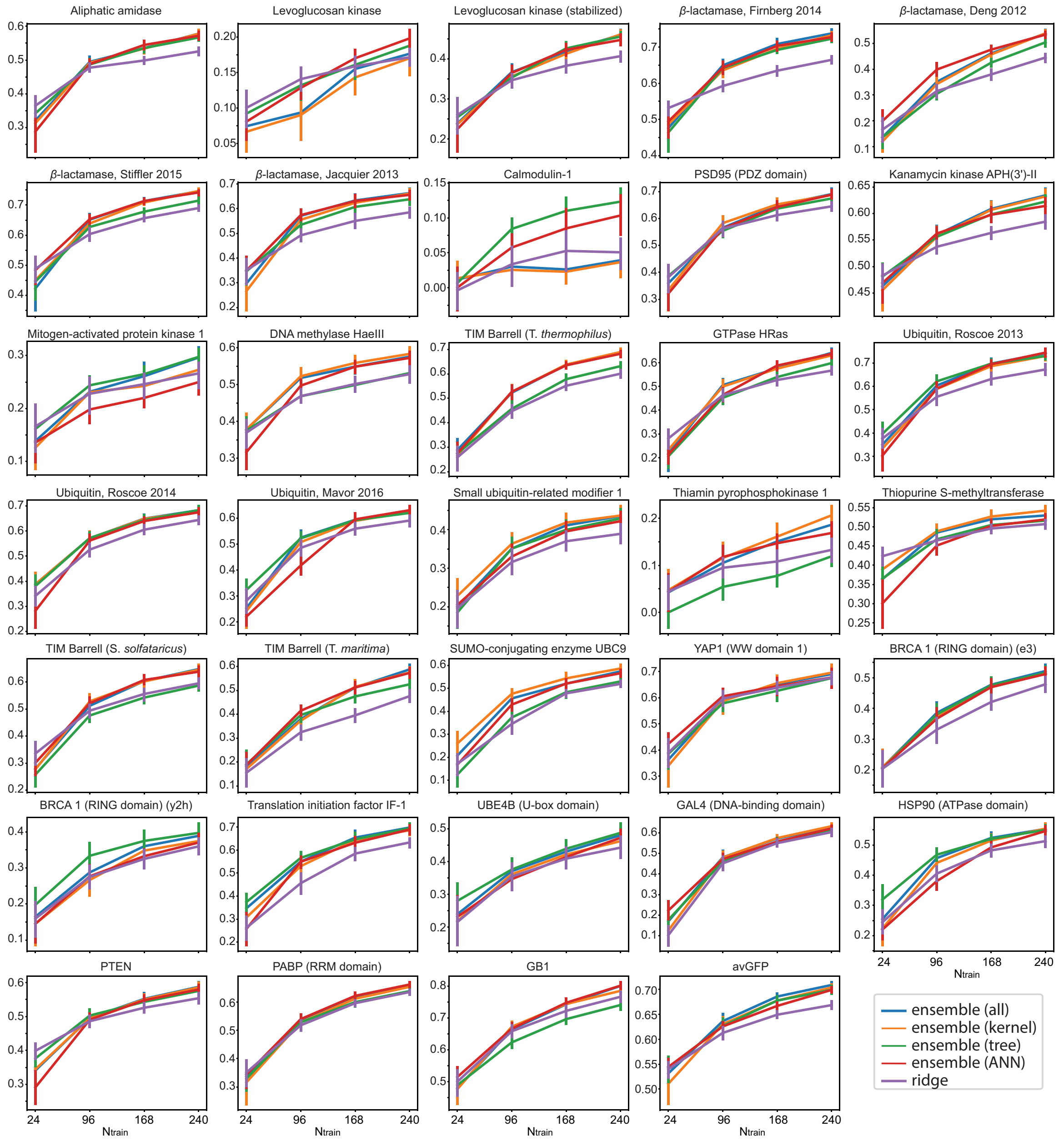


**Supplementary Figure 4: Comparisons of regressors using Spearman correlation.** It is the supplement to Figure 3**d** to show performance on individual dataset. The ensemble model for all regressors is compared with ensemble models for single type of regressors in kernels, trees, and ANNs. The ridge regression is also compared. The model uses the PST embedding as features. Error bars show 95% confidence interval from $n = 20$ independent repeats.

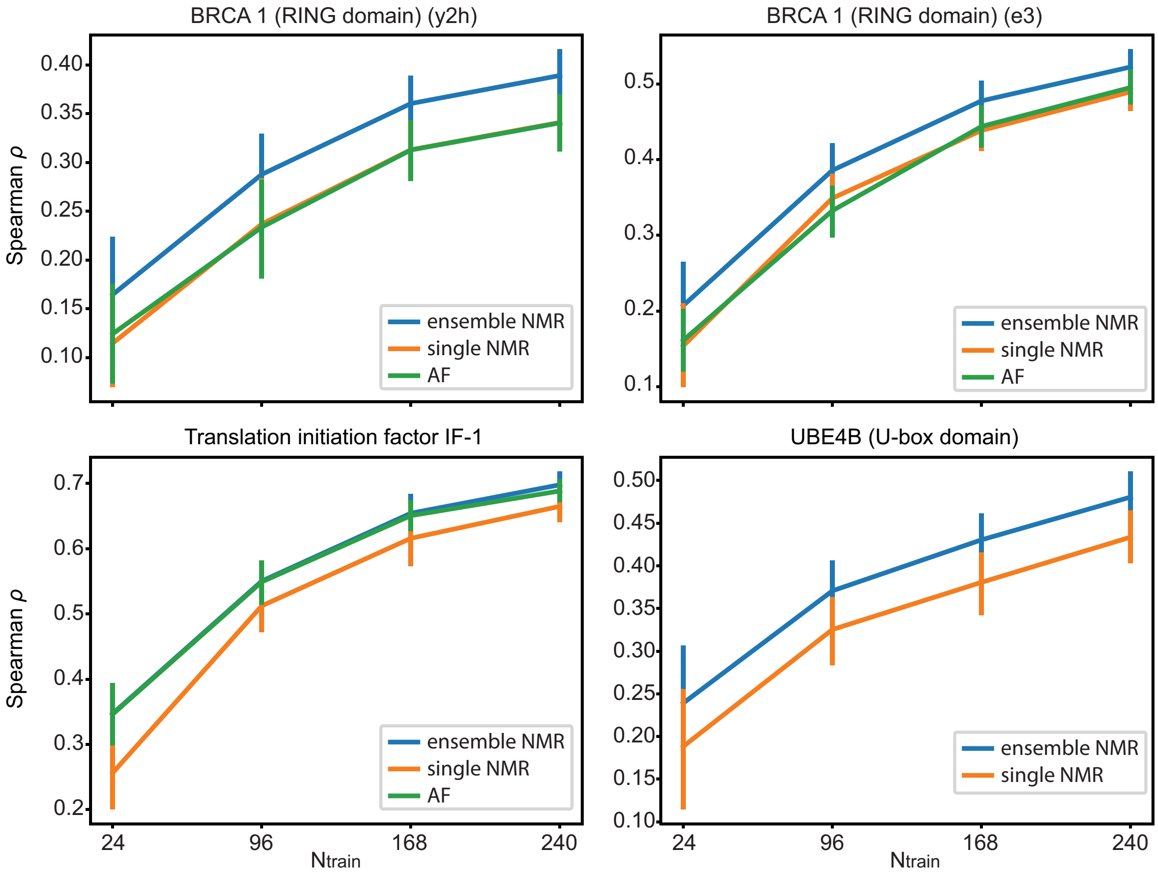


**Supplementary Figure 5: Comparisons for performance utilizing different types of structure modality evaluated by Spearman correlation.** Nuclear magnetic resonance (NMR) and AlphaFold (AF) are used for comparisons. Since one NMR data contains multiple structure models in one experiment, "Ensemble NMR" indicates average predictions from all NMR models. "Single NMR" indicates predictions from the first NMR model. The PST embedding is used. Error bars show 95% confidence interval from $n = 20$ independent repeats.

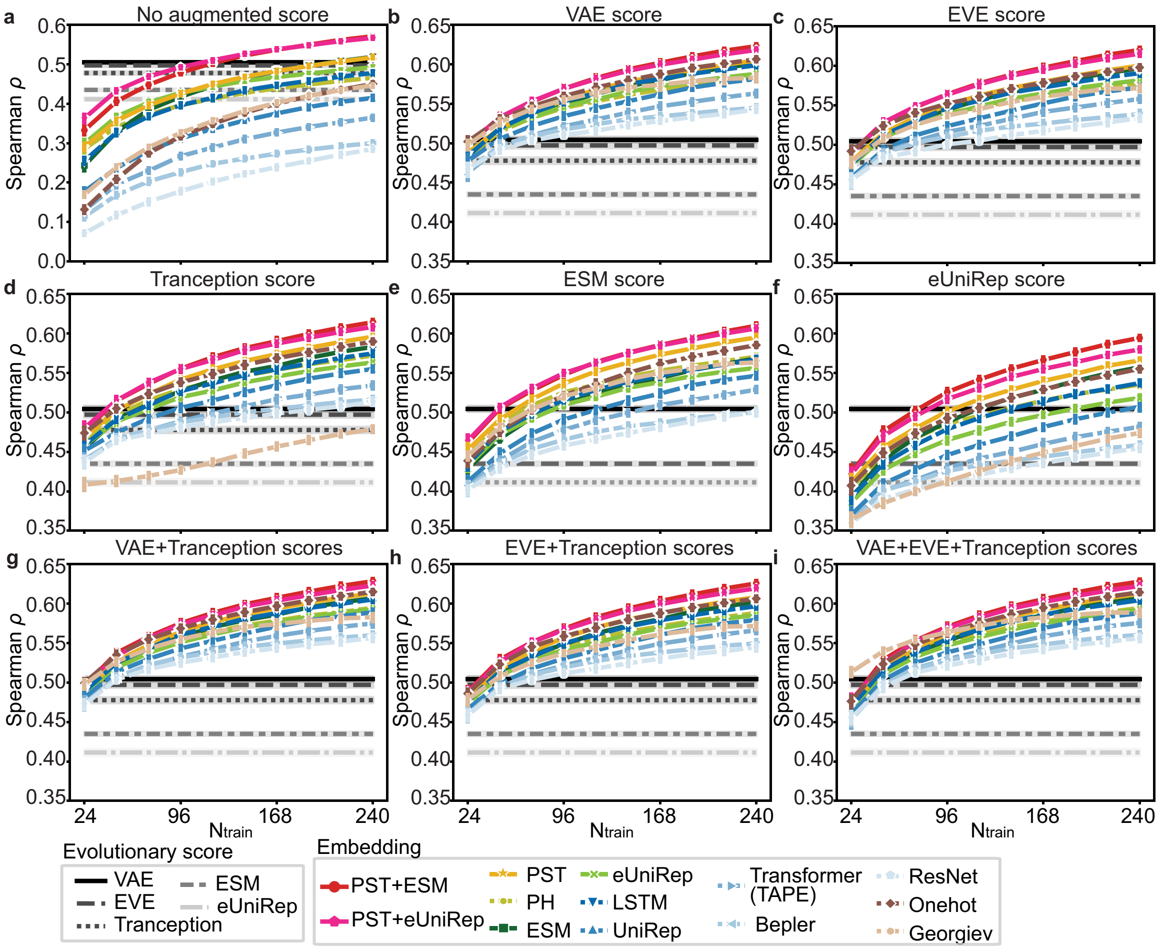


**Supplementary Figure 6: The average performance of embeddings augmented with different evolutionary scores.** Embeddings are augmented single score: **(a).** no evolutionary score; **(b).** DeepSequence VAE score; **(c).** EVE score; **(d).** Tranception score; **(e).** ESM score; **(f).** eUniRep score, and combinations of two scores: **(g).** DeepSequence VAE and Tranception scores; **(h).** EVE and Tranception scores and combinations of three scores; and **(i).** DeepSequence VAE, EVE and Tranception scores. Spearman correlation is used, and the results show average performance across 34 datasets. Ridge regression is used. The width of shade shows 95% confidence interval from $n = 20$ independent repeats.

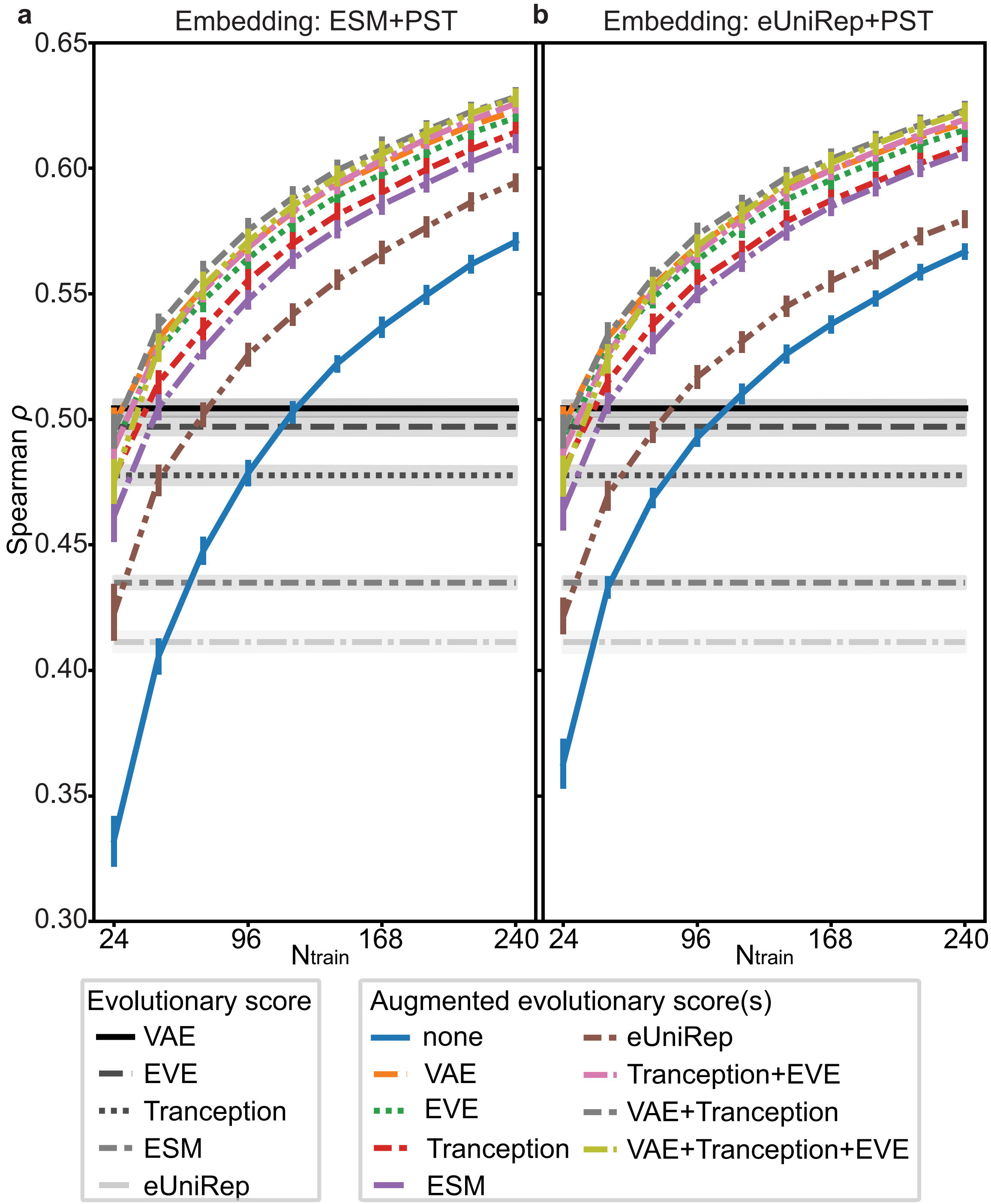


**Supplementary Figure 7: The average performance of TopFit augmented with single or multiple evolutionary scores.** The embeddings for **(a).** ESM+PST and **(b).** eUniRep+PST are augmented with various evolutionary scores. The augmentations are done with five single evolutionary scores: VAE, EVE, Tranception, ESM, and eUniRep. Additional augmentations are carried out with three combinations of two or three scores, namely, Tranception+EVE, VAE+Tranception, and VAE+Tranception+EVE. Spearman correlation is used, and the results show average performance across 34 datasets. Ridge regression is used. The width of shade shows 95% confidence interval from $n = 20$ independent repeats.

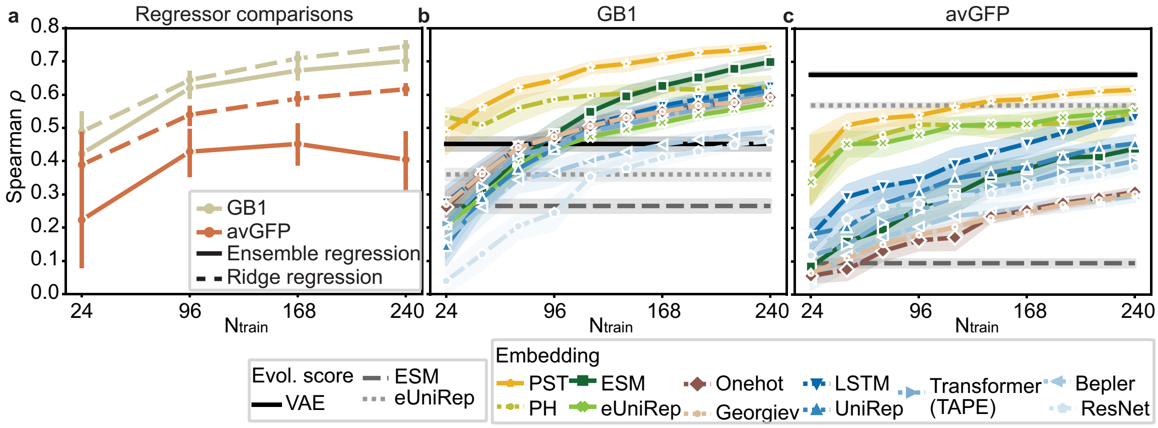


**Supplementary Figure 8: Extrapolation task predicting multiple-mutation datasets from single-mutation data.** **(a)** Comparisons between ridge regression and ensemble regression for extrapolation task. The performance of PST embedding using ridge regression for various sizes of training data for **(b).** GB1 dataset and **(c).** avGFP dataset. In **(a)-(c)**, Spearman correlation is used as the evaluation metric. The width of shade or error bars show 95% confidence interval from $n = 20$ independent repeats.

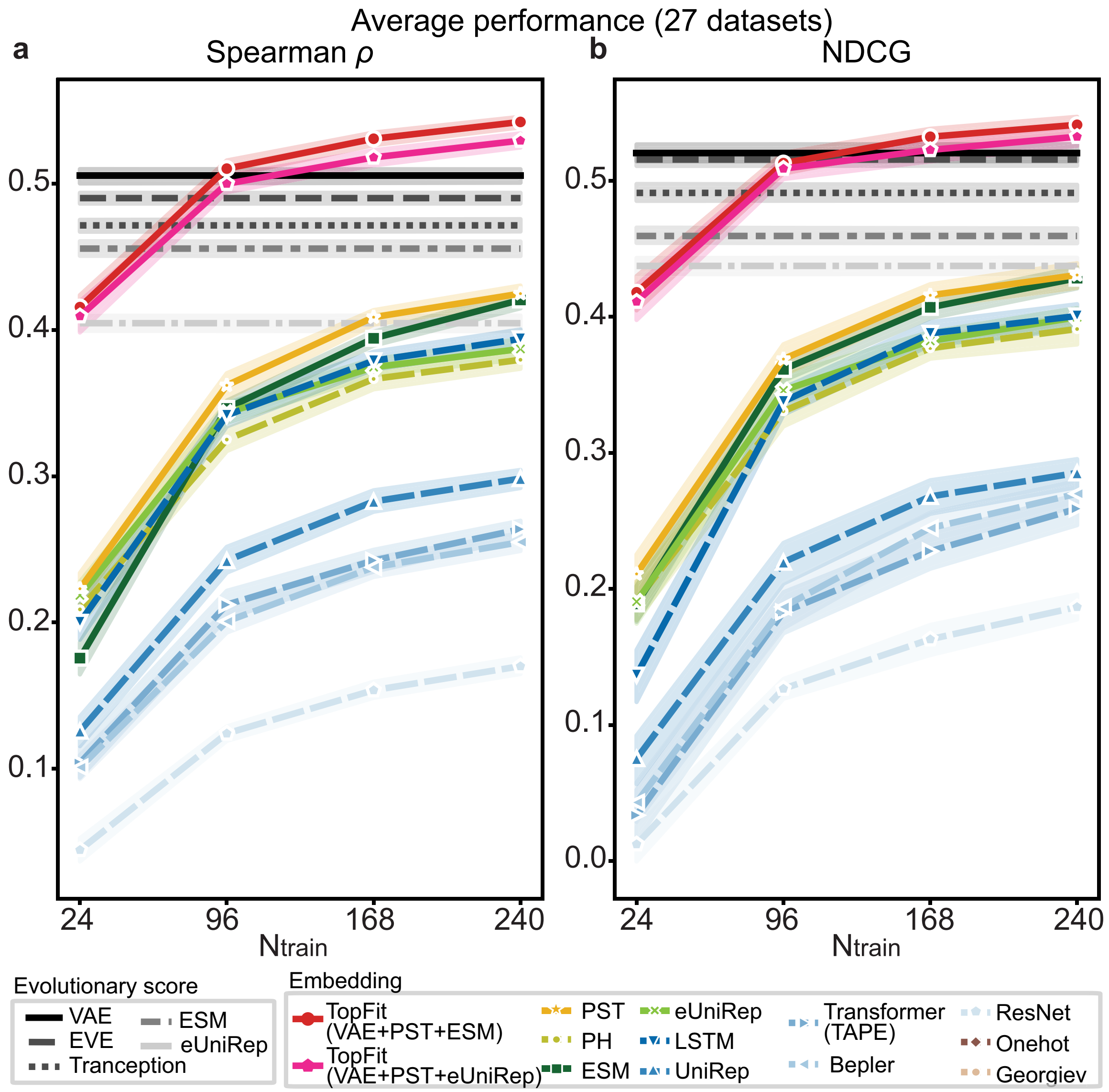


**Supplementary Figure 9: The average performance of various models over 27 single-mutation datasets for predicting unseen mutational sites in training sets.** Results are evaluated by **(a).** Spearman correlation and **(b).** NDCG. Ensemble regression for 18 regressors is used. The width of shade shows 95% confidence interval from $n = 20$ independent repeats. We showed and used absolute values of **(a).** Spearman correlation or **(b).** NDCG for evolutionary scores.

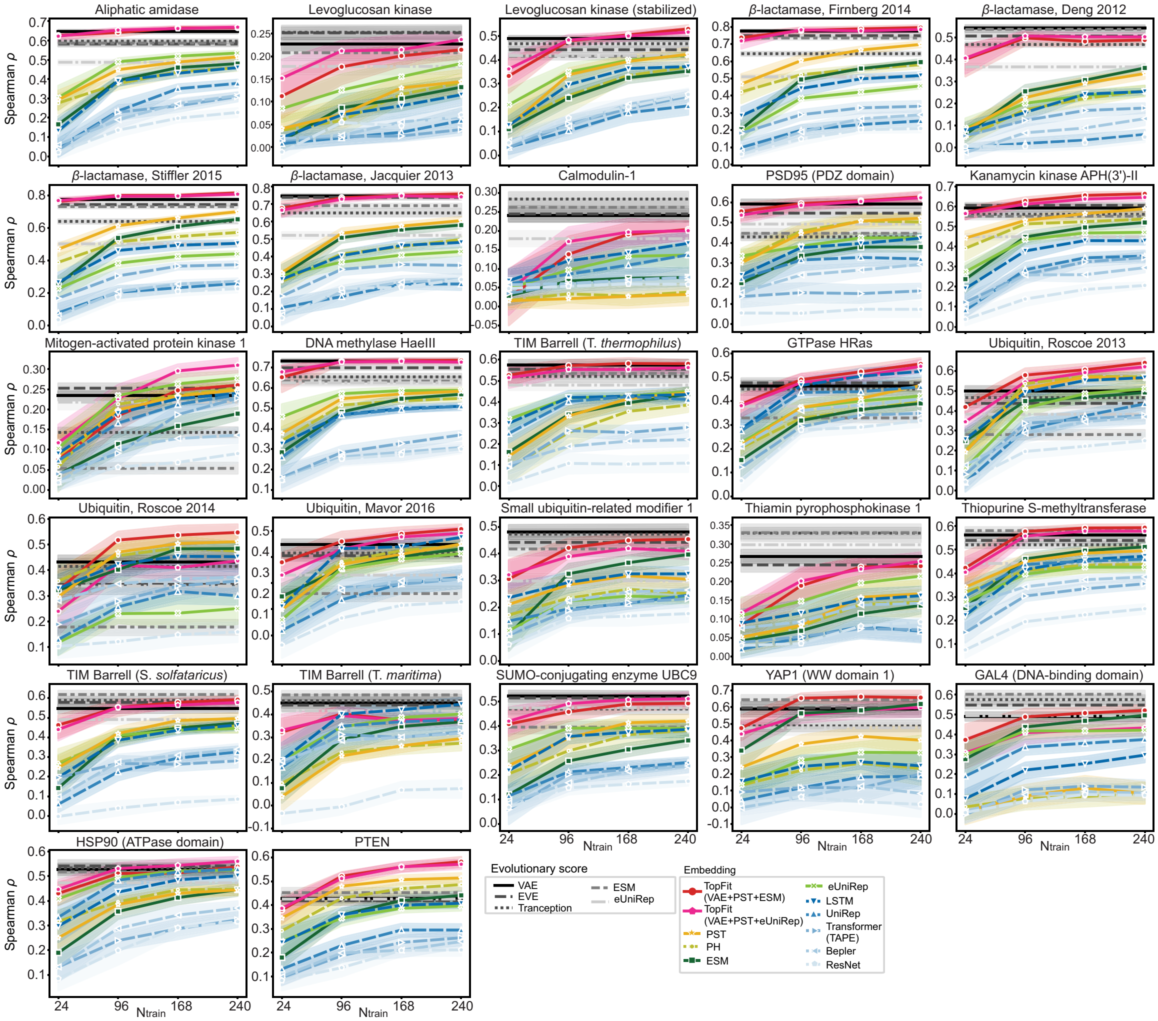


**Supplementary Figure 10: Spearman correlation for various models on individual datasets for predicting unseen mutational sites in training sets.** The ensemble regression is used. The results were shown for training data sizes 24, 96, 168, and 240 from $n = 20$ independent repeats. The width of the shade shows 95% confidence interval. We showed and used absolute values of Spearman correlations for evolutionary scores.

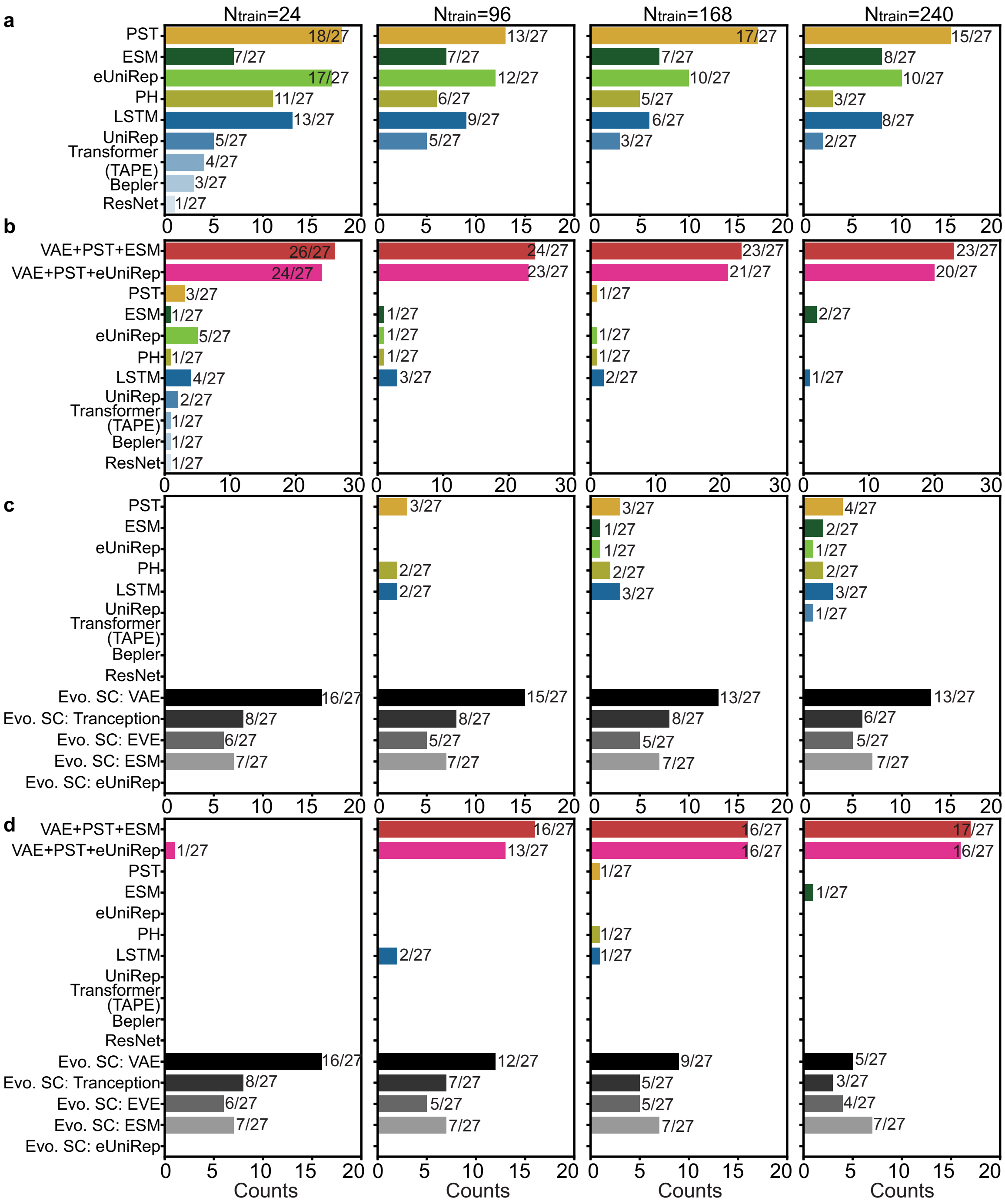


**Supplementary Figure 11: The frequency that an embedding is ranked as the best across 27 single-mutation datasets for predicting unseen mutational sites in training sets, where performance is evaluated by Spearman correlation.** For each dataset, the embedding with the highest average $\rho$ is identified as the top embedding. The best embedding includes the top embedding and others within the 95% confidence interval of the top embedding. Comparisons for **(a).** structure-based embeddings, sequence-based embeddings, and two sets of TopFit (VAE+PST+ESM and VAE+PST+eUniRep) **(b).** TopFit, structure-based and sequence-based embeddings; **(c).** structure-based embeddings, sequence-based embeddings, and evolutionary scores; **(d).** structure-based embeddings, sequence-based embeddings, evolutionary scores, and two sets of TopFit (VAE+PST+ESM and VAE+PST+eUniRep). We showed and used absolute values Spearman correlation for evolutionary scores. Each embedding has 20 independent repeats.

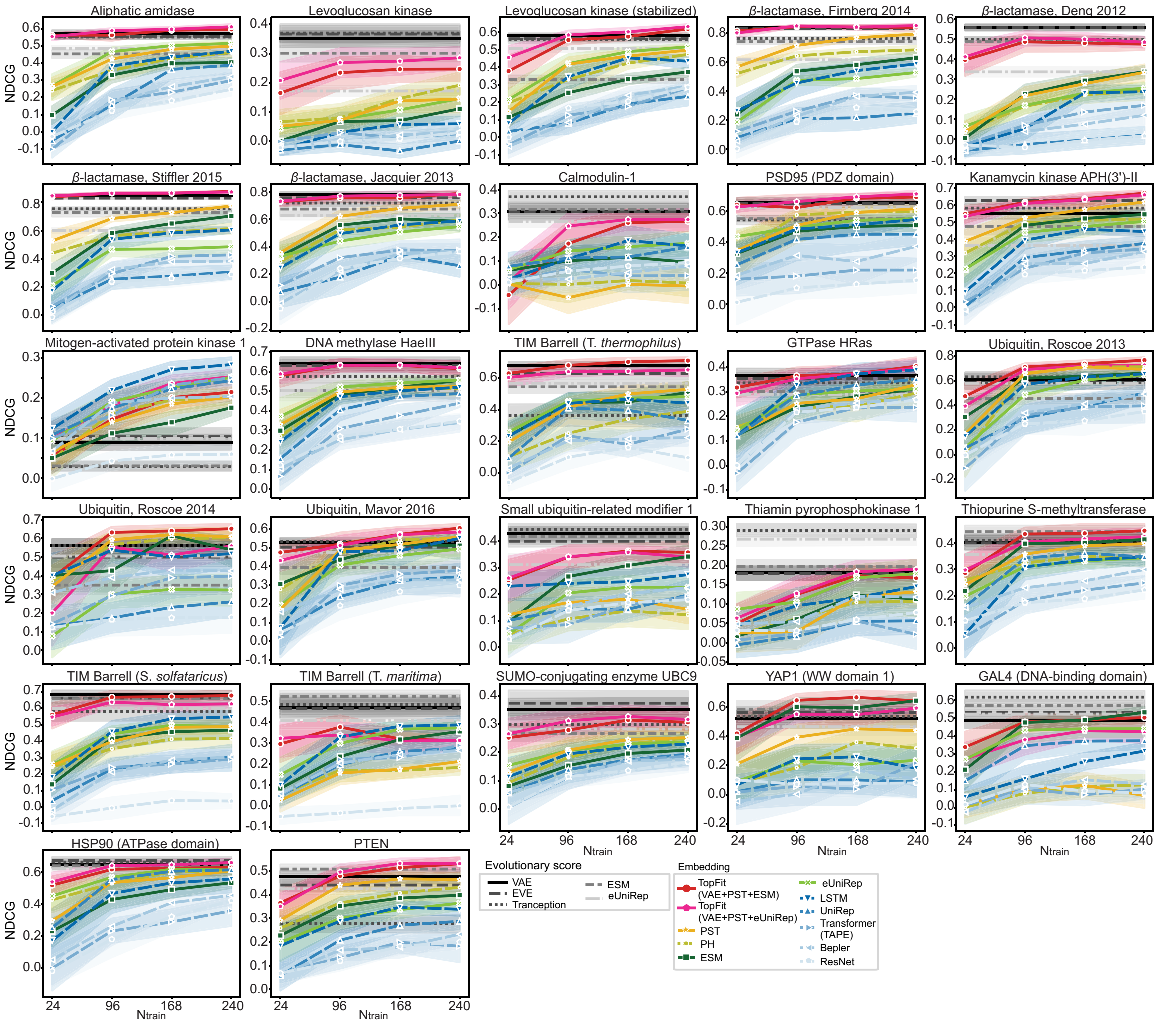

**Supplementary Figure 12: NDCG for various models on individual datasets for predicting unseen mutational sites in training sets.** The ensemble regression is used. The results were shown for training data sizes 24, 96, 168, and 240 from $n = 20$ independent repeats. The width of the shade shows 95% confidence interval. We showed and used absolute values of NDCG for evolutionary scores.

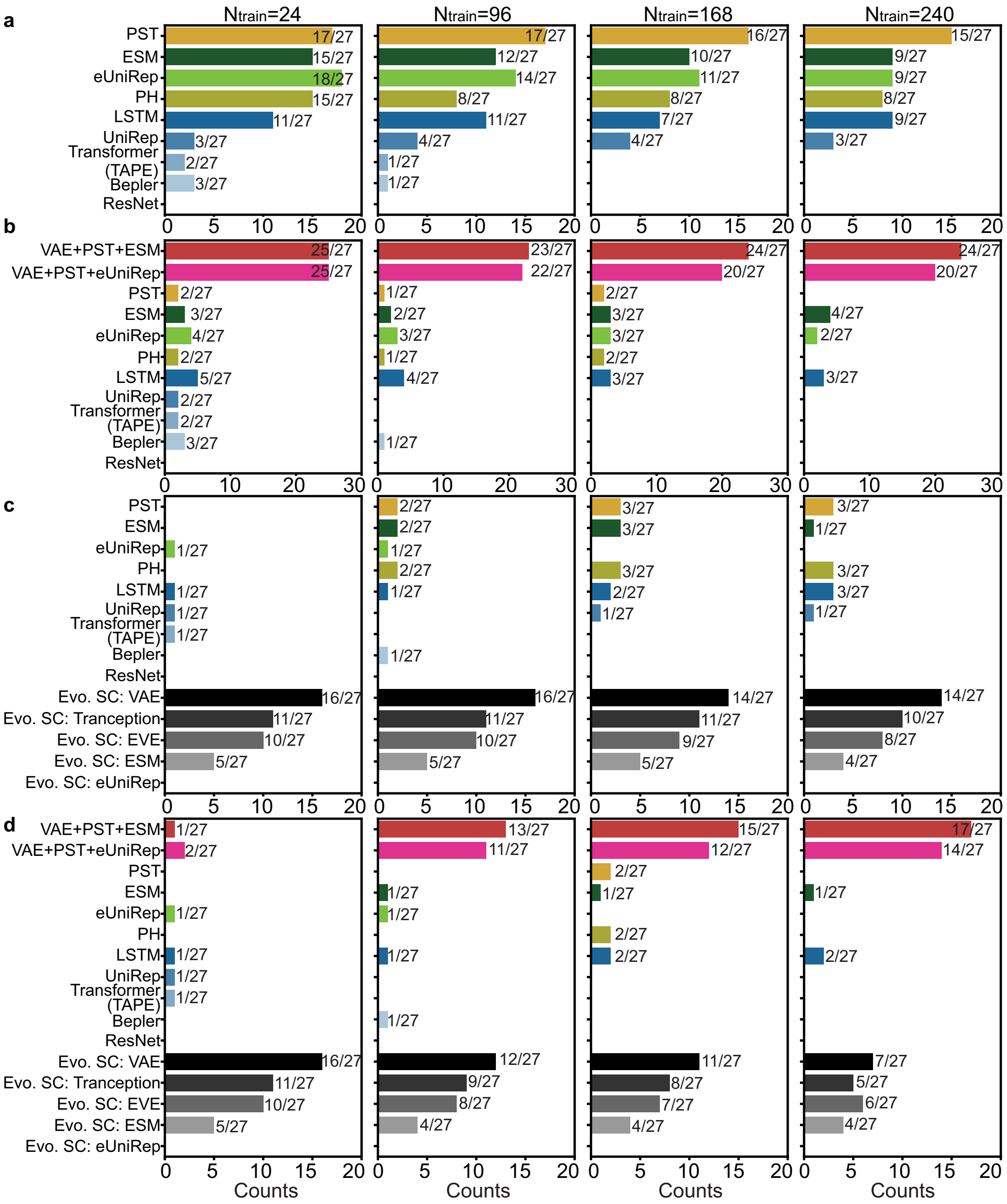


**Supplementary Figure 13: The frequency that an embedding is ranked as the best across 27 single-mutation datasets for predicting unseen mutational sites in training sets, where performance is evaluated by NDCG.** For each dataset, the embedding with the highest average NDCG is identified as the top embedding. The best embedding includes the top embedding and others within the 95% confidence interval of the top embedding. Comparisons for **(a).** structure-based embeddings, sequence-based embeddings, and two sets of TopFit (VAE+PST+ESM and VAE+PST+eUniRep) **(b).** TopFit, structure-based and sequence-based embeddings; **(c).** structure-based embeddings, sequence-based embeddings, and evolutionary scores; **(d).** structure-based embeddings, sequence-based embeddings, evolutionary scores, and two sets of TopFit (VAE+PST+ESM and VAE+PST+eUniRep). We showed and used absolute values NDCG for evolutionary scores. Each embedding has 20 independent repeats.

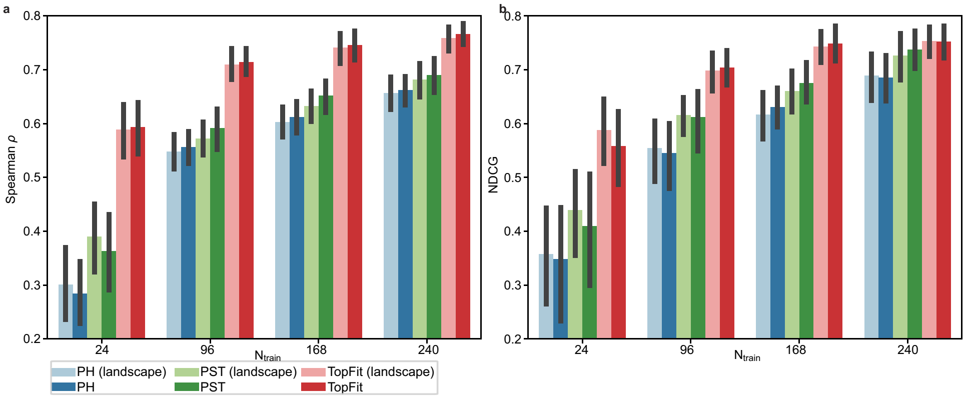


**Supplementary Figure 14: Structure-based embeddings using persistent landscape representations.** PH is persistent homology. PH (landscape) is the persistent homology feature where the vectorization for dimensions 1 and 2 uses the persistent landscape. PST is the persistent spectral theory feature used in the main text. PST (landscape) is the persistent spectral theory feature where the vectorization for dimensions 1 and 2 uses the persistent landscape. TopFit is the feature combining VAE score, PST embedding, and ESM embedding. TopFit (landscape) is the variant of TopFit feature that replaces PST by PST (landscape) feature. The results are calculated on YAP1 (WW domain 1) dataset [1]. The results were shown for training data sizes 24, 96, 168, and 240 from $n = 20$ independent repeats. Bars show average values. Error bars show 95% confidence interval. The ensemble regression is used. **(a).** Spearman correlation and **(b).** NDCG are reported.

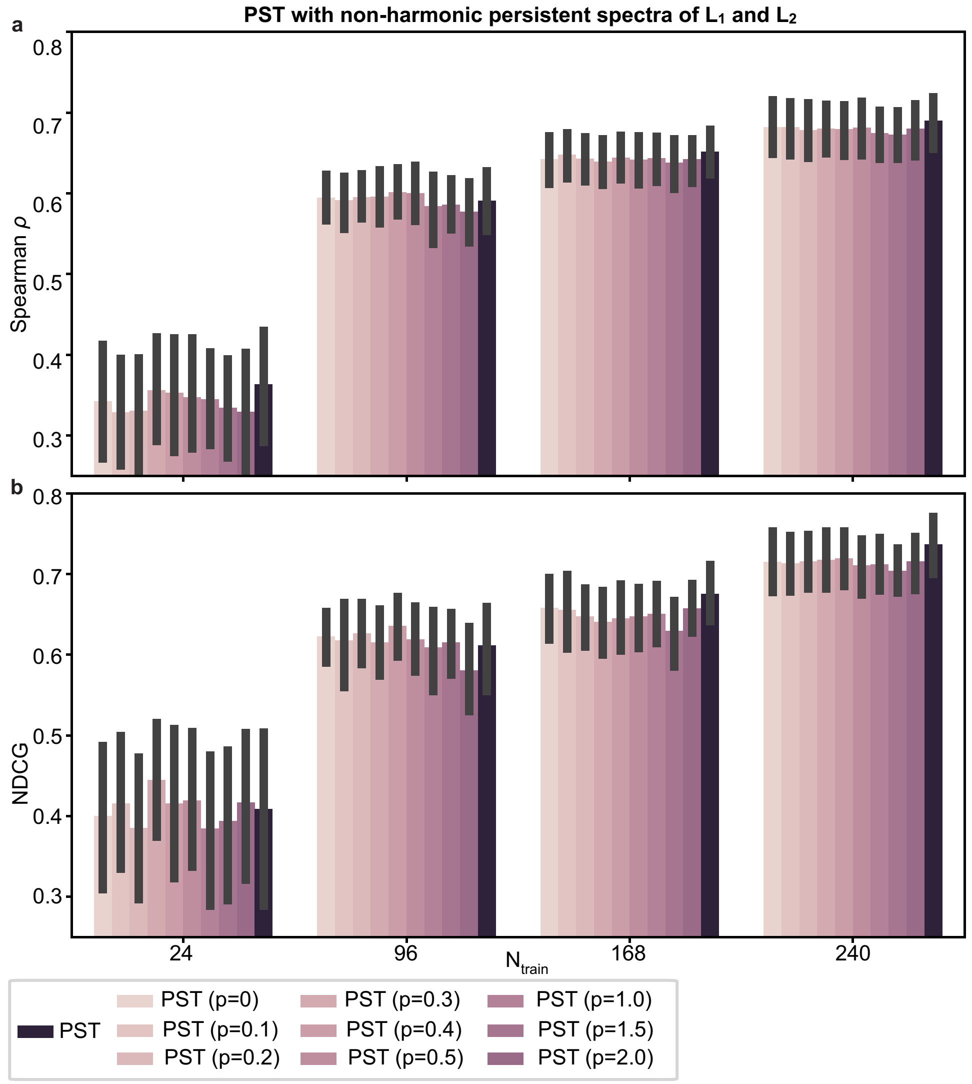


**Supplementary Figure 15: PST embedding with additional non-harmonic persistent spectral features from $L_1$ and $L_2$.** PST is the persistent spectral theory feature used in the main text. PST with $p$ values is the feature with additional non-harmonic persistent spectral features. Here, $p$ is the value of persistence. Specifically, $p = 0$ means no persistence is considered. The results are calculated on YAP1 (WW domain 1) dataset [1]. TThe results were shown for training data sizes 24, 96, 168, and 240 from $n = 20$ independent repeats. Bars show average values. Error bars show 95% confidence interval. The ensemble regression is used. **(a).** Spearman correlation and **(b).** NDCG are reported.

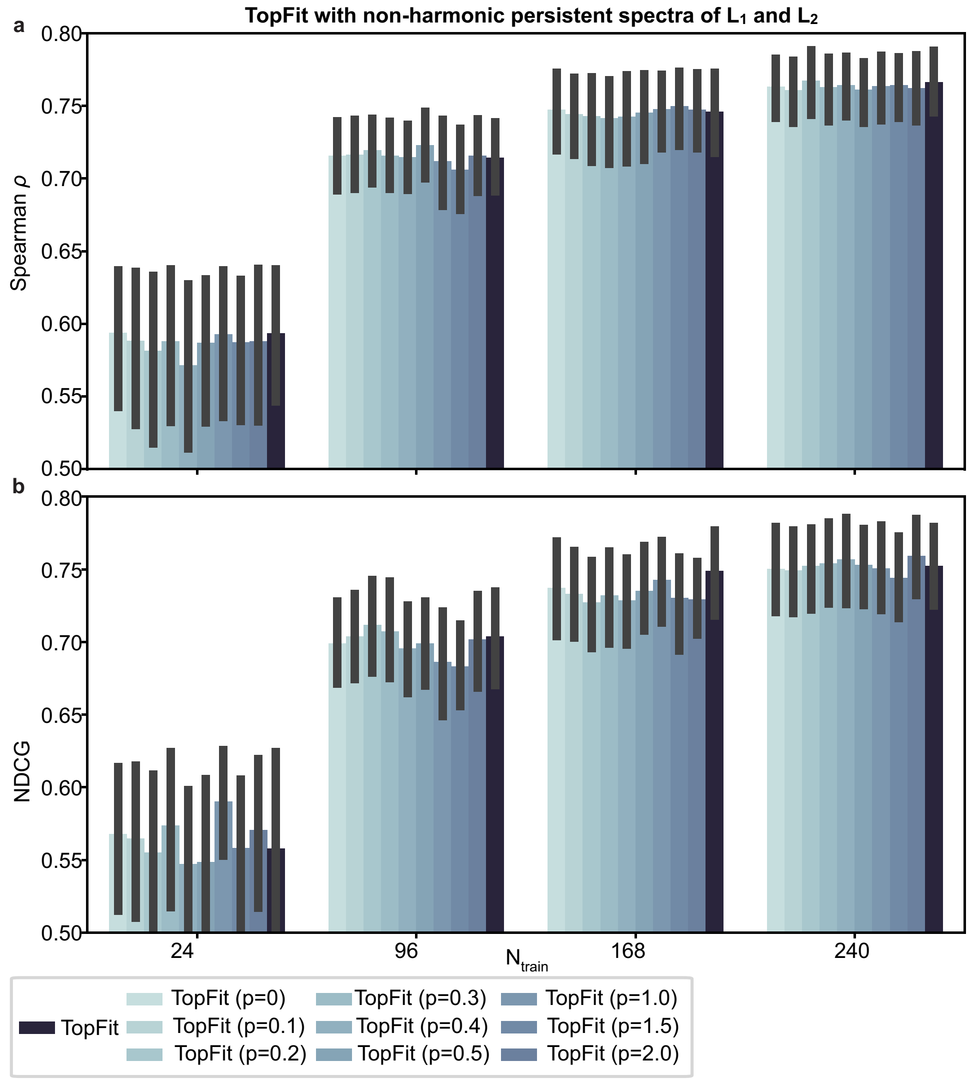


**Supplementary Figure 16: TopFit embedding with additional non-harmonic persistent spectral features from $L_1$ and $L_2$.** TopFit is the feature combining VAE score, PST embedding and ESM embedding used in the main text. TopFit with $p$ values is the feature with additional non-harmonic persistent spectral features. Here, $p$ is the value of persistence. Specifically,, $p = 0$ means no persistence is considered. The results are calculated on YAP1 (WW domain 1) dataset [1]. The results were shown for training data sizes 24, 96, 168, and 240 from $n = 20$ independent repeats. Bars show average values. Error bars show 95% confidence interval. The ensemble regression is used. **(a).** Spearman correlation and **(b).** NDCGs are reported.

| Model type | Packages | Model Name | Rounds of hyper-opt | Small training data? | Large training data? |
|---|---|---|---|---|---|
| Tree | XGB | Tree | 40 | Yes | No |
| Kernel | XGB | Linear | 40 | Yes | No |
| Tree | sklearn-regressor | GradientBoostingRegressor | 0 | Yes | Yes |
| Tree | sklearn-regressor | RandomForestRegressor | 0 | Yes | Yes |
| Kernel | sklearn-regressor | BayesianRidge | 40 | Yes | No |
| Kernel | sklearn-regressor | LinearSVR | 40 | Yes | No |
| Kernel | sklearn-regressor | ARDRegression | 0 | Yes | No |
| Kernel | sklearn-regressor | KernelRidge | 40 | Yes | No |
| Tree | sklearn-regressor | BaggingRegressor | 0 | Yes | Yes |
| Kernel | sklearn-regressor | LassoLarsCV | 40 | Yes | No |
| Tree | sklearn-regressor | DecisionTreeRegressor | 0 | Yes | No |
| Kernel | sklearn-regressor | SGDRegressor | 40 | Yes | No |
| Kernel | sklearn-regressor | KNeighborsRegressor | 0 | Yes | No |
| Kernel | sklearn-regressor | ElasticNet | 40 | Yes | No |
| Kernel | sklearn-regressor | Ridge | 40 | Yes | No |
| ANN | pytorch | OneHidden | 10 | Yes | No |
| ANN | pytorch | TwoHidden | 10 | Yes | No |
| ANN | pytorch | ThreeHidden | 10 | Yes | No |
| ANN | pytorch | OneHidden (wide) | 10 | No | Yes |
| ANN | pytorch | FiveHidden (wide) | 10 | No | Yes |

**Supplementary Table 1: List of regressors used in the ensemble regression.** Hyperparameter optimization was performed on all models except the tree-based method implemented by sklearn-regressor. Numbers of hyperperameters optimization rounds are given in column "Round of hyperopt". Here,"Small training data?" indicates whether the model is used for the fixed and small training data size with $N_{\text{train}} = 24$, 96, 168, and 240. Here, "Large training data?" indicates whether the model is used for 80/20 train/test split or five-fold cross-validation.

| Model | Parameter | Default | Search range | Search method |
|---|---|---|---|---|
| GradientBoostingRegressor | n_estimators | 500 | – | – |
| RandomForestRegressor | n_estimators | 500 | – | – |
| BaggingRegressor | n_estimators | 500 | – | – |
| DecisionTreeRegressor | – | – | – | – |
| Tree (XGB) | eta | 0.3 | $10^{-2} \sim 1$ | loguniform |
| | max_depth | 6 | 3∼8 | quniform; q=1 |
| | lambda | 1 | $10^{-3} \sim 10^{2}$ | loguniform |
| | alpha | 0 | $10^{-3} \sim 10^{2}$ | loguniform |
| | num_boost_round | 500 | 100∼2000 | quniform; q=10 |
| Linear (XGB) | lambda | 1 | $10^{-3} \sim 10^{2}$ | loguniform |
| | alpha | 0 | $10^{-3} \sim 10^{2}$ | loguniform |
| | num_boost_round | 500 | 100∼2000 | quniform; q=10 |
| Ridge | alpha | 1.0 | $10^{-4} \sim 10^{5}$ | loguniform |
| KernelRidge | alpha | 1.0 | $10^{-4} \sim 10^{5}$ | loguniform |
| | kernel | rbf | rbf; laplacian; ploynomial; chi2; sigmoid | choice |
| BayesianRidge | tol | $10^{-3}$ | $10^{-5} \sim 10^{-3}$ | loguniform |
| | alpha_1 | $10^{-6}$ | $10^{-7} \sim 10^{-5}$ | loguniform |
| | alpha_2 | $10^{-6}$ | $10^{-7} \sim 10^{-5}$ | loguniform |
| | lambda_1 | $10^{-6}$ | $10^{-7} \sim 10^{-5}$ | loguniform |
| | lambda_2 | $10^{-6}$ | $10^{-7} \sim 10^{-5}$ | loguniform |
| LinearSVR | tol | $10^{-4}$ | $10^{-5} \sim 10^{-3}$ | loguniform |
| | C | 1 | 0.1∼10 | uniform |
| | dual | True | True; False | choice |
| LassoLarsCV | max_iter | 500 | 10∼1000 | quniform; q=1 |
| | cv | 5 | 2∼10 | quniform; q=1 |
| | max_n_alphas | 1000 | 10∼2000 | quniform; q=1 |
| SGDRegressor | alpha | $10^{-4}$ | $10^{-3} \sim 10^{2}$ | loguniform |
| | l1_ratio | 0.15 | 0∼1 | uniform |
| | tol | $10^{-3}$ | $10^{-5} \sim 10^{-3}$ | loguniform |
| ElasticNet | l1_ratio | 0.5 | 0∼1 | uniform |
| | alpha | 1 | $10^{-3} \sim 10^{2}$ | loguniform |

**Supplementary Table 2: List of hyperparameters and their optimization ranges for kernel and tree models.** If parameters are not specified in "Default", default parameters defined in the software are used. If search range is not given, the default parameter will be used all the time in hyperopt. Search method indicates the sampling method in the given search range. In a given range lower ∼upper, "uniform" uniformly samples within the range, and "loguniform" first maps the range to the log scale and uniformly samples within the mapped range. Here, "quiniform" is a round-off the value from uniformly random sampling in the given range with parameter q: if the sampled value is a, then the round-off value is round(a/q)*q. Here, "choice" returns one of the options in the list given in the search range.

| Model | Parameter | Default | Search range | Search method |
| --- | --- | --- | --- | --- |
| OneHidden; TwoHidden; ThreeHidden | dropout | 0.5 | 0.2∼0.95 | uniform |
| | weight_decay | $10^{-2}$ | $10^{-4} \sim 1$ | loguniform |
| | patience | 20 | – | – |
| | batch_size | 16 | – | – |
| | tol | $10^{-4}$ | – | – |
| | num_epochs | 1000 | – | – |
| | lr | $10^{-3}$ | – | – |
| OneHidden | size1 | 0.1 | – | – |
| TwoHidden | size1 | 0.1 | – | – |
| | size2 | 0.05 | – | – |
| ThreeHidden | size1 | 0.1 | – | – |
| | size2 | 0.05 | – | – |
| | size3 | 0.01 | – | – |
| OneHidden (wide); FiveHidden (wide) | dropout | 0.5 | 0.2∼0.8 | uniform |
| | weight_decay | $10^{-4}$ | $10^{-5} \sim 1$ | loguniform |
| | patience | 20 | – | – |
| | batch_size | 64 | – | – |
| | tol | $10^{-4}$ | – | – |
| | num_epochs | 2000 | – | – |
| | lr | $10^{-3}$ | – | – |
| OneHidden (wide) | size1 | 1 | – | – |
| FiveHidden (wide) | size1,size2,size3,size4,size5 | 1 | – | – |

**Supplementary Table 3: List of hyperparameters and their optimization ranges for ANN models.** If parameters are not specified in "Default", default parameters defined in the software are used. If the search range is not given for the parameter, its default value is used all the time even if the hyperopt is used. The search method indicates the sampling method in the given search range. In a given range lower∼upper, "uniform" uniformly samples within the range, and "loguniform" first maps the range to the log scale and uniformly samples within the mapped range. Here, "quiniform" is a round-off value from uniformly random sampling in the given range with parameter q: if the sampled value is a, then the round-off value is round(a/q)*q. Here, "choice" returns one of the options in the list given in the search range. Here, "size1", "size2", and "size3" are the relative sizes of the first, second, and third layers, respectively. The round-off integer by multiplying the size parameter with the number of features is the number of hidden units at the corresponding layer. If evolutionary scores are used in the features, the number of neurons at the last hidden layer includes an additional hidden unit for the score. The training is stopped when either the epochs reach "num_epochs", or the training error reaches "tol" in patience consecutive epochs. Here, "lr" is the learning rate , and "weight_decay" is the weight decay coefficient used in the Adam optimizer.

| Dataset | 24 | 96 | 168 | 240 | 80/20 split | 5-fold CV |
|---|---|---|---|---|---|---|
| avGFP, Sarkisyan 2016 | 0.607 | 0.71 | 0.732 | 0.746 | – | – |
| GB1, Olson 2014 | 0.478 | 0.682 | 0.756 | 0.809 | – | – |
| Kanamycin kinase APH(3')-II, Melnikov 2014 | 0.553 | 0.661 | 0.687 | 0.701 | 0.789 | 0.791 |
| $\beta$-lactamase, Firnberg 2014 | 0.713 | 0.785 | 0.812 | 0.831 | 0.909 | 0.911 |
| $\beta$-lactamase, Jacquier 2013 | 0.696 | 0.769 | 0.775 | 0.771 | 0.797 | 0.793 |
| $\beta$-lactamase, Deng 2012 | 0.506 | 0.565 | 0.579 | 0.615 | 0.823 | 0.818 |
| $\beta$-lactamase, Stiffler 2015 | 0.771 | 0.812 | 0.834 | 0.844 | 0.908 | 0.909 |
| PSD95 (PDZ domain), McLaughlin 2012 | 0.585 | 0.641 | 0.685 | 0.71 | 0.825 | 0.808 |
| YAP1 (WW domain 1), Araya 2012 | 0.594 | 0.714 | 0.746 | 0.766 | 0.776 | 0.771 |
| Ubiquitin, Roscoe 2013 | 0.443 | 0.654 | 0.73 | 0.762 | 0.839 | 0.848 |
| Ubiquitin, Roscoe 2014 | 0.4 | 0.598 | 0.655 | 0.693 | 0.787 | 0.782 |
| Ubiquitin, Mavor 2016 | 0.425 | 0.581 | 0.639 | 0.662 | 0.738 | 0.751 |
| DNA methylase HaeIII, Rockah-Shmuel 2015 | 0.663 | 0.709 | 0.721 | 0.726 | 0.753 | 0.768 |
| Aliphatic amidase, Wrenbeck 2017 | 0.561 | 0.656 | 0.671 | 0.692 | 0.783 | 0.784 |
| Calmodulin-1, Roth 2017 | 0.083 | 0.132 | 0.193 | 0.221 | 0.289 | 0.289 |
| TIM Barrell (S. $solfataricus$), Chan 2017 | 0.502 | 0.622 | 0.68 | 0.702 | 0.792 | 0.801 |
| Thiamin pyrophosphokinase 1, Roth 2017 | 0.047 | 0.192 | 0.207 | 0.256 | 0.337 | 0.353 |
| GTPase HRas, Matreyek 2017 | 0.41 | 0.58 | 0.642 | 0.686 | 0.844 | 0.845 |
| Small ubiquitin-related modifier 1, Roth 2017 | 0.372 | 0.478 | 0.515 | 0.541 | 0.651 | 0.645 |
| SUMO-conjugating enzyme UBC9, Roth 2017 | 0.369 | 0.553 | 0.585 | 0.62 | 0.759 | 0.753 |
| Levoglucosan kinase, Klesmith 2016 | 0.166 | 0.231 | 0.247 | 0.258 | 0.431 | 0.429 |
| Levoglucosan kinase (stabilized), Klesmith 2015 | 0.423 | 0.498 | 0.529 | 0.555 | 0.731 | 0.723 |
| TIM Barrell (T. $maritima$), Chan 2017 | 0.409 | 0.47 | 0.559 | 0.637 | 0.783 | 0.789 |
| Thiopurine S-methyltransferase, Matreyek 2019 | 0.447 | 0.574 | 0.604 | 0.614 | 0.686 | 0.673 |
| TIM Barrell (T. $thermophilus$), Chan 2017 | 0.543 | 0.669 | 0.728 | 0.771 | 0.852 | 0.861 |
| Mitogen-activated protein kinase 1, Brenan 2016 | 0.163 | 0.248 | 0.293 | 0.321 | 0.558 | 0.554 |
| PTEN, Matreyek 2019 | 0.383 | 0.551 | 0.609 | 0.642 | 0.768 | 0.769 |
| GAL4 (DNA-binding domain), Kitzman 2015 | 0.496 | 0.615 | 0.659 | 0.686 | 0.74 | 0.74 |
| HSP90 (ATPase domain), Mishra 2016 | 0.446 | 0.543 | 0.586 | 0.611 | 0.754 | 0.751 |
| Translation initiation factor IF-1, Kelsic 2016 | 0.449 | 0.59 | 0.665 | 0.708 | – | – |
| BRCA 1 (RING domain) (e3), Findlay 2018 | 0.402 | 0.519 | 0.553 | 0.58 | – | – |
| BRCA 1 (RING domain) (y2h), Findlay 2018 | 0.174 | 0.321 | 0.365 | 0.391 | – | – |
| UBE4B (U-box domain), Starita 2013 | 0.431 | 0.537 | 0.543 | 0.546 | – | – |
| PABP (RRM domain), Melamed 2013 | 0.495 | 0.663 | 0.719 | 0.747 | – | – |

**Supplementary Table 4: Spearman correlation obtained from TopFit by combining PST, ESM embedding, and VAE score.** The model was performed on six types for training data size. Four fixed numbers of training are taken as 24, 96, 168, and 240, and the average Spearman correlation from 20 independent repeats was shown. Either 80/20 train/test or five-fold cross-validation was performed and the average Spearman correlation from 10 independent repeats was shown.

| Dataset | Augmented VAE | | | | | ECNet |
|---|---|---|---|---|---|---|
| | 24 | 96 | 168 | 240 | 80/20 split | 5-fold CV |
| avGFP, Sarkisyan 2016 | 0.663 | 0.682 | 0.703 | 0.725 | – | – |
| GB1, Olson 2014 | 0.491 | 0.651 | 0.714 | 0.759 | – | – |
| Kanamycin kinase APH(3')-II, Melnikov 2014 | 0.61 | 0.621 | 0.636 | 0.653 | 0.737 | 0.754 |
| $\beta$-lactamase, Firnberg 2014 | 0.778 | 0.787 | 0.792 | 0.802 | 0.853 | 0.868 |
| $\beta$-lactamase, Jacquier 2013 | 0.757 | 0.759 | 0.761 | 0.761 | 0.77 | 0.658 |
| $\beta$-lactamase, Deng 2012 | 0.573 | 0.612 | 0.641 | 0.667 | 0.827 | 0.816 |
| $\beta$-lactamase, Stiffler 2015 | 0.783 | 0.793 | 0.802 | 0.814 | 0.875 | 0.868 |
| PSD95 (PDZ domain), McLaughlin 2012 | 0.607 | 0.634 | 0.65 | 0.671 | 0.721 | 0.737 |
| YAP1 (WW domain 1), Araya 2012 | 0.615 | 0.67 | 0.715 | 0.729 | 0.731 | 0.746 |
| Ubiquitin, Roscoe 2013 | 0.492 | 0.653 | 0.69 | 0.714 | 0.771 | 0.834 |
| Ubiquitin, Roscoe 2014 | 0.431 | 0.593 | 0.628 | 0.641 | 0.706 | 0.764 |
| Ubiquitin, Mavor 2016 | 0.472 | 0.565 | 0.596 | 0.627 | 0.69 | 0.711 |
| DNA methylase HaeIII, Rockah-Shmuel 2015 | 0.72 | 0.722 | 0.721 | 0.728 | 0.749 | 0.654 |
| Aliphatic amidase, Wrenbeck 2017 | 0.644 | 0.648 | 0.66 | 0.664 | 0.75 | 0.781 |
| Calmodulin-1, Roth 2017 | 0.231 | 0.234 | 0.231 | 0.227 | 0.296 | 0.268 |
| TIM Barrell (S. $solfataricus$), Chan 2017 | 0.567 | 0.616 | 0.655 | 0.694 | 0.749 | 0.767 |
| Thiamin pyrophosphokinase 1, Roth 2017 | 0.235 | 0.238 | 0.263 | 0.268 | 0.332 | 0.365 |
| GTPase HRas, Matreyek 2017 | 0.488 | 0.554 | 0.598 | 0.64 | 0.781 | 0.828 |
| Small ubiquitin-related modifier 1, Roth 2017 | 0.458 | 0.481 | 0.52 | 0.541 | 0.615 | 0.641 |
| SUMO-conjugating enzyme UBC9, Roth 2017 | 0.503 | 0.577 | 0.622 | 0.65 | 0.737 | 0.724 |
| Levoglucosan kinase, Klesmith 2016 | 0.202 | 0.229 | 0.254 | 0.254 | 0.406 | 0.477 |
| Levoglucosan kinase (stabilized), Klesmith 2015 | 0.451 | 0.502 | 0.508 | 0.509 | 0.626 | 0.72 |
| TIM Barrell (T. *maritima*), Chan 2017 | 0.466 | 0.551 | 0.632 | 0.663 | 0.742 | 0.759 |
| Thiopurine S-methyltransferase, Matreyek 2019 | 0.573 | 0.573 | 0.576 | 0.586 | 0.63 | 0.562 |
| TIM Barrell (T. *thermophilus*), Chan 2017 | 0.606 | 0.683 | 0.722 | 0.746 | 0.807 | 0.816 |
| Mitogen-activated protein kinase 1, Brenan 2016 | 0.175 | 0.241 | 0.244 | 0.275 | 0.485 | 0.426 |
| PTEN, Matreyek 2019 | 0.392 | 0.46 | 0.512 | 0.551 | 0.733 | 0.698 |
| GAL4 (DNA-binding domain), Kitzman 2015 | 0.53 | 0.625 | 0.656 | 0.673 | 0.713 | 0.672 |
| HSP90 (ATPase domain), Mishra 2016 | 0.538 | 0.553 | 0.573 | 0.586 | 0.702 | 0.689 |
| Translation initiation factor IF-1, Kelsic 2016 | 0.514 | 0.548 | 0.578 | 0.602 | – | – |
| BRCA 1 (RING domain) (e3), Findlay 2018 | 0.497 | 0.533 | 0.556 | 0.572 | – | – |
| BRCA 1 (RING domain) (y2h), Findlay 2018 | 0.162 | 0.297 | 0.329 | 0.392 | – | – |
| UBE4B (U-box domain), Starita 2013 | 0.538 | 0.539 | 0.559 | 0.564 | – | – |
| PABP (RRM domain), Melamed 2013 | 0.719 | 0.745 | 0.774 | 0.79 | – | – |

**Supplementary Table 5: Spearman correlation from Augmented VAE model and ECNet model.** The numbers of training data are 24, 96, 168, and 240 with the 80/20 train/test split for the augmented VAE model. The average Spearman from 20 independent repeats was shown. ECNet was performed on five-fold cross-validation. The average Spearman from multiple repeats was obtained from the original work.

### Supplementary Note 1 Embedding comparisons

Here, we categorized embeddings into various classes, and compared their performance within different classes. This is an expansion of Section: Comparing PST embedding with sequence-based embedding .

First, we compared all sequence-based embeddings evaluated by Spearman correlation. Two constant embeddings, i.e., one-hot and Georgiev, provide baselines (Figure 3**a**). In general, Georgiev embedding tends to have better performance with a small number of training data than one-hot embedding while opposite observations can be found with a larger number of training data. Three deep protein language models, i.e., TAPE ResNet, TAPE Transformer, and Bepler embedding, underperform these two constant embeddings. The UniRep embedding achieves better average performance than both constant embeddings, and its fine-tune version (eUniRep) further improves the performance. The eUniRep along with TAPE LSTM and ESM rank at the top in three sequence embeddings, and their performance varies on the number of training data (Figure 3**a**, Extended Data Figure 3). With small training data, TAPE LSTM and eUniRep have similar performance and they have a higher frequency as the best embeddings over all datasets than ESM (Extended Data Figure 3). The eUniRep with fine-tuning on MSAs exhibits partial unsupervised behavior as evolutionary scores, that promises good performance for low-N case. With larger training data, the performance of ESM is improved, and it outperforms TAPE LSTM and eUnirep, especially for the case with 240 training data. Overall, eUniRep and TAPE LSTM models rank as the two best embeddings for the low-N case. As the number of training data increases, the performance of ESM embedding is improved and achieves the best performance when over 168 training data are available (Extended Data Figure 3**a**).

Similarly, for NDCG metrics, ESM embedding ranks most frequently as the best embedding for all numbers of training data, including low-N case. It demonstrates the advantage of ESM over other embeddings in ranking high-fitness variants (Extended Data Figure 5**a**). Unlike Spearman correlation, embeddings ranked by NDCG reveal that two constant embeddings, one-hot and Georgiev, are also top strategies for the low-N case.

Next, we compared the embeddings including structure-based and sequence-based. As discussed in the main text, PST embedding ranks as the best embedding disregarding the large number or small number of training data. This conclusion holds for both Spearman and NDCG metrics (Extended Data Figure 3**b** and Extended Data Figure 5**b**). While for NDCG, the superb performance is more clear where persistent homology (PH) embedding can outperform other sequence-based embeddings, and PST embedding shows further improvement over PH.

Then, we compared all single embeddings and evolutionary scores. As expected, the unsupervised manner of evolutionary scores allows accurate predictions on fitness with no or low-N training data for both Spearman and NDCG metrics (Extended Data Figure 3**c** and Extended Data Figure 5**c**). Since the evolutionary scores provide predictions independent of training data, the regression models built on embeddings tend to surpass evolutionary scores when a sufficient number of training data is available. Particularly, PST embedding shows better performance than evolutionary scores when relatively large training data (i.e., 240) are provided. The model performance changes regarding various training data sizes can be observed in detail at individual datasets for their average Spearman correlation and NDCG (Extended Data Figure 2 and Extended Data Figure 4).

Last, we further included two TopFit embeddings for comparison. The two TopFit models all integrate PST embedding and VAE evolutionary score, and one of them is further aided by ESM embedding and the other is aided by eUniRep embedding. The evolutionary scores consistently show superb performance for the low-N case. However, TopFit embeddings become completely dominant when more training data is available. The two TopFit models all show over 2.5-fold frequently ranked as the best features over other single strategies with over 96 training data provided (Extended Data Figure 3**d** and Extended Data Figure 5**d**). These results further demonstrate the strong generalizability and accuracy in TopFit by combining multiple strategies.

### Supplementary Note 2 Evolutionary scores for unsupervised fitness predictions

In this work, we compared 5 advanced evolutionary scores, DeepSequence VAE [2], EVE [3], Tranception [4], ESM [5], and eUniRep [6], for predicting the relative fitness of protein mutations in an unsupervised manner. More comprehensive comparisons of extensive evolutionary scores were studied previously [7, 8, 4]. All methods tested achieve relatively high Spearman correlations, especially, DeepSequence VAE achieves the best average performance on 34 datasets (Figure 3**a**) and it ranks as the best model on a majority of datasets (21/34).

Overall, evolutionary scores provide the most economic way of predicting protein fitness without labeled data. They generally achieve higher Spearman correlations than supervised models with small training data (e.g., 24 training data). When sufficient training data is available, supervised models tend to surpass unsupervised models (Figure 3**a**, Extended Data Figure 2 and Extended Data Figure 3).

### Supplementary Note 3 TopFit augmented with different evolutionary scores

In the main text, we only combined embeddings with VAE evolutionary score for fitness predictions. Here, we further investigate embeddings combined with other evolutionary scores. When the embedding is augmented with any evolutionary scores, the improvement in Spearman correlation can be seen. Especially, the improvement is significant on poorly performing embedding, for example, TAPE ResNet achieves over 4-fold improvement for the low-N case and around 2-fold improvement for large training data after augmented by any evolutionary scores we tested (Supplementary Figure 6**a-h**). This indicates that the augmentation with an evolutionary score ensures a better lower bound of regression task. Meanwhile, the embedding can further improve the performance with sufficient training data.

Next, we combine PST embedding with either ESM or eUniRep embedding. The combined two embeddings are further augmented with evolutionary scores to predict protein fitness (Supplementary Figure 7). Without the evolutionary score, the combined embeddings alone lead to the lowest average Spearman correlation regardless of the number of training data, and the average Spearman correlation increases as the number of training data increases. With the augmentation of single evolutionary score, the second best score EVE leads to the best performance in TopFit. We further tested the augmentation of multiple evolutionary score in TopFit. All combinations show clear improvement over the single-score augmentation. Interestingly, the two-score combination between VAE and Tranception scores achieves the best average Spearman correlation, and it even outperforms the three-score combination by the inclusion of EVE. Although Tranception is a mixture of local and global evolutionary model, it primarily contains global evolutionary information from large sequence database. The purely local evolutionary model VAE contains evolutionary information from local MSAs complement to Tranception for the best combination in TopFit. While EVE is another local evolutionary information which is redundant in the three-score combination.

In sequence-based feature generation, both evolutionary score and embedding are generated from deep protein language models that learn from millions of protein sequences. The evolutionary score provides a high-level probability distribution for protein mutation in a one-dimensional number, while the embedding has low-level features stored in a high-dimensional vector. Then, the regression model can easily extract the most important information from the evolutionary score, while the embedding needs to be further engineered in a complicated regression architecture. As a result, the augmentation with a one-dimensional evolutionary can largely boost the performance of the regression. Also, the evolutionary score ensures a robust lower bound of predictive performance for the low-N case.

### Supplementary Note 4 Improvements of TopFit

TopFit shows clear improvement over sequence-based methods with the inclusion of structure data (Figure 4 and Figure 5). Such improvement is complicated with integrates featurizaiton and the downstream supervised model. Indeed, multiple factors may contribute to the improvement. In this section, we dissect TopFit improvement in various circumstance.

First, the quality of structure data is critical to structure-based performance, consequently, affects TopFit improvement over sequence-based method (Extended Data Figure 6), as we discussed previously in Section 2.5. Random coils in 3D structure have unstable structures which are difficult to be accurately determined. Indeed, accuracy of structure-based method depends on the percentage of random coils in the target protein (Extended Data Figure 6**a**). B-factor quantifies the atomic displacement in X-ray structures, which can serve as another indicator of structure quality. Indeed, the accuracy of structure-based methods also depends on the B-factor values (Extended Data Figure 6**b**).

By classifying target proteins by their taxonomy into three classes with 11 eukaryote, 12 prokaryote, and 11 human datasets, we looked into the average performance of various models on each class (Supplementary Figure 1). In each class, TopFit shows clear improvement over all single type of embedding regardless of training data size, and it also improves the unsupervised evolutionary scores when sufficient training data is given (i.e. 96 here). In particular, when 240 training data is available, TopFit achieves at least 10.0% improvement on Spearman correlation over sequence-base methods when 240 training data is available. Specifically, TopFit using ESM embedding improves 10.0% and 14.2% over ESM embedding on eukaryote and prokaryote for Spearman correlation, while larger margins are observed for human datasets with 16.6% (Supplementary Figure 1**a-c**). A small improvement (7.5%) over VAE is observed for prokaryote datasets, while TopFit shows significant improvement over VAE on eukaryote and human datasets with 31.5% and 25.1%. The largest improvement from TopFit is observed on human datasets due to the overall lower performance from single strategy on this class of datasets.

Furthermore, the type of fitness measured by experiments may have impacts on supervised model performance and consequently affect the TopFit improvement over sequence-based methods. Indeed, we classify 34 datasets into three classes depending on their fitness experiment: binding, enzyme activity, and others. Fitness for binding includes protein-ligand, protein-DNA and protein-protein binding. Enzyme activity is related to kinase activity and phosphorylation. Rather than these two major fitness types, there are various types of fitness, such as protein fluorescence, protein stability, drug resistance, and growth experiments studied in the benchmark datasets. Since they are only studied in a few datasets, we classify these types of fitness into one single class: others (Supplementary Data 1). In the class of binding fitness, all embeddings including sequence- and structure-based achieve relatively higher performance than other classes (Supplementary Figure 2). Due to the overall good performance of all approaches, TopFit improvement is smallest over other classes. In particular, TopFit using ESM embedding only improves 7.2% in Spearman correlation over ESM embedding when 240 training data is available (Supplementary Figure 2**a**), while it has an extremely large improvement over VAE with margin 48.8%. For the class of enzyme activity, despite the small improvement from TopFit over VAE with margin 8.2% (when $N_{train} = 240$), TopFit improves 29.3% in Spearman correlation over ESM embedding (Supplementary Figure 2**b**). For the class with "others" fitness, all methods achieved lower average performance than other classes. Consequently, TopFit maximize the complementary information from both structure- and sequence-based methods, with 12.7%, 15.4%, and 28.0% improvement over PST embedding, ESM embedding, and VAE scores, respectively (Supplementary Figure 2**c**). Similar behaviors can also be observed in NDCG metrics. The DMS for binding and enzyme activity have standard piplines for experimental assay which may have smaller experimental errors. This may be the reason the average performance of methods tested in these two classes is significantly better than "others" type of fitness. But TopFit consistently shows improvement over sequence-based methods in all classes, which indicates the combination of sequence- and structure-based methods can complement to each other to improve the accuracy in predicting fitness landscape.

### Supplementary Note 5 Predictions of unseen mutational sites

In the main text, we perform train and test sets by random sampling. In protein engineering, it is also important to predict mutations at unseen sites from training set. Indeed, we further tested TopFit and other methods for their ability in generalization for predicting mutations at unseen sites. We performance our benchmark on single-mutation datasets by excluding the datasets with NMR structures, which results in 27 datasets.

To have a split for the training and test sets with different mutational sites, we first split the mutational sites equally into two initial sets, and sample training and test sets within these two sets, respectively. If one of the two initial set contains insufficient number of data for training or testing set sampling, we add an additional mutational site from the other initial set to this initial set. This adjustment will repeat until we have two initial sets containing sufficient number of data for both training and testing sets. Our benchmarks only use fixed sizes of training set with 24, 96, 168, and 240, and test set contains 20% of data . The equal split of two initial sets is usually enough for training and test set splits. The adjustment is occasionally needed for small datasets.

First, we compared various embeddings. From the average performance over 27 datasets, PST embedding outperforms all sequence-based embedding regardless of sizes of training data for both Spearman correlation and NDCG (Supplementary Figure 9), while PST achieves a small margin over the best sequence-based ESM embedding. PST is also ranked most frequently as the best embedding over others evaluated either by Spearman correlation (Supplementary Figure 10, Supplementary Figure 11**a**) or NDCG (Supplementary Figure 12, Supplementary Figure 13**a**).

Next, we compared embeddings with evolutionary scores. All five evolutionary scores show dominant performance over the embeddings with higher average performance in both Spearman correlation and NDCG (Supplementary Figure 9). Only the worst-performing eUniRep score underperforms PST and ESM embedding when sufficient training data is provided. Unlike the random training and testing split (Extended Data Figure 1), the unsupervised evolutionary scores turn to be more powerful to generalize to unseen sites (Supplementary Figure 11**c** and Supplementary Figure 13**c**).

By combining both embeddings and evolutionary score, TopFit has dominant average performance over all embeddings (Supplementary Figure 9). TopFit indeed ranks most frequently as the best methods among all embeddings over 27 datasets evaluated by both Spearman (Supplementary Figure 11**b**) and NDCG (Supplementary Figure 13**b**). However, TopFit can surpass the average performance of evolutionary scores with over 168 training data, and the improvement over evolutionary scores is limited.

Predictions for unseen sites require the models to predict out-of-distribution data. This is extremely difficult for the supervised models of either structure- or sequence-based ones. Our PST embedding consistently outperforms sequence-based embedding for this extrapolation task. However, the unsupervised evolutionary scores are better candidate to handle this type of tasks. On the other hand, in random training and test split, both structure- and sequence-based embeddings can easily surpass evolutionary scores with limited number of training data. TopFit integrate all these approaches together largely enhance its generalizabikity and accuracy in predicting protein fitness landscape. The extrapolation may be needed for the very early stage of protein engineering to screen a few mutations. Due to the much higher computational costs and limited improvement from TopFit than evolutionary scores, evolutionary scores may be more suitable for an initial screen. The supervised TopFit may be more suitable for a later stage with hundreds of available training data.

### Supplementary Note 6 Representations of persistent features

In this work, the element- and site-specific strategies are utilized to generate many sets (i.e., 9 or 10) of point cloud data. PST or persistent homology features indeed have high dimensions even a small dimensional vectorization is employed for each set of point cloud. The high-dimensional features may causes strong overfitting issues in machine learning, especially, for the small training set studied in our work. To accommodate this issue, in topological dimensions 1 and 2, we only calculate seven statistics of Betti-bars to extract the most essential features, which is followed by the previous work in studying protein mutations [9]. This featurization results in a 420-dimensional vector for each mutational entry.

Alternatively, the persistent landscape [10] and the persistent image [11] are two stable representations for persistent homology widely used in machine learning models. Here, we further tested the persistent landscape for dimensions 1 and 2 under our element- and site-specific strategies. We pick the top 3 landscapes with small resolution 10 to obtain the representation, which results in a 30-D vector for each point cloud at topological each dimension. Even with such a small size of representation, a $30 \times 10 \times 2 \times 3 = 1800$-D vector (i.e., 10 sets of point clouds, 2 is due to dimensions 1 and 2, 3 is for the features from wildtype, mutant, and their difference) is generated for each mutational entry. We replaced our featurization for dimensions 1 and 2 by the persistent landscape features in the site- and element-specific settings, and compared with our statistics-based features on YAP1 (WW domain 1) dataset [1]. For PH features, the persistent landscape-type features cause slightly reduction in Spearman correlation when over 96 training data are available (Supplementary Figure 14**a**), while they show slight outperformance on NDCG for several values of training data sizes (Supplementary Figure 14**b**). Similarly, the persistent landscape-based features cause a reduction of performance for both PST and TopFit evaluated by both Spearman correlation and NDCG. Nonetheless, using or not using such persistent landscape-based features, no statistical significant changes can be observed for the model performance (Supplementary Figure 14).

In dimensions 1 and 2, our PST only includes the persistent Betti bars features (i.e., equivalent to harmonic spectra), while the non-harmonic spectra are not considered. Here, we further tested the inclusion of non-harmonic spectra in dimensions 1 and 2. Especially, we tested nine persistence parameters in building the $p$-persistent $q$-combinatorial Laplacian with $p = 0$, 0.1, 0.2, 0.3, 0.4, 0.5, 1.0, 1.5, and 2.0 which are implemented by HERMES [12]. The $p$-persistent Laplacians are calculated at 10 fixed filtration parameters: 2Å, 3Å, $\cdots$, 11Å. Five statistical values of non-harmonic spectra: minimum, maximum, mean, standard deviation, and sum, are used for vectorization. It results in a 50-dimensional vector for each point cloud. With the same strategy using in persistent landscape, this approach results in a 3000-dimensional vector. By adding the non-harmonic spectra to our PST feature used in the main text, we consistently observe the reduced Spearman correlation to our original PST feature (Supplementary Figure 15**a**). While a few cases of inclusion of non-harmonic spectra achieve slightly improved NDCG than our original PST feature when the training set is small with 24 or 96 data, no significant improvement is observed (Supplementary Figure 15**b**). Similarly, the inclusion of non-harmonic spectra in TopFit does not provide any improvement, and reduced average performance is observed more frequently (Supplementary Figure 16).

Enrich information may be obtained from the persistent landscape or non-harmonic spectra at high dimensions. However, the high-dimensional features introduced by these approaches may make the machine learning models more difficult to handle the overfitting and generalization issues. Meanwhile, the high-dimensional features make machine learning models more computational expensive. In our work, since the critical structural information may be mainly contained in topological dimension 0 from both harmonic and non-harmonic spectra, we designed a comprehensive representation for $L_0$ features in TopFit. The high-dimensional feature may provide further improvement, but we use simple and small vectorization of it to overcome overfitting issues.

### Supplementary Note 7 Sources of datasets

The list of data is provided in Supplementary Data 1 with original data sources [13, 14, 15, 16, 1, 17, 18, 19, 20, 21, 22, 23, 24, 25, 26, 27, 28, 29, 30, 31, 32, 33, 34, 35, 36, 37].

[11] Henry Adams, Tegan Emerson, Michael Kirby, Rachel Neville, Chris Peterson, Patrick Shipman, Sofya Chepushtanova, Eric Hanson, Francis Motta, and Lori Ziegelmeier. Persistence images: A stable vector representation of persistent homology. *Journal of Machine Learning Research*, 18, 2017.
